# Response to geographic variation in song is associated with differential gene expression in the blood of a songbird

**DOI:** 10.64898/2026.05.20.726641

**Authors:** Gabriel Macedo, Brianna McKenna, Susan Peters, Stephen Nowicki, Sara Lipshutz

## Abstract

Birdsong mediates territory acquisition and mate choice. In agonistic interactions, local songs generally elicit stronger responses than songs from more distant populations. However, the molecular mechanisms associated with differential responses to local vs. foreign songs are poorly understood. We addressed this knowledge gap by combining behavioral assays in the field with blood transcriptomic analysis, using a within-subjects design to ask whether male song sparrows (*Melospiza melodia*) show differential gene expression when exposed to playback of local and foreign songs. Transcriptomic profiles reflected the difference in behavioral response to local vs. foreign songs, with individuals exposed to local songs showing greater expression of genes associated with song perception and production, anti-inflammatory responses and energy metabolism. Our study suggests that changes in expression of key molecular pathways correlate with behavioral responses to geographic song variation, providing insight into the potential mechanisms regulating signal recognition and response to social challenges.

**Highlights:**

- Gene expression in sparrow blood was measured after simulated territorial intrusion.
- Stronger response to local songs was associated with differential gene expression.
- Song-associated genes (*FOXP2, NRXN1*) had higher expression when birds heard local songs.
- Gene expression in the blood contains potential biomarkers of song recognition.

## Introduction

Birdsong, like social signals across many taxa, plays a key role in important life history events, including territory defense and mate attraction^1,2^. Studies have long documented that both territorial responses of males and copulation solicitations of females are stronger towards songs from their local population than geographically distant populations^3–6^. These geographic patterns of response indicate that local, conspecific songs are more salient signals in sociosexual contexts than foreign songs, which has broad implications for understanding population divergence and speciation^7–9^. While a robust body of research exists on many ecological and evolutionary aspects of signal-mediated social challenges, the molecular mechanisms associated with behavioral responses to social signals are poorly known (Louder et al., 2018, 2020).

Behavioral responses during social interactions can elicit rapid changes (i.e., within minutes to hours) of gene expression in the brain^10–12^. Neurotranscriptomic responses to social challenges are similar across diverse taxa such as sticklebacks, mice and honeybees, suggesting these responses may be evolutionarily conserved^10–13^. Obtaining brain tissue to measure gene expression levels requires lethal collection of specimens, however, which limits study designs to a single condition of interest per specimen^15^ and prohibits the comparison of behavioral responses to different stimuli in the same individual.

Sampling gene expression in the blood may present an alternative to lethal collection, in cases where extracellular mRNA reaches the bloodstream^15,16^, or similar genes are expressed in both brain and blood^17–19^. The connection in gene expression between blood, brain and other tissues can facilitate non-lethal sampling of individuals across different experimental conditions^14^, and through time^20,21^, as well as studies of endangered taxa^22–24^. An experiment with zebra finches (*Taeniopygia guttata*) comparing gene expression in response to conspecific versus heterospecific songs found a positive correlation between gene expression in the auditory cortex and blood sampled from peripheral tissues^17^. Thus, gene expression in the blood may be informative for understanding physiological responses to different, ecologically relevant stimuli. For instance, a study with wild red-winged blackbirds (*Agelaius phoeniceus*) found that calls from conspecific individuals and from a heterospecific nest parasite (brown-headed cowbird; *Molothrus ater*) were associated with down-regulated expression of genes associated with immune response in the blood relative to control calls of a harmless, non-parasitic heterospecific (mourning dove; *Zenaida macroura*)^14^. In contrast, only calls from conspecifics induced in the blood upregulation of metabolism-associated genes relative to the two heterospecific calls^14^. However, it is currently not known whether gene expression differs between individuals exposed to local vs. foreign signals within the same species, leaving a gap in our knowledge on the molecular correlates of responses to social challenges involving conspecific signals.

The song sparrow is a widely distributed songbird in North America^25^ that has long been a model in studies of song development, recognition, and geographic variation^3,6,8,26–28^. Individual male song sparrows sing a complex repertoire of five to 16 song types, with different song types typically composed of several distinct phrases and each phrase including a number of different note types^8,26,27^. Behavioral tests demonstrate that both males and females recognize differences in acoustic structure associated with geographic variation in song, responding significantly less to songs from more distant populations^3,6,8,26^. An early molecular study using in-situ hybridization found that neuronal expression levels of *ZENK*, an immediate early gene associated with auditory responses, increased when song sparrows were exposed to conspecific songs relative to silent controls^11^. This extensive foundational work on song sparrow response to song provides an excellent opportunity to investigate gene expression in relation to conspecific song recognition.

Here, we use behavioral experiments and transcriptomic analyses to test whether gene expression in the blood reflects geographic discrimination of conspecific social challenges in free-living male song sparrows. We conducted paired, within-subjects simulated territorial intrusions to evaluate the impact of local vs. foreign song treatments on gene expression in the blood. Furthermore, we contrasted the effect of a social challenge (local and foreign songs) with the absence of social challenge by sampling individuals that were not subject to any behavioral trial. We expected that, relative to foreign songs, local songs would elicit stronger aggressive responses and differential expression in the blood^14^. Consistent with earlier studies, we found that individuals responded more weakly to foreign songs than to local songs. Furthermore, gene expression profiles of individuals exposed to foreign songs were similar to individuals that were not subjected to any social challenge. In contrast, we found that stronger territorial responses to local songs were associated with up-regulated expression of genes involved in song perception and production pathways (e.g., *FOXP2, NRXN1, GABBR2*), as well as immune response and energy metabolism. Our findings suggest that the blood of a wild bird contains relevant biomarkers for discrimination of intraspecific song variation, providing insights into the potential molecular mechanisms of the response to social challenges.

## Results

### Behavioral responses are stronger toward local songs and during first trials

We tested 15 individuals in within-subjects paired behavioral trials. Seven subjects heard local songs in their first trial and then foreign songs in a second trial. Eight subjects heard foreign songs in their first trial, but one of these individuals disappeared from its territory before being tested again, with the remaining 7 individuals given a second trial in which they heard local songs (see STAR Methods). We found that individuals responded more strongly (i.e., approached more closely on average during trials) to local songs than to foreign songs (Figure 1a; β = 1.08 [0.04: 2.14], P = 0.04; henceforth, numbers in square brackets are 95% credible intervals). We also observed an effect of trial order: individuals responded more strongly in their first trial than in their second trial (Figure 1b; β = 1.17 [0.09: 2.21]; P = 0.03). We found no evidence for an interaction between treatment and trial order (β = 0.11 [-2.00: 2.32]; P = 0.94). We found qualitatively similar results with a non-parametric model, with significant effects of treatment (P = 0.03) and trial order (P < 0.001).

**Figure 1.**
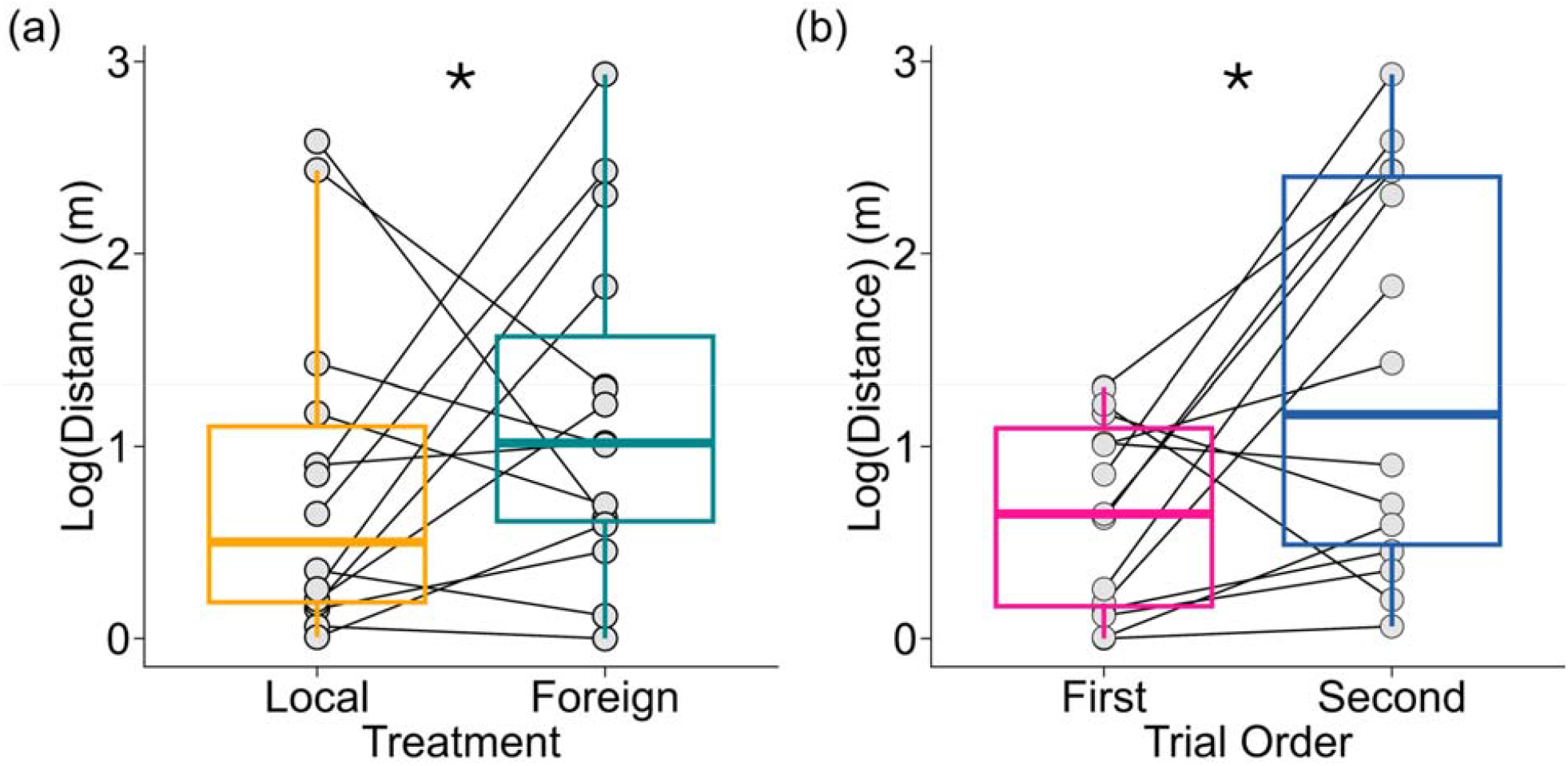
Male song sparrows approached more closely (i.e. smaller distance) during trials with (a) local songs than in trials with foreign songs, and (b) in their first trial than in their second trial. The sample sizes are 15 individuals in the first trial (N = 7 local song treatment, N = 8 foreign song treatment) and 14 individuals in the second trial (N = 7 local song treatment, N = 7 foreign song treatment). Boxplots are in the style of Tukey and lines connect responses of the same individuals. Circles represent individual birds. Asterisks indicate that comparisons are credibly different from zero.

### Expression profiles in the local song treatment differed from the foreign song treatment and the baseline

To understand the effects of social challenge vs. the absence of a social challenge on gene expression, we investigated whether expression profiles differed between individuals tested with local or foreign songs in their first trials and individuals that were not subjected to any behavioral trial (baseline control). Sample sizes were 10 baseline individuals, 8 individuals in the foreign song treatment, 7 individuals in the local song treatment. We found that expression profiles of baseline individuals were more similar to individuals in the foreign song treatment than to individuals in the local song treatment (Figure 2). We detected 21 differentially expressed genes (DEGs; 16 up-regulated, 4 down-regulated) between local and foreign treatments, 38 DEGs (27 up-regulated, 11 down-regulated) between the local treatment and the baseline, and 5 DEGs (4 up-regulated, 1 down-regulated) between the foreign treatment and the baseline (Tables S1-3). Contrasts with the local song treatment (local vs. foreign, local vs. baseline) shared 15% (8 genes) of the DEGs, whereas contrasts with the foreign song treatment (foreign vs. local, foreign vs. baseline) shared 4% (2 genes) of the DEGs, and no DEG was common to all contrasts (Figure 2). Through gene set enrichment analysis, we found that up-regulated DEGs in the contrast between local and foreign songs are associated with neuronal and cognitive development, vocal and auditory learning, and social behavior (e.g., *FOXP2, GABBR2, NRXN1, CNTNAP2, GLI3*; Supplementary Material Table S1), and down-regulated DEGs are associated with immune and spermatic functions (e.g., *ZPBP2*; Table S2). Conversely, up-regulated DEGs in the contrast between local song and baseline were associated with presynaptic membrane organization (e.g., *NRXN1, IL1RAPL1*) and phosphatase activity (e.g., *PFKFB3, PPP1R14C*; Table S3), and down-regulated DEGs were associated with DNA damage sensors and tumor suppression (e.g., *BRCA1-CtIP-ZBRK1* repressor complex; Table S4). Down-regulated DEGs in the contrast between foreign song and baseline were associated with protein and lipid metabolism (*DBT*; Table S5), whereas up-regulated DEGs were not associated with any ontology terms.

**Figure 2.**
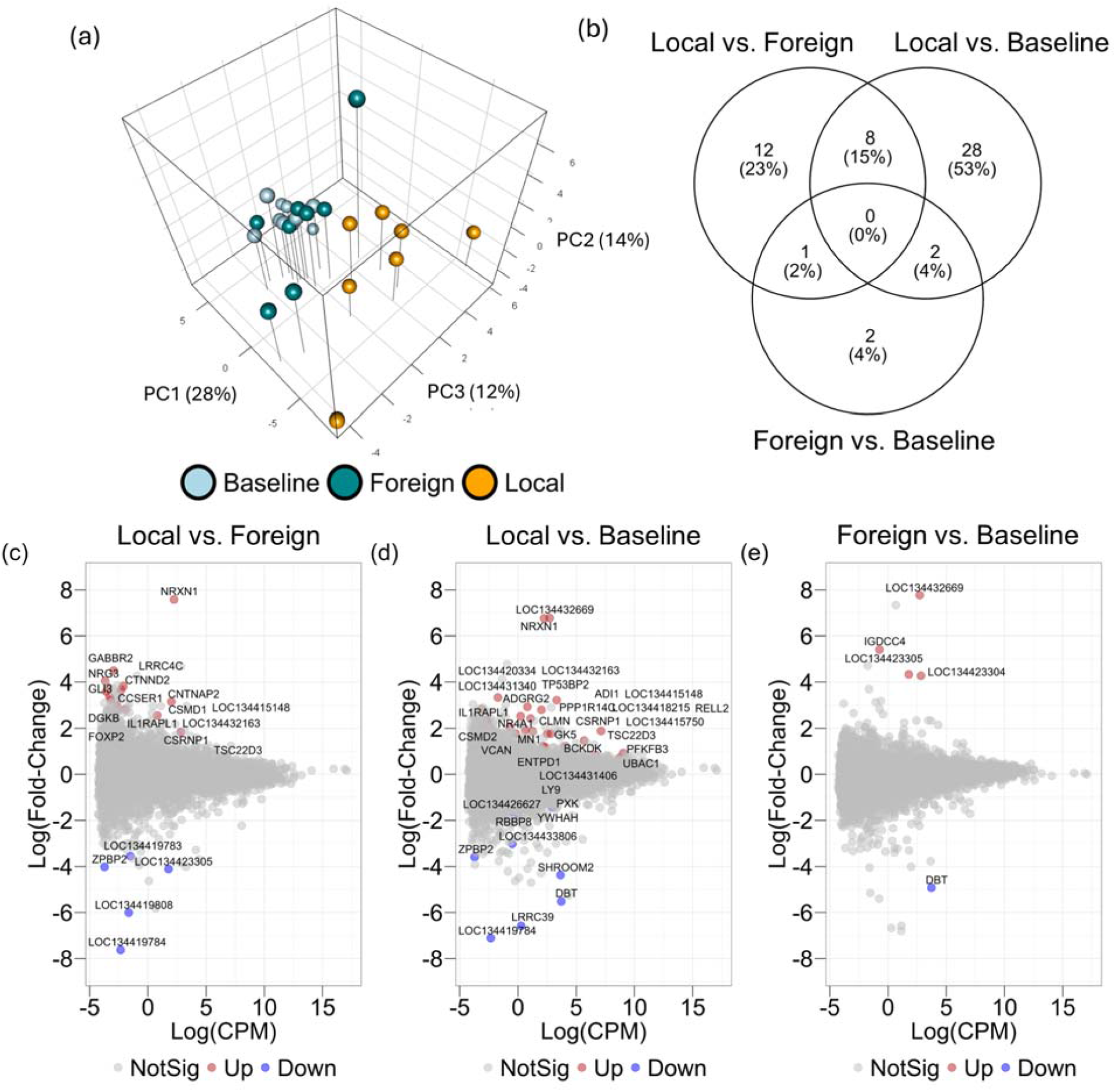
In first trials, expression in the local song treatment differs from both the foreign song treatment and the baseline, whereas fewer differentially expressed genes (DEGs) are present in the contrast between the foreign song treatment and the baseline. (a) PCA plot of all DEGs across treatments. Spheres represent individual birds in each treatment. Sample sizes are 10 baseline individuals, 8 individuals in the foreign song treatment, and 7 individuals in the local song treatment. (b) Venn diagram showing the number and percentage of DEGs in the contrasts between treatments. (c-e) Mean-difference plots between treatments showing log fold-change versus log counts per million. Each point represents a gene; genes in grey and not differentially expressed, genes in red are significantly up-regulated and genes in blue are significantly down-regulated.

To examine the collective action of genes across the transcriptome, we used Weighted Gene Co-expression Network Analysis (WGCNA). One gene module was significantly associated with treatment (turquoise; r = 0.53; P = 0.006). Module expression in the local song treatment differed from both the baseline and the foreign song treatment (Figure 3a). In contrast, module expression was similar between the foreign song treatment and baseline. Genes in the turquoise module are transcription factors, regulation of RNA synthesis and metabolism, social behavior, speech development, and diverse somatic developmental abnormalities (*DOT1L, ACVR2A, NSD1, NCOR1, MED13, CREBBP*; Table S6). We did not find any gene modules associated with time to capture (time in minutes from the start of the capture playback until the individual was retrieved from the mist net).

**Figure 3.**
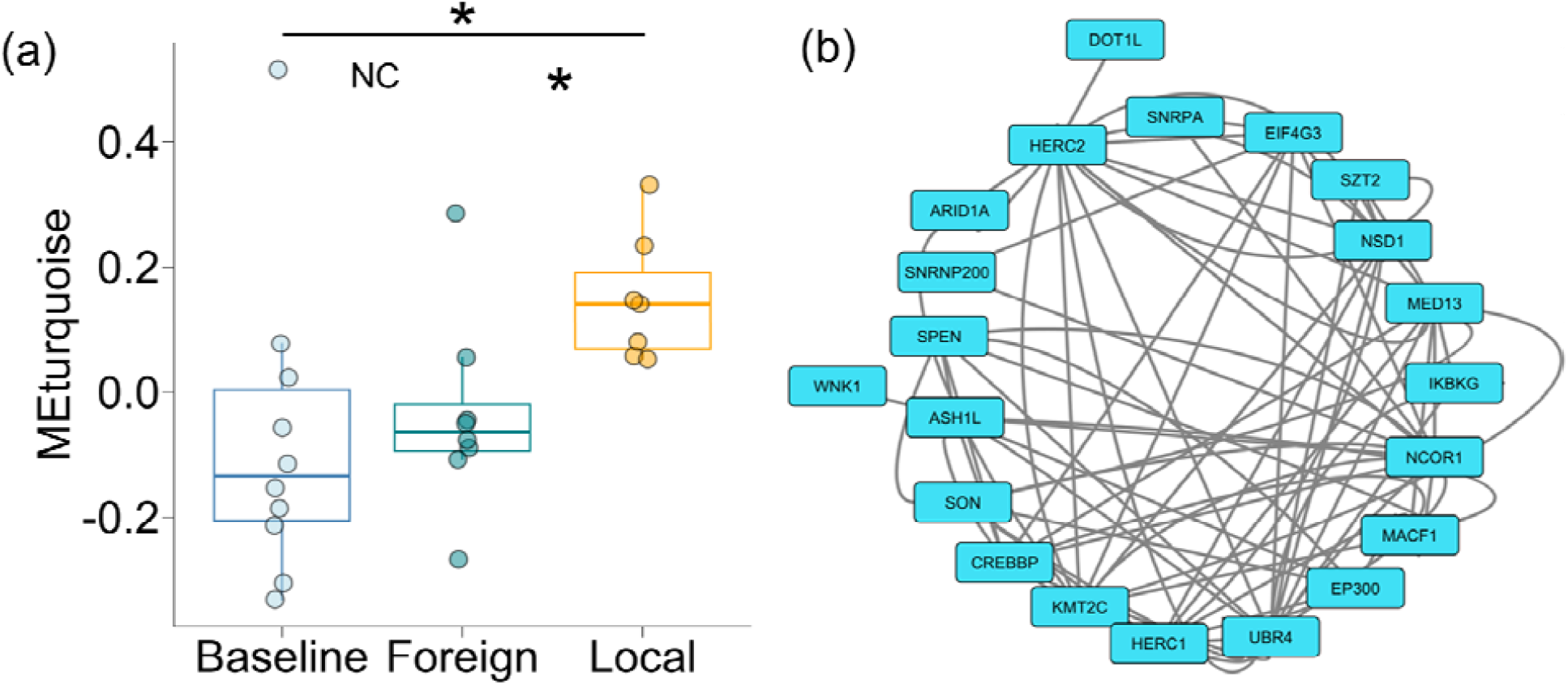
The turquoise gene network is significantly associated with treatment. (a) WGCNA eigengene values of the turquoise network show differential expression in the local song treatment in relation to the foreign song treatment and the baseline. Sample sizes are 10 baseline individuals, 8 individuals in the foreign song treatment, and 7 individuals in the local song treatment. Boxplots are in the style of Tukey and circles represent individual birds in each treatment. Asterisks indicate contrasts that are credibly different from zero, and NC, not credibly different from zero. (b) Turquoise network diagram depicting genes with network membership >|0.6|.

### Treatment but not trial order affected gene expression in paired trials

We analyzed paired trials to estimate the effects of both treatment and trial order on gene expression (N = 8). Baseline individuals do not have a trial order and were excluded from this analysis. Song treatment, but not trial order, affected gene expression (Figure 4). We found 14 up-regulated DEGs in the comparison between local and foreign songs, whereas there was none associated with trial order. These DEGs are associated with cell adhesion (e.g., *CDH18, PCDH7, CADM2*; Table S7). With WGCNA, we found one gene module associated with treatment, darkorange2 (r = 0.54; P = 0.03), but none associated with trial order nor time to capture (Figure 5; Figure S2). The genes in the darkorange2 module are associated with cell junctions, nervous system development, and autistic behavior. Some of the genes in this module were the same DEGs detected when comparing the first trials and the baseline (*FOXP2, GABBR2, NRXN1, CNTNAP2*; Table S8).

**Figure 4.**
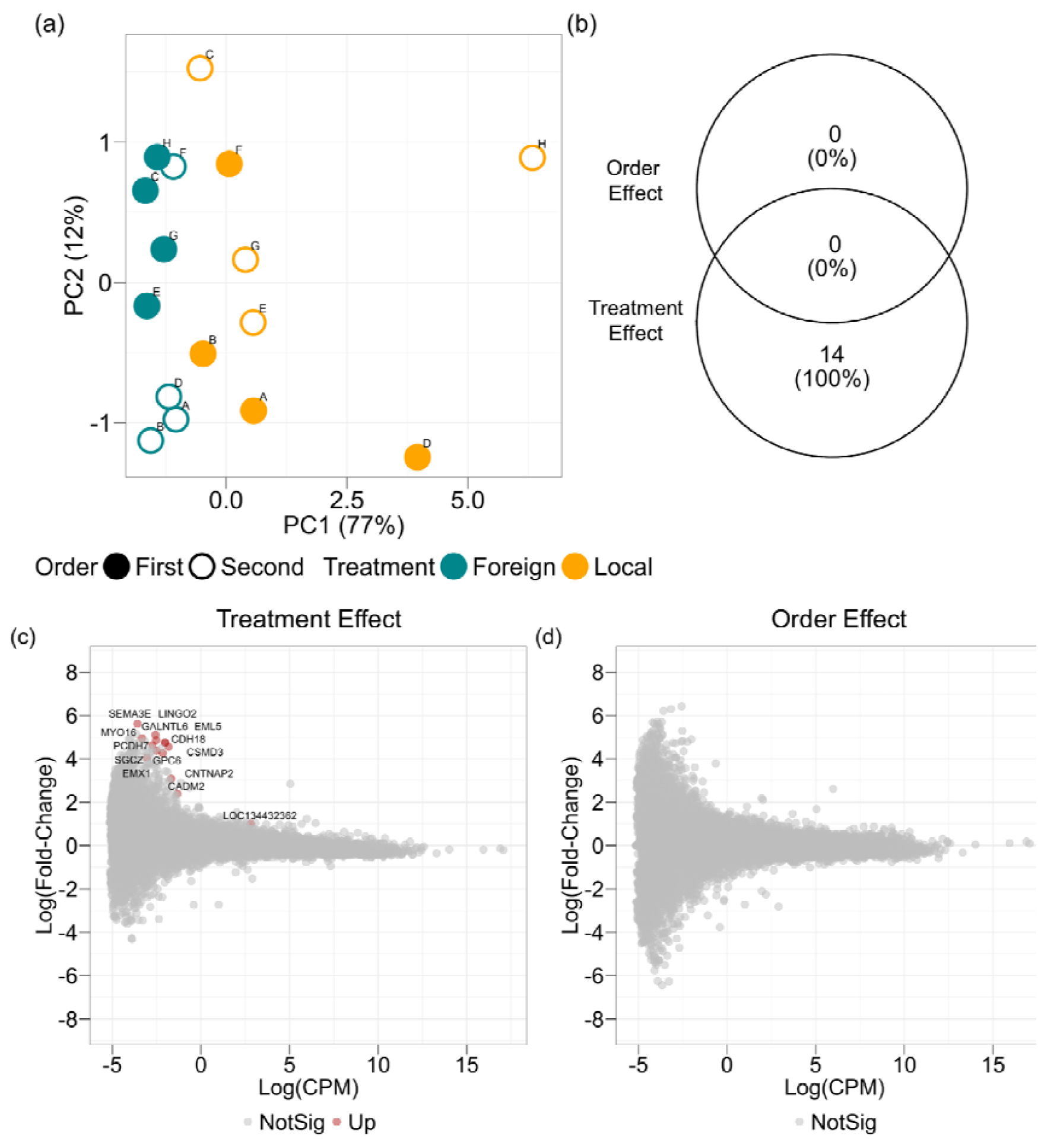
In paired trials, expression in the local song treatment differs from the foreign song treatment, but not in relation to trial order. (a) PCA plot of all DEGs between treatments and trial order; circles represent individual birds in each treatment, filled circles represent individuals in their first trial, and open circles represent individuals in their second trial; letters indicate individual identification. The sample size is 8 individuals, each tested twice with local and foreign songs in the first or second trial. (b) Venn diagram showing the number and percentage of DEGs in the effect of treatments and the effect of trial order. (c-d) Mean-difference plots of the treatment and order effects showing log fold-change versus log counts per million. Each point represents a gene; genes in grey are not differentially expressed and genes in red are significantly up-regulated.

**Figure 5.**
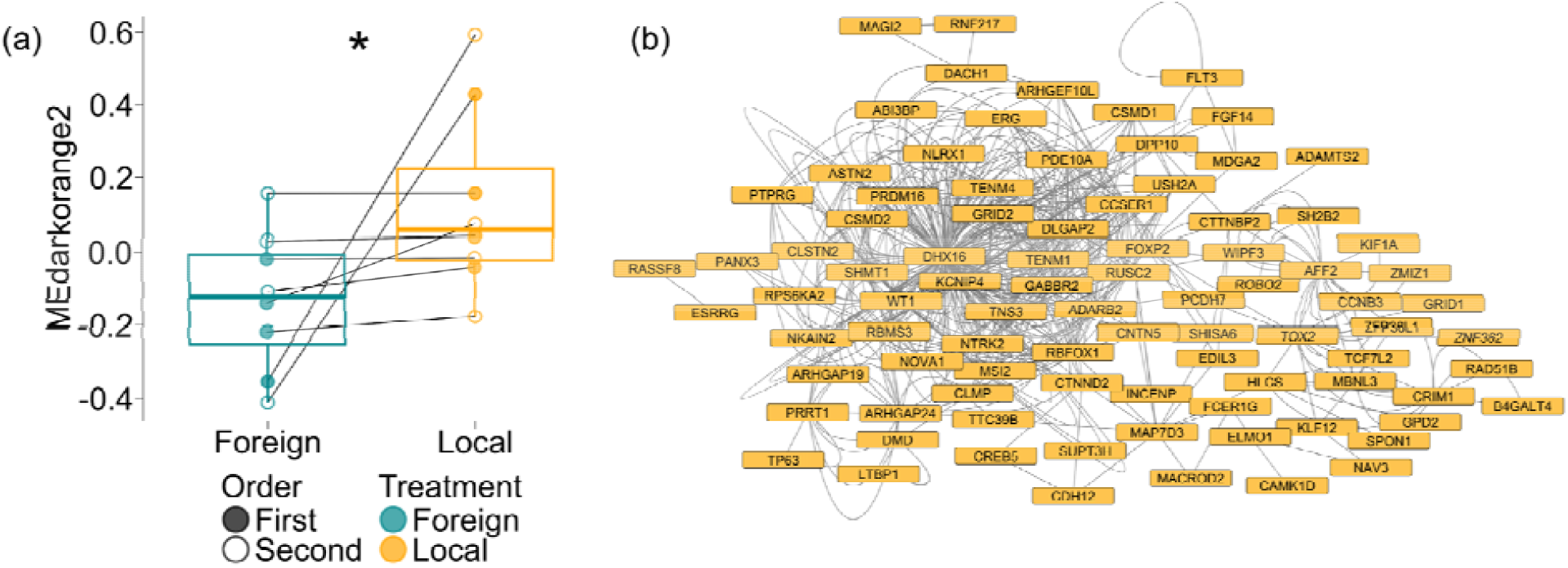
The darkorange2 gene network is significantly associated with treatment, but not trial order. (a) WGCNA eigengene values of the darkorange2 network show differential expression in the local song treatment in relation to the foreign song treatment. The asterisk indicates that the contrast is credibly different from zero. The sample size is 8 individuals, each tested twice with local and foreign songs in the first or second trial. Boxplots are in the style of Tukey and circles represent individual birds in each treatment, filled circles represent individuals in their first trial, and open circles represent individuals in their second trial. Lines connect the same individuals in the first and second trials. (b) Darkorange2 network diagram depicting genes with network membership >|0.6|.

## Discussion

Using a within-subjects design, we replicated earlier findings that song sparrows show stronger territorial responses to local songs than to foreign songs^3,6,8^. Notably, we found that exposure to these different song treatments are associated with differential gene expression in the blood. Individuals exposed to foreign songs show more similar expression profiles to baseline individuals that were not subjected to any social challenge, as compared to individuals exposed to local songs. Additionally, we found that birds showed stronger behavioral responses in their first trials than in second trials, whereas gene expression was only explained by song treatment, with no effect of trial order. Taken together, our results suggest that recognition of subtle geographical song variation elicits fast transcriptional changes (i.e., in less than 1.5 hours) that are detectable in the blood of song sparrows. These findings have implications for our understanding of the molecular correlates of signal of recognition and population divergence of signals, as well as the feasibility of blood sampling as a less-invasive, non-lethal method of detection of biological markers of signal recognition.

### Behavioral responses and gene expression in the first trials

We found an effect of song treatment and trial order on the behavioral responses, with individuals approaching more when exposed to local songs and in their first trials in relation to foreign songs and second trials. The detection in the local song treatment of up-regulated genes typically associated with vocal and auditory learning and social behavior is intriguing (e.g., *FOXP2, GABBR2, NRXN1, CNTNAP2*), given that these gene functions have been described in brain nuclei of birds, mammals, and other animals associated with vocal communication^29,30^. However, in peripheral tissues of adult individuals these genes are also associated with physiological functions beyond those associated with neural processing of songs. For instance, the expression of *FOXP2* in in peripheral tissues of adult rats and humans is also associated with inflammatory response and cancer^31,32^, *GABBR2* is associated with angiogenesis in body parts such as limbs^33^, and *NRXN1* is also expressed in leucocytes in the peripheral blood of humans with Autism Spectrum Disorder^34^. Therefore, these “song-associated” genes may reflect other aspects such as greater stress levels, physical movement, and metabolic demands induced by local songs during the trials, and not necessarily vocal behavior or auditory response. There is one line of evidence, however, that may support an explanation associated with song discrimination for these specific gene expression patterns. We conducted the experiments in early spring during the reproductive season, when songbirds undergo neuroplastic seasonal changes in the brain, with the growth of song-related nuclei and expression of genes associated with increased vocal and territorial activities^35,36^. Because gene expression in the brain and peripheral blood have been shown to be correlated^17,19,37^, it may be possible that the expression of genes in the brain associated with song hearing is detectable in the blood. Unfortunately, our study was not designed to measure this association (see Limitations below).

Trials with local songs elicited gene expression profiles that were more distinguishable (i.e., with more DEGs and network eigengenes) from trials with foreign songs and from baseline individuals than the comparison between trials with foreign songs and baseline individuals. This finding is consistent with the interpretation that local songs represent a more salient signal that elicits greater changes in gene expression. Additionally, the contrasts of the song treatments with baseline individuals reduces concerns about false positives that may occur in transcriptomic studies lacking control groups^38^. In comparison to the baseline, the local song treatment elicited DEGs associated with sensing of DNA damage and tumor suppression (e.g., *BRCA1-CtIP-ZBRK1* repressor complex, which also function in oxidative stress response^37,39^), whereas foreign songs elicited greater expression of genes associated with protein and lipid metabolism (e.g., *IGDCC4, DBT*). In relation to baseline expression levels, these DEGs may reflect the stress and physical exertion associated with responding to a social challenge.

### Within-subjects designs and effect of trial order

Within-subject paired designs help to control for individual variation while (ideally) varying only the treatment of interest^40^. In the present study, however, because we needed to capture the same individual to sample blood after their local and foreign song trials, capture presented an additional disturbance not related to the content of experimental stimuli. During the first trials, the birds were naïve to our experimental setup (presence of the experimenters, equipment, capturing, blood sampling), whereas in the second trials, their weaker behavioral responses and evasive maneuvers to avoid being recaptured likely indicate they were aware of the experimenters and/or our setup. Nonetheless, we found that birds exposed to local songs show differential gene expression in relation to birds exposed to foreign songs, regardless of trial order. Previous within-subjects behavioral studies with song sparrows did not find an order effect, likely because the birds were not captured, marked, and bled after a first trial, and only captured and marked after completing all behavioral assays^6,8^. Additionally, these previous studies used experimental stimuli with shorter durations (6 min)^6,8^ than the present study (30 min), which may also explain the presence of the effect of trial order observed here.

Despite the order effect on behavioral responses, when comparing expression levels of the individuals sampled both in the first and second trials (paired trials), we did not detect any gene modules associated with trial order. In contrast, we did detect DEGs and a gene module that differed between local and foreign song treatments. In our analysis focusing on individual DEGs, differential expression was mainly associated with genes involved in neuronal cell adhesion (e.g., *CDH18, PCDH7, CADM2*), although these genes also play roles in heart muscle function, tumor suppression, and energy homeostasis in mice and humans^41–43^. Through WGCNA, we found that the gene network associated with song treatment in the paired trials contained song-associated genes in common with the analysis of only the first trials and the baseline (e.g., *FOXP2, GABBR2, NRXN1*). Therefore, paired trials indicate that, despite trial order and blood sampling in both trials introducing noise in behavioral responses, local and foreign songs are consistently associated with differential gene expression in the blood. The functional significance of these genes, and the extent to which they reflect brain gene expression, should be a focus of future investigations.

### Implications for population divergence and speciation

Genes in the forkhead box (*FOX*) group of transcription factors such as *FOXP2* and others such as Neurexin 1 (*NRXN1*) are interesting candidate genes for population divergence^44,45^. In the brain, these genes are involved in vocal and auditory learning and vocal production, and numerous studies have demonstrated their importance for song learning and production in birds^29,30,45^. Moreover, *FOX* and neurexin genes are frequent targets of selection, forming hotspots of divergence between closely related taxa across vertebrates^46^. For instance, hybrids between species of *Pogoniulus* tinkerbirds with signals of admixture at *NRXN1* sing less stable songs with disrupted paces than non-admixed individuals, which may explain signals of selection and high differentiation at this locus^45^. Our detection of differential expression of *FOXP2, NRXN1* and other song-associated genes in the contrast of local vs. foreign song treatments suggest that these genes may be relevant in the early stages of population divergence. Genomic studies testing differentiation of song-associated loci and conducting behavioral assays, followed by gene expression analysis in the blood and the brain, will be necessary to validate the role of these genes in the speciation process.

### Limitations of the study

The present study has several limitations that should be considered when interpreting the results. Small sample sizes are one limitation, reducing the statistical power to estimate differential behavioral responses as well as expression profiles across treatments. Due to ethical concerns of testing an excessive number of wild songbirds, we were able to test a maximum of 15 individuals in the field with paired behavioral trials and blood sampling and 10 additional individuals that had their blood sampled but were not tested with behavioral trials. As mentioned above, sample sizes of blood samples were further reduced in second trials to 8 individuals due to the increased difficulty of capturing subjects following second trials after they had been captured, banded and bled in their first trials. Such change in behavior in second trials points to an additional limitation. Although paired designs are key to studies in behavioral ecology, and blood sampling is a less invasive method to measure gene expression^6,8,14^, capturing and blood sampling constitute a disturbance that not only affected the behavior of the birds during second trials but may also have influenced gene expression in their blood. Finally, because we did not sample brain tissue, as lethal sampling would make a paired design impossible, we cannot know whether the song-associated genes in the blood derive from the brain. Future studies sampling both the blood and the brain are necessary to test the correlation between gene expression in these tissues in relation to local and foreign songs.

## STAR Methods

### Behavioral trials

From May 5 to May 26, 2024, we conducted fieldwork in Crawford County, Pennsylvania, USA, on Pennsylvania State Gamelands 213 and 214, Pymatuning State Park, and the University of Pittsburgh’s Pymatuning Laboratory of Ecology. We followed simulated territorial intrusion (STI, i.e., “playback”) field protocols developed by ^27^ and ^8^. We conducted STIs with 15 male song sparrows using a within-subjects (paired), counterbalanced design of local and foreign songs 3–5 days apart. That is, we tested each of the 15 males twice with randomly ordered local and foreign song treatments, waiting 3–5 days between trials. We identified focal males by listening and observing natural vocal and territorial activities. Though we did not locate nests to identify egg or nestling stage, all males were singing on territories and had a cloacal protuberance, indicating they were actively conducting territory defense and were in breeding condition.

After identifying a focal male, we placed on the ground an Anchor AN-MINI loudspeaker connected to a Marantz PMD660 recorder close to a vegetation patch that we had observed the focal individual using as a singing perch. We placed the loudspeaker near the center of the focal birds’ territories, avoiding individuals in neighboring territories. The songs used as experimental stimuli were recorded by ^8^. Local songs were recorded in 1992–1994 in the vicinity of the Pymatuning Laboratory of Ecology in Linesville, Pennsylvania, and foreign songs were recorded in 1987–1988 in the Rockefeller Field Research Station in Millbrook, New York. Playbacks of songs recorded in 1987-1988 and in 2025 in the same localities did not elicit different behavioral responses from male song sparrows, indicating that the songs used in the present study constitute relevant social challenges (S. N.; unpublished results). We prepared three experimental tracks for the local population and three experimental tracks for the foreign population, and the experimenters in the field (G. M. and S. N.) were blind to the source population of the tracks. Each experimental track consisted of five different song types from the local or foreign populations, and each song type was played at the rate of 1 song per 10 s, with 10 repetitions per song until changing to a different song type. We used orange flagging placed on surrounding vegetation to mark distance categories of 0–2 m, 2–4 m, 4–8 m, 8–16 m, and >16 m from the loudspeaker. The distance that the focal individual was in relation to the loudspeaker was recorded in 5-second blocks for a total of 30 min, with the mean distance score for the trial calculated as in Peters et al. (1980). Distance to stimulus source has been shown to be the most reliable measure of territorial aggressive response in male song sparrows ^8,27,47^.

### Blood sampling

After a behavioral trial, we waited 30 min in silence (silent phase) for transcription to occur in the blood. Next, we captured the focal individual with a mist net, banded it with a United States Fish and Wildlife Service metal band and a unique combination of color bands, weighed it with a Pesola (to the neared 0.1 g), and took morphological measurements of the right tarsus and flattened right wing length with an ornithological ruler. Average time from the end of the silent phase to capture was 1.27 ± 0.52 min (mean ± SE) in first trials and 4.88 ± 1.68 min in second trials. Lastly, using nitrile gloves and tools cleaned with RNAse-away and 95% ethanol, we obtained a sample of 20 µL of blood via brachial venipuncture. We placed the blood sample in cryotubes (Thermo Fisher Scientific) and flash froze the tubes in dry ice (-78.5°C). Blood samples were later stored in a -80°C freezer until downstream analyses.

We conducted paired behavioral trials with all 15 focal individuals with local and foreign songs, except for one individual that was not found on its territory for the second trial. After capturing, banding, and obtaining a blood sample in their first trial, individuals became noticeably aware of the mist net and some avoided being captured a second time. If we could not capture a focal individual within 30 min after the silent phase, which could bias gene expression, we considered that the recapture attempt had failed ^17^. In total, we recaptured 8 of the 15 focal individuals but failed to recapture the 7 remaining individuals; one of which being the individual that was missing from its territory.

### Baseline (control) group

To obtain control, baseline gene expression profiles of individuals that were not exposed to experimental social challenges, we also captured and took blood samples from 10 additional male song sparrows with which we did not conduct behavioral trials, only luring them to the mist net with a short-duration song recording (less than 1 min). Baseline individuals were in similar breeding conditions to the tested individuals, i.e., defending territories and with developed cloacal protuberances. Additionally, capturing and obtaining a blood sample, although being relatively brief procedures, may on their own affect the stress level and gene expression of individuals irrespectively of the content of experimental stimuli. Baseline individuals therefore also serve to account for these other effects of the study design.

### Statistical analysis of behavioral trials

We compared the distance from the loudspeaker across treatments and trials with Bayesian multilevel models using *brms*^48,49^. Because distance from the loudspeaker was right skewed and not normally distributed, we subtracted the minimum value from each observation, so that the minimum distance values were at zero. We then used a model with the hurdle lognormal distribution family with the hurdle at zero. Hurdle lognormal models employ a two-step process: first, a logit link function models the binary outcome of whether the observation is zero or positive, and then positive values are modeled with a lognormal distribution with an identity link function^50,51^. The hurdle lognormal distribution provided a better fit to the data than a normal, lognormal, or skewed normal distributions.

As explanatory variables, we specified treatment (local or foreign song), trial order (first or second trial) as population-level (fixed) effects, and individual identification as a group-level (random) effect. We used weakly-informative normal priors for population-level effects and the intercept (*N* ∼ 0, 2; mean, SD), and gamma priors for the group-level effect and the residual standard deviation (*Gamma* ∼ 1, 1; rate, shape parameters). We ran models using four Markov chains, a thinning rate of one, and 10,000 iterations (50% as warm-up), and we evaluated model convergence and fit with trace plots, the Gelman-Rubin diagnostic, posterior predictive checks, and chain autocorrelation plots. We obtained P-values for the Bayesian model with the R package *bayestestR*^52^, which converts the Bayesian maximum probability of effect (or probability of direction) into a P-value. Because our sample size is small (15 individuals, each tested twice), we also ran a non-parametric kernel regression using the R package *np*^53^ to compare with the results of the Bayesian model. We also used treatment and trial order as fixed effects, but, because current implementations of the *np* package do not support random effects, we specified individual identification as a fixed effect. We tested the significance of effects in the non-parametric model with 1,000 bootstrap replicates^53^.

### RNA extraction

We extracted total RNA from whole blood using a RiboPure™ RNA Purification Kit (Ambion). After extraction, we added DNAse I and incubated samples at 37°C for 30 min to remove genomic DNA from the RNA. Average RNA concentration was 40.26 ng/uL. Using a TapeStation Fragment Analyzer (Agilent Technologies) with a high sensitivity RNA kit, the average RIN was 9.37.

### Library preparation and sequencing

We submitted total RNA to the Duke Sequencing and Genomics Technologies Core Facility (SGT). Though RNA studies of human blood typically involve hemoglobin depletion^54^, we opted instead to increase the number of reads, as there are currently no probes available for birds. Library preparation was conducted with a KAPA HyperPrep^55^ and included mRNA poly A enrichment. Paired end sequencing of 150bp reads was conducted using 3 lanes of a NovaSeq X Plus 10B lane to generate an average of 108.5 million reads per sample.

### Bioinformatics

RNA-seq data was processed using the *fastp* toolkit ^56^, trimming low-quality bases and sequencing adapters from the 3’ end of reads, then mapped to the song sparrow reference genome bMelMel2 (RefSeq assembly GCF_035770615.1^57^) using the STAR RNA-seq alignment tool^58^. Reads aligning to a single genomic location were summed across genes. We identified 21,141 total genes. As expected for blood, the most highly expressed genes were *LOC134426298* (hemoglobin subunit alpha-A) with an average of 11,438,449 counts, followed by *LOC134431771* (hemoglobin subunit beta) with 9,938,971, and *LOC134426297* (hemoglobin subunit alpha-D) with 5,234,548 counts.

### Differential expression analysis

We conducted differential expression analysis using the *edgeR* and *DESeq2* R packages^59,60^. Sample sizes in the first trials were 7 individuals in the local song treatment and 8 individuals in the foreign song treatment. As baseline controls, the sample size was 10 individuals. Paired trials (i.e., individuals sampled both in the first and second trials) had the sample size of 8 individuals. First, we investigated whether expression profiles differed between individuals tested in the first trials with local and foreign song treatments (the social challenge treatments) as well as baseline individuals (absence of social challenge). In the first trials, we retained genes with at least 3 counts per million in at least 7 samples (the lowest sample size in this case). Then, to investigate the effects of treatment and trial order on expression profiles, we compared local and foreign song treatments of paired trials. We accounted for repeated measures of paired samples by including individual identification in the model. In paired trials, we retained genes with at least 5 counts in at least 1 sample. Baseline samples were not included in the analysis of paired trials because baselines do not have a trial order, which would make the model’s matrix not full rank due to empty combinations of factors. After false discovery rate correction (FDR), we considered DEGs (differentially expressed genes) if P < 0.05. We performed gene set enrichment analysis with the R package *gprofiler2*^61^ to identify gene ontology terms and pathways associated with DEGs.

Because expression profiles often show coordinated changes of many interacting genes, analyses of individual DEGs may fail to capture these emergent properties^62^. Therefore, we conducted Weighted Gene Co-expression Network Analysis using the R package *WGCNA*^*62*^. We tested the effect of time to capture (in minutes), treatment/baseline (baseline, local song, foreign song), and, in paired trials, the effect of treatment and trial order. We filtered genes as in the analyses with *edgeR* and *DESeq2* above, normalized for read-depth and extracted variance stabilizing transformed (vst) read counts using *DESeq2*. We built the co-expression matrix using soft thresholding power (β_t_) values wherein mean connectivity plateaued (β_t_ = 12 in first trials; β_t_ = 9 in paired trials) to obtain scale-free topologies, and we merged highly correlated modules using a dissimilarity threshold of 0.25. We considered modules with P < 0.05 as significantly correlated with treatment, trial order, or time to capture. Then, we used the eigengene value, which is analogous to the first principal component of a module, as a response variable in Bayesian multilevel models and non-parametric kernel models to test the effect of treatment. In paired trials, we tested the effects of treatment and trial order with individual ID as a group-level effect (or fixed-effect in the non-parametric model).

## Resource Availability

### Lead contact

Inquiries about this study and its supporting data should be directed to G. M. and S. L.

### Materials availability

Raw sequencing data and raw gene counts for each sample are archived in NCBI GEO (Accession Number GSE330656; https://www.ncbi.nlm.nih.gov/geo/query/acc.cgi?acc=GSE330656).

### Data and code availability

Behavioral and gene count data and the R code used in the analyses reported in this study will be archived in Dryad.

## Acknowledgements

We thank the Duke University School of Medicine for the use of the Sequencing and Genomic Technologies Shared Resource, which provided library preparation, RNAseq, and bioinformatic services, especially to Joseph Prinz for bioinformatics, and Devi Swain Lenz for sequencing support. We are also grateful to the Pymatuning Lab of Ecology for the use of their facilities and hospitality during fieldwork, and the Pennsylvania State Game Commission and Pennsylvania Bureau of State Parks for access to field sites. We thank Tessa Patton for providing helpful code for RNAseq analysis and all members of the Lipshutz Lab for discussions and comments on earlier versions of the manuscript.

## Ethics Statement

All procedures used in this study were approved by the Duke University Institutional Animal Care and Use Committee (protocol #A077-24-04), with all applicable federal, state and local permits for land use and handling birds. The behavioral trials conducted in this study elicited normal behavioral responses of the animals; and capturing and blood sampling were conducted so as to minimize handling time, discomfort, and pain. The amount of blood sampled (20 μL) was far less than 1% of body mass, and the animals were promptly released in their natural habitat, in their territories, as soon as blood sampling had been completed.

## Declaration of generative AI and AI-assisted technologies

The authors did not use generative AI in any stage of this study or preparation of the manuscript.

## Resource Availability

### Lead contact

Inquiries about this study and its supporting data should be directed to G. M. and S. L.

### Data and code availability

Behavioral and gene count data and the R code used in the analyses reported in this study will be archived in Mendeley Data.

## Declaration of generative AI and AI-assisted technologies

The authors did not use generative AI in any stage of this study or preparation of the manuscript.

## References

1. Catchpole, C. K. & Slater, P. J. Bird Song: Biological Themes and Variations. (Cambridge university press, 2003).

2. Marler, P. R. & Slabbekoorn, H. Nature’s Music: The Science of Birdsong. (Elsevier, 2004).

3. Harris, M. A. & Lemon, R. E. Songs of song sparrows: Reactions of males to songs of different localities. The Condor 76, 33–44 (1974).

4. Parker, T. H., Greig, E. I., Nakagawa, S., Parra, M. & Dalisio, A. C. Subspecies status and methods explain strength of response to local versus foreign song by oscine birds in meta-analysis. Anim. Behav. 142, 1–17 (2018).

5. Wheatcroft, D. et al. Species-specific song responses emerge as a by-product of tuning to the local dialect. Curr. Biol. 32, 5153–5158 (2022).

6. Searcy, W. A., Nowicki, S., Hughes, M. & Peters, S. Geographic Song Discrimination in Relation to Dispersal Distances in Song Sparrows. Am. Nat. 159, 221–230 (2002).

7. Lipshutz, S. E., Overcast, I. A., Hickerson, M. J., Brumfield, R. T. & Derryberry, E. P. Behavioural response to song and genetic divergence in two subspecies of whiteLcrowned sparrows (*Zonotrichia leucophrys*). Mol. Ecol. 26, 3011–3027 (2017).

8. Searcy, W. A., Nowicki, S. & Hughes, M. The response of male and female song sparrows to geographic variation in song. The Condor 99, 651–657 (1997).

9. Searcy, W. A. & Nowicki, S. Animal Communication. in The Behavior of Animals, 2nd Edition (eds Bolhuis, J. J., Giraldeau, L. & Hogan, J. A.) 367–396 (Wiley, 2021). doi:10.1002/9781119109556.ch14.

10. Bukhari, S. A. et al. Temporal dynamics of neurogenomic plasticity in response to social interactions in male threespined sticklebacks. PLoS Genet. 13, e1006840 (2017).

11. Jarvis, E. D., Schwabl, H., Ribeiro, S. & Mello, C. V. Brain gene regulation by territorial singing behavior in freely ranging songbirds. Neuroreport 8, 2073–2077 (1997).

12. Rittschof, C. C. et al. Neuromolecular responses to social challenge: Common mechanisms across mouse, stickleback fish, and honey bee. Proc. Natl. Acad. Sci. 111, 17929–17934 (2014).

13. George, E. M. & Rosvall, K. A. How a territorial challenge changes sex steroid-related gene networks in the female brain: A field experiment with the tree swallow. Horm. Behav. 169, 105698 (2025).

14. Louder, M. I. et al. Shared transcriptional responses to con-and heterospecific behavioral antagonists in a wild songbird. Sci. Rep. 10, 4092 (2020).

15. Kirian, R. D., Steinman, D., Jewell, C. M. & Zierden, H. C. Extracellular vesicles as carriers of mRNA: Opportunities and challenges in diagnosis and treatment. Theranostics 14, 2265 (2024).

16. Smirnova, L. et al. Blood extracellular vesicles carrying brain-specific mRNAs are potential biomarkers for detecting gene expression changes in the female brain. Mol. Psychiatry 29, 962–973 (2024).

17. Louder, M. I., Hauber, M. E. & Balakrishnan, C. N. Early social experience alters transcriptomic responses to species-specific song stimuli in female songbirds. Behav. Brain Res. 347, 69–76 (2018).

18. Sun, A.-G. et al. Identifying distinct candidate genes for early Parkinson’s disease by analysis of gene expression in whole blood. Neuro Endocrinol Lett 35, 398–404 (2014).

19. Tylee, D. S., Kawaguchi, D. M. & Glatt, S. J. On the outside, looking in: A review and evaluation of the comparability of blood and brain “Lomes”. Am. J. Med. Genet. B Neuropsychiatr. Genet. 162, 595–603 (2013).

20. Anderson, J. A. et al. High social status males experience accelerated epigenetic aging in wild baboons. Elife 10, e66128 (2021).

21. Campbell, C. R. et al. A femaleLbiased gene expression signature of dominance in cooperatively breeding meerkats. Mol. Ecol. 33, e17467 (2024).

22. Huang, Z. et al. A nonlethal sampling method to obtain, generate and assemble whole blood transcriptomes from small, wild mammals. Mol. Ecol. Resour. 16, 150–162 (2016).

23. Luo, J. et al. Blood transcriptome analysis revealing aging gene expression profiles in red panda. PeerJ 10, e13743 (2022).

24. Shah, S. A. U. R. et al. Comparative blood transcriptome analysis reveals changes in immunity, and transcripts related to metabolism and development in the critically endangered Yangtze finless porpoise (Neophocaena asiaeorientalis asiaeorientalis) with age. Comp. Immunol. Rep. 200255 (2025).

25. Arcese, P., Sogge, M. K., Marr, A. B. & Patten, M. A. Song Sparrow (Melospiza melodia), version 1.0. Birds World https://doi.org/10.2173/bow.sonspa.01 (2020) doi:10.2173/bow.sonspa.01.

26. Hughes, M., Nowicki, S., Searcy, W. A. & Peters, S. Song-type sharing in song sparrows: implications for repertoire function and song learning. Behav. Ecol. Sociobiol. 42, 437–446 (1998).

27. Peters, S. S., Searcy, W. A. & Marler, P. Species song discrimination in choice experiments with territorial male swamp and song sparrows. Anim. Behav. 28, 393–404 (1980).

28. Marler, P. & Peters, S. The Role of Song Phonology and Syntax in Vocal Learning Preferences in the Song Sparrow, *Melospiza melodia*. Ethology 77, 125–149 (1988).

29. Konopka, G. et al. Human-specific transcriptional networks in the brain. Neuron 75, 601–617 (2012).

30. Scharff, C. & Haesler, S. An evolutionary perspective on FoxP2: strictly for the birds? Curr. Opin. Neurobiol. 15, 694–703 (2005).

31. Herrero, M. J. & Gitton, Y. The untold stories of the speech gene, the FOXP2 cancer gene. Genes Cancer 9, 11 (2018).

32. Liu, S., Gao, M., Zhang, X., Wei, J. & Cui, H. FOXP2 overexpression upregulates LAMA4 expression and thereby alleviates preeclampsia by regulating trophoblast behavior. Commun. Biol. 7, 1427 (2024).

33. Zhang, H., Zhou, H., Yuan, J., Nan, Y. & Liu, J. Endothelial GABBR2 regulates postischemic angiogenesis by inhibiting the glycolysis pathway. Front. Cardiovasc. Med. 8, 696578 (2021).

34. Kuwano, Y. et al. Autism-associated gene expression in peripheral leucocytes commonly observed between subjects with autism and healthy women having autistic children. PloS One 6, e24723 (2011).

35. Orije, J. E. M. J. & Van Der Linden, A. A brain for all seasons: An in vivo MRI perspective on songbirds. J. Exp. Zool. Part Ecol. Integr. Physiol. 337, 967–984 (2022).

36. Thompson, C. K. et al. Seasonal changes in patterns of gene expression in avian song control brain regions. PloS One 7, e35119 (2012).

37. Sun, B. et al. Non canonical BRCA1 promotes cell survival via modulating PARP13-mediated SEC61G mRNA decay. Oncogenesis 14, 35 (2025).

38. Williams, A. G., Thomas, S., Wyman, S. K. & Holloway, A. K. RNALseq Data: Challenges in and Recommendations for Experimental Design and Analysis. Curr. Protoc. Hum. Genet. 83, (2014).

39. Bae, I. et al. BRCA1 induces antioxidant gene expression and resistance to oxidative stress. Cancer Res. 64, 7893–7909 (2004).

40. McGregor, P. K. Playback experiments: design and analysis. Acta Ethologica 3, 3–8 (2000).

41. Bai, Y. et al. A novel tumor-suppressor, CDH18, inhibits glioma cell invasiveness via UQCRC2 and correlates with the prognosis of glioma patients. Cell. Physiol. Biochem. 48, 1755–1770 (2018).

42. Junghof, J. et al. CDH18 is a fetal epicardial biomarker regulating differentiation towards vascular smooth muscle cells. Npj Regen. Med. 7, 14 (2022).

43. Yan, X. et al. Cadm2 regulates body weight and energy homeostasis in mice. Mol. Metab. 8, 180–188 (2018).

44. Kozma, R., Rödin-Mörch, P. & Höglund, J. Genomic regions of speciation and adaptation among three species of grouse. Sci. Rep. 9, 812 (2019).

45. Sebastianelli, M. et al. A genomic basis of vocal rhythm in birds. Nat. Commun. 15, 3095 (2024).

46. Song, X., Wang, Y. & Tang, Y. Rapid diversification of FOXP2 in teleosts through gene duplication in the teleost-specific whole genome duplication event. PLoS One 8, e83858 (2013).

47. Searcy, W. A., McArthur, P. D., Peters, S. S. & Marler, P. Response of male song and swamp sparrows to neighbour, stranger, and self songs. Behaviour 77, 152–163 (1981).

48. Bürkner, P.-C. brms: An R Package for Bayesian Multilevel Models Using Stan. J. Stat. Softw. 80, 1–28 (2017).

49. Bürkner, P.-C. et al. brms: Bayesian Regression Models using ‘Stan’. (2024).

50. Mullahy, J. Specification and testing of some modified count data models. J. Econom. 33, 341–365 (1986).

51. Rose, C. E., Martin, S. W., Wannemuehler, K. A. & Plikaytis, B. D. On the Use of Zero-Inflated and Hurdle Models for Modeling Vaccine Adverse Event Count Data. J. Biopharm. Stat. 16, 463–481 (2006).

52. Makowski, D., Ben-Shachar, M. & Lüdecke, D. bayestestR: Describing Effects and their Uncertainty, Existence and Significance within the Bayesian Framework. J. Open Source Softw. 4, 1541 (2019).

53. Racine, J. S. & Hayfield, T. np: Nonparametric Kernel Smoothing Methods for Mixed Data Types. (2026).

54. Shin, H. et al. Variation in RNA-Seq transcriptome profiles of peripheral whole blood from healthy individuals with and without globin depletion. PloS One 9, e91041 (2014).

55. Van Kets, V., Kitzman, J., Snyder, M., Shendure, J. & Gray, P. Kapa Hyper Prep: A next-generation kit for fast and efficient library construction from challenging DNA samples. in Proceedings of the Advances in Genome Biology and Technology (AGBT) Meeting, Marco Island, FL, USA 12–15 (2014).

56. Chen, S. fastp 1.0: An ultraLfast allLround tool for FASTQ data quality control and preprocessing. iMeta 4, e70078 (2025).

57. Louha, S., Ray, D. A., Winker, K. & Glenn, T. C. A high-quality genome assembly of the North American Song Sparrow, Melospiza melodia. G3 Genes Genomes Genet. 10, 1159–1166 (2020).

58. Dobin, A. et al. STAR: ultrafast universal RNA-seq aligner. Bioinformatics 29, 15–21 (2013).

59. Chen, Y., Chen, L., Lun, A. T., Baldoni, P. L. & Smyth, G. K. edgeR v4: powerful differential analysis of sequencing data with expanded functionality and improved support for small counts and larger datasets. Nucleic Acids Res. 53, gkaf018 (2025).

60. Love, M., Anders, S. & Huber, W. Differential analysis of count data–the DESeq2 package. Genome Biol 15, 10–1186 (2014).

61. Kolberg, L., Raudvere, U., Kuzmin, I., Vilo, J. & Peterson, H. gprofiler2–an R package for gene list functional enrichment analysis and namespace conversion toolset g: Profiler. F1000Research 9, ELIXIR-709 (2020).

62. Langfelder, P. & Horvath, S. WGCNA: an R package for weighted correlation network analysis. BMC Bioinformatics 9, 559 (2008).

