## Supplemental Figures and Tables for "Response to geographic variation in song is associated with differential gene expression in the blood of a songbird"

**Supplementary Material**

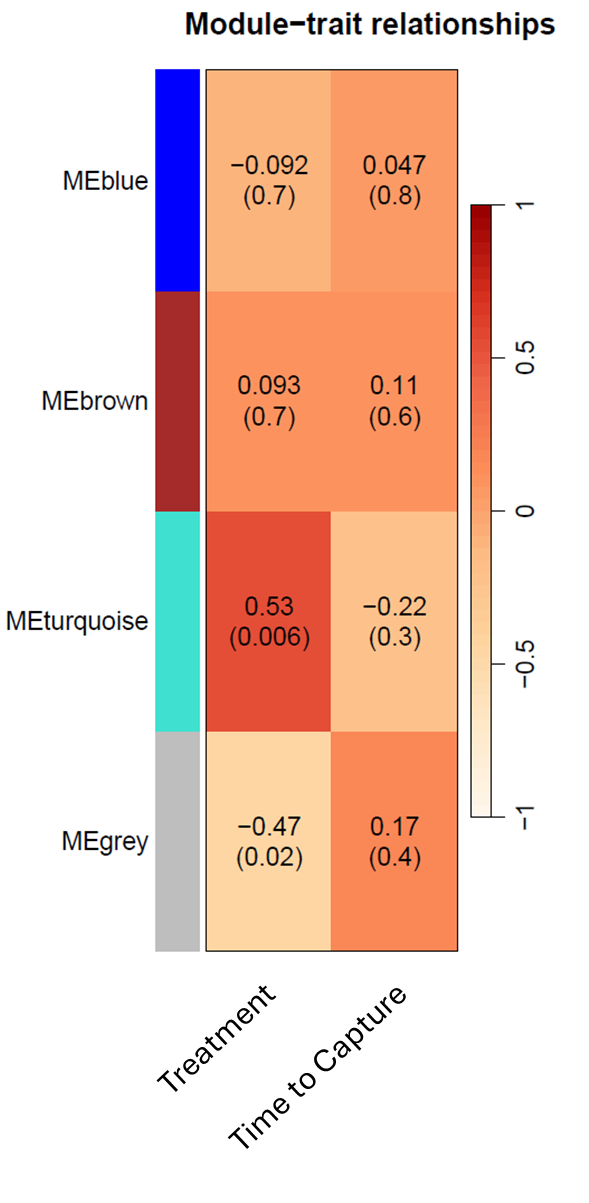

**Figure S1.** Module-trait relationships of the WGCNA analysis with baseline individuals, and individuals in the local and foreign song treatments in the first trials.

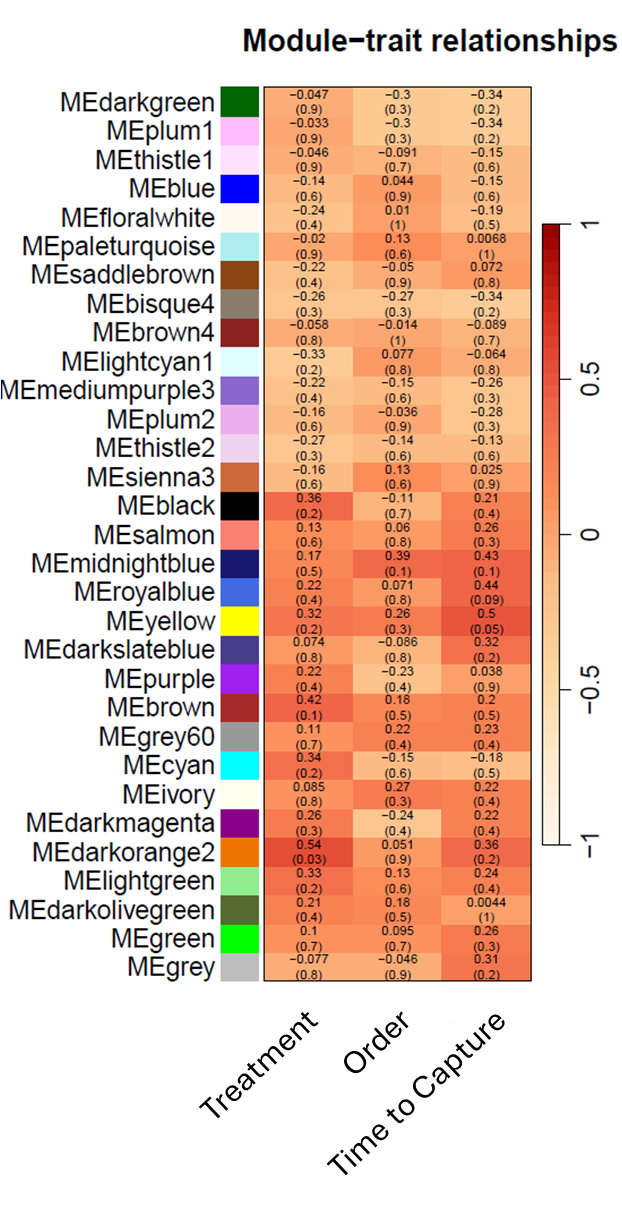

**Figure S2.** Module-trait relationships of the WGCNA analysis with individuals in the local and foreign song treatments in the paired trials.

**Table S1.** Gene ontology terms (GO.ID) significantly associated with up-regulated genes in the contrast between individuals in the local and foreign song treatments in the first trials. P (FDR) are P-values after false discovery rate correction.

| GO.ID | Description | P | P (FDR) | Genes |
| --- | --- | --- | --- | --- |
| CORUM:7344 | NLGN1-NRXN1 complex | 0.003 | 0.003 | NRXN1 |
| CORUM:828 | TRPC1-STIM1-ORAI1 complex | 0.003 | 0.003 | CTNND2 |
| CORUM:5418 | GABBR2-HTR1A complex | 0.003 | 0.003 | GABBR2 |
| CORUM:6437 | GABBR1-GABBR2 complex | 0.003 | 0.003 | GABBR2 |
| GO:0071625 | vocalization behavior | <0.0001 | <0.0001 | NRXN1, CNTNAP2, GLI3 |
| GO:0099560 | synaptic membrane adhesion | 0.001 | 0.001 | NRXN1, LRRC4C, IL1RAPL1 |
| GO:0034330 | cell junction organization | 0.001 | 0.001 | NRXN1, CNTNAP2, LRRC4C, IL1RAPL1, NRG3, DGKB, CTNND2 |
| GO:0042297 | vocal learning | 0.001 | 0.001 | NRXN1, CNTNAP2 |
| GO:0001964 | startle response | 0.001 | 0.001 | NRXN1, CNTNAP2, CSMD1 |
| GO:0021544 | subpallium development | 0.001 | 0.001 | CNTNAP2, FOXP2, GLI3 |
| GO:0098596 | imitative learning | 0.001 | 0.001 | NRXN1, CNTNAP2 |
| GO:0050808 | synapse organization | 0.001 | 0.001 | NRXN1, LRRC4C, IL1RAPL1, NRG3, DGKB, CTNND2 |
| GO:0098597 | observational learning | 0.001 | 0.001 | NRXN1, CNTNAP2 |
| GO:0060443 | mammary gland morphogenesis | 0.001 | 0.001 | CSMD1, GLI3, NRG3 |
| GO:0098598 | learned vocalization behavior or vocal learning | 0.001 | 0.001 | NRXN1, CNTNAP2 |
| GO:0060592 | mammary gland formation | 0.001 | 0.001 | GLI3, NRG3 |
| GO:0048858 | cell projection morphogenesis | 0.001 | 0.001 | NRXN1, CNTNAP2, LRRC4C, IL1RAPL1, GLI3, CTNND2 |
| GO:0048812 | neuron projection morphogenesis | 0.001 | 0.001 | NRXN1, CNTNAP2, LRRC4C, IL1RAPL1, GLI3, CTNND2 |
| GO:0120039 | plasma membrane bounded cell projection morphogenesis | 0.001 | 0.001 | NRXN1, CNTNAP2, LRRC4C, IL1RAPL1, GLI3, CTNND2 |
| GO:0097105 | presynaptic membrane assembly | 0.002 | 0.002 | NRXN1, IL1RAPL1 |
| GO:0031223 | auditory behavior | 0.002 | 0.002 | NRXN1, CNTNAP2 |
| GO:0099537 | trans-synaptic signaling | 0.002 | 0.002 | NRXN1, LRRC4C, IL1RAPL1, GABBR2, NRG3, DGKB |
| GO:0099536 | synaptic signaling | 0.002 | 0.002 | NRXN1, LRRC4C, IL1RAPL1, GABBR2, NRG3, DGKB |
| GO:0097090 | presynaptic membrane organization | 0.002 | 0.002 | NRXN1, IL1RAPL1 |
| GO:0060322 | head development | 0.002 | 0.002 | NRXN1, CNTNAP2, FOXP2, GLI3, NRG3, CSRNP1 |
| GO:0007638 | mechanosensory behavior | 0.002 | 0.002 | NRXN1, CNTNAP2 |
| GO:0099173 | postsynapse organization | 0.003 | 0.003 | NRXN1, IL1RAPL1, DGKB, CTNND2 |
| GO:0060134 | prepulse inhibition | 0.003 | 0.003 | NRXN1, CNTNAP2 |
| GO:0021537 | telencephalon development | 0.004 | 0.004 | CNTNAP2, FOXP2, GLI3, NRG3 |
| GO:1901890 | positive regulation of cell junction assembly | 0.005 | 0.005 | NRXN1, CNTNAP2, IL1RAPL1 |
| GO:0009653 | anatomical structure morphogenesis | 0.005 | 0.005 | NRXN1, CNTNAP2, LRRC4C, CSMD1, IL1RAPL1, GLI3, NRG3, CSRNP1, CTNND2 |
| GO:0000902 | cell morphogenesis | 0.006 | 0.006 | NRXN1, CNTNAP2, LRRC4C, IL1RAPL1, GLI3, CTNND2 |
| GO:0048667 | cell morphogenesis involved in neuron differentiation | 0.006 | 0.006 | NRXN1, LRRC4C, IL1RAPL1, GLI3, CTNND2 |
| GO:0031175 | neuron projection development | 0.006 | 0.006 | NRXN1, CNTNAP2, LRRC4C, IL1RAPL1, GLI3, CTNND2 |
| GO:0048699 | generation of neurons | 0.007 | 0.007 | NRXN1, CNTNAP2, LRRC4C, IL1RAPL1, GLI3, NRG3, CTNND2 |
| GO:0021756 | striatum development | 0.007 | 0.007 | CNTNAP2, FOXP2 |
| GO:0021987 | cerebral cortex development | 0.007 | 0.007 | CNTNAP2, FOXP2, GLI3 |
| GO:0030879 | mammary gland development | 0.007 | 0.007 | CSMD1, GLI3, NRG3 |
| GO:0022612 | gland morphogenesis | 0.007 | 0.007 | CSMD1, GLI3, NRG3 |
| GO:0010996 | response to auditory stimulus | 0.008 | 0.008 | NRXN1, CNTNAP2 |
| GO:0098916 | anterograde trans-synaptic signaling | 0.009 | 0.009 | NRXN1, LRRC4C, GABBR2, NRG3, DGKB |
| GO:0099175 | regulation of postsynapse organization | 0.009 | 0.009 | NRXN1, IL1RAPL1, DGKB |
| GO:0007268 | chemical synaptic transmission | 0.009 | 0.009 | NRXN1, LRRC4C, GABBR2, NRG3, DGKB |
| GO:0007420 | brain development | 0.009 | 0.009 | NRXN1, CNTNAP2, FOXP2, GLI3, NRG3 |
| GO:0050905 | neuromuscular process | 0.009 | 0.009 | NRXN1, CNTNAP2, CSMD1 |
| GO:0007612 | learning | 0.009 | 0.009 | NRXN1, CNTNAP2, CSMD1 |
| GO:0048666 | neuron development | 0.009 | 0.009 | NRXN1, CNTNAP2, LRRC4C, IL1RAPL1, GLI3, CTNND2 |
| GO:0048731 | system development | 0.010 | 0.010 | NRXN1, CNTNAP2, LRRC4C, CSMD1, IL1RAPL1, FOXP2, GLI3, NRG3, CSRNP1, CTNND2 |
| GO:1905606 | regulation of presynapse assembly | 0.010 | 0.010 | NRXN1, IL1RAPL1 |
| GO:0099174 | regulation of presynapse organization | 0.010 | 0.010 | NRXN1, IL1RAPL1 |
| GO:0030900 | forebrain development | 0.010 | 0.010 | CNTNAP2, FOXP2, GLI3, NRG3 |
| GO:0098609 | cell-cell adhesion | 0.010 | 0.010 | NRXN1, LRRC4C, IL1RAPL1, GLI3, CTNND2 |
| GO:0007267 | cell-cell signaling | 0.010 | 0.010 | NRXN1, LRRC4C, IL1RAPL1, GABBR2, NRG3, DGKB |
| GO:0022008 | neurogenesis | 0.010 | 0.010 | NRXN1, CNTNAP2, LRRC4C, IL1RAPL1, GLI3, NRG3, CTNND2 |
| GO:0007155 | cell adhesion | 0.011 | 0.011 | NRXN1, CNTNAP2, LRRC4C, IL1RAPL1, GLI3, CTNND2 |
| GO:0098742 | cell-cell adhesion via plasma-membrane adhesion molecules | 0.012 | 0.012 | NRXN1, LRRC4C, IL1RAPL1 |
| GO:0021543 | pallium development | 0.012 | 0.012 | CNTNAP2, FOXP2, GLI3 |
| GO:0007399 | nervous system development | 0.012 | 0.012 | NRXN1, CNTNAP2, LRRC4C, IL1RAPL1, FOXP2, GLI3, NRG3, CTNND2 |
| GO:0032501 | multicellular organismal process | 0.012 | 0.012 | NRXN1, CNTNAP2, LRRC4C, CSMD1, IL1RAPL1, FOXP2, GLI3, NRG3, TSC22D3, CSRNP1, DGKB, CTNND2 |
| GO:1905608 | positive regulation of presynapse assembly | 0.013 | 0.013 | NRXN1 |
| GO:0060875 | lateral semicircular canal development | 0.013 | 0.013 | GLI3 |
| GO:0050804 | modulation of chemical synaptic transmission | 0.013 | 0.013 | NRXN1, LRRC4C, NRG3, DGKB |
| GO:0060366 | lambdoid suture morphogenesis | 0.013 | 0.013 | GLI3 |
| GO:0060873 | anterior semicircular canal development | 0.013 | 0.013 | GLI3 |
| GO:0090126 | protein-containing complex assembly involved in synapse maturation | 0.013 | 0.013 | NRXN1 |
| GO:0070231 | T cell apoptotic process | 0.013 | 0.013 | GLI3, TSC22D3 |
| GO:0060367 | sagittal suture morphogenesis | 0.013 | 0.013 | GLI3 |
| GO:0051703 | biological process involved in intraspecies interaction between organisms | 0.013 | 0.013 | NRXN1, CNTNAP2 |
| GO:0021840 | directional guidance of interneurons involved in migration from the subpallium to the cortex | 0.013 | 0.013 | NRG3 |
| GO:0021842 | chemorepulsion involved in interneuron migration from the subpallium to the cortex | 0.013 | 0.013 | NRG3 |
| GO:0035176 | social behavior | 0.013 | 0.013 | NRXN1, CNTNAP2 |
| GO:0021757 | caudate nucleus development | 0.013 | 0.013 | FOXP2 |
| GO:0034329 | cell junction assembly | 0.013 | 0.013 | NRXN1, CNTNAP2, IL1RAPL1, NRG3 |
| GO:0021758 | putamen development | 0.013 | 0.013 | FOXP2 |
| GO:1905518 | regulation of presynaptic active zone assembly | 0.013 | 0.013 | NRXN1 |
| GO:0150036 | regulation of trans-synaptic signaling by endocannabinoid, modulating synaptic transmission | 0.013 | 0.013 | NRXN1 |
| GO:0099177 | regulation of trans-synaptic signaling | 0.013 | 0.013 | NRXN1, LRRC4C, NRG3, DGKB |
| GO:0099054 | presynapse assembly | 0.013 | 0.013 | NRXN1, IL1RAPL1 |
| GO:0022018 | lateral ganglionic eminence cell proliferation | 0.013 | 0.013 | GLI3 |
| GO:0022012 | subpallium cell proliferation in forebrain | 0.013 | 0.013 | GLI3 |
| GO:1905520 | positive regulation of presynaptic active zone assembly | 0.013 | 0.013 | NRXN1 |
| GO:0071709 | membrane assembly | 0.015 | 0.015 | NRXN1, IL1RAPL1 |
| GO:0030182 | neuron differentiation | 0.016 | 0.016 | NRXN1, CNTNAP2, LRRC4C, IL1RAPL1, GLI3, CTNND2 |
| GO:0060997 | dendritic spine morphogenesis | 0.016 | 0.016 | IL1RAPL1, CTNND2 |
| GO:0007416 | synapse assembly | 0.016 | 0.016 | NRXN1, IL1RAPL1, NRG3 |
| GO:0044091 | membrane biogenesis | 0.016 | 0.016 | NRXN1, IL1RAPL1 |
| GO:0051932 | synaptic transmission, GABAergic | 0.016 | 0.016 | NRXN1, GABBR2 |
| GO:0099172 | presynapse organization | 0.016 | 0.016 | NRXN1, IL1RAPL1 |
| GO:0007275 | multicellular organism development | 0.017 | 0.017 | NRXN1, CNTNAP2, LRRC4C, CSMD1, IL1RAPL1, FOXP2, GLI3, NRG3, CSRNP1, CTNND2 |
| GO:0051965 | positive regulation of synapse assembly | 0.017 | 0.017 | NRXN1, IL1RAPL1 |
| GO:0070227 | lymphocyte apoptotic process | 0.017 | 0.017 | GLI3, TSC22D3 |
| GO:1901888 | regulation of cell junction assembly | 0.018 | 0.018 | NRXN1, CNTNAP2, IL1RAPL1 |
| GO:0022029 | telencephalon cell migration | 0.019 | 0.019 | GLI3, NRG3 |
| GO:0007417 | central nervous system development | 0.019 | 0.019 | NRXN1, CNTNAP2, FOXP2, GLI3, NRG3 |
| GO:0021885 | forebrain cell migration | 0.020 | 0.020 | GLI3, NRG3 |
| GO:0007611 | learning or memory | 0.020 | 0.020 | NRXN1, CNTNAP2, CSMD1 |
| GO:0071109 | superior temporal gyrus development | 0.020 | 0.020 | CNTNAP2 |
| GO:0097117 | guanylate kinase-associated protein clustering | 0.020 | 0.020 | NRXN1 |
| GO:0120224 | larynx development | 0.020 | 0.020 | GLI3 |
| GO:0120223 | larynx morphogenesis | 0.020 | 0.020 | GLI3 |
| GO:0120036 | plasma membrane bounded cell projection organization | 0.020 | 0.020 | NRXN1, CNTNAP2, LRRC4C, IL1RAPL1, GLI3, CTNND2 |
| GO:0097118 | neuroligin clustering involved in postsynaptic membrane assembly | 0.020 | 0.020 | NRXN1 |
| GO:0030030 | cell projection organization | 0.022 | 0.022 | NRXN1, CNTNAP2, LRRC4C, IL1RAPL1, GLI3, CTNND2 |
| GO:0051962 | positive regulation of nervous system development | 0.022 | 0.022 | NRXN1, IL1RAPL1, GLI3 |
| GO:0007610 | behavior | 0.023 | 0.023 | NRXN1, CNTNAP2, CSMD1, GLI3 |
| GO:0070236 | negative regulation of activation-induced cell death of T cells | 0.025 | 0.025 | TSC22D3 |
| GO:0021775 | smoothened signaling pathway involved in ventral spinal cord interneuron specification | 0.025 | 0.025 | GLI3 |
| GO:0035846 | oviduct epithelium development | 0.025 | 0.025 | CSMD1 |
| GO:0043585 | nose morphogenesis | 0.025 | 0.025 | GLI3 |
| GO:0021910 | smoothened signaling pathway involved in ventral spinal cord patterning | 0.025 | 0.025 | GLI3 |
| GO:1900073 | regulation of neuromuscular synaptic transmission | 0.025 | 0.025 | NRXN1 |
| GO:0097116 | gephyrin clustering involved in postsynaptic density assembly | 0.025 | 0.025 | NRXN1 |
| GO:0097061 | dendritic spine organization | 0.025 | 0.025 | IL1RAPL1, CTNND2 |
| GO:1900075 | positive regulation of neuromuscular synaptic transmission | 0.025 | 0.025 | NRXN1 |
| GO:0021776 | smoothened signaling pathway involved in spinal cord motor neuron cell fate specification | 0.025 | 0.025 | GLI3 |
| GO:0070235 | regulation of activation-induced cell death of T cells | 0.025 | 0.025 | TSC22D3 |
| GO:0050890 | cognition | 0.025 | 0.025 | NRXN1, CNTNAP2, CSMD1 |
| GO:0050807 | regulation of synapse organization | 0.025 | 0.025 | NRXN1, IL1RAPL1, DGKB |
| GO:0060596 | mammary placode formation | 0.025 | 0.025 | NRG3 |
| GO:0060594 | mammary gland specification | 0.025 | 0.025 | GLI3 |
| GO:0050803 | regulation of synapse structure or activity | 0.025 | 0.025 | NRXN1, IL1RAPL1, DGKB |
| GO:0060364 | frontal suture morphogenesis | 0.025 | 0.025 | GLI3 |
| GO:0106027 | neuron projection organization | 0.026 | 0.026 | IL1RAPL1, CTNND2 |
| GO:0071887 | leukocyte apoptotic process | 0.027 | 0.027 | GLI3, TSC22D3 |
| GO:0060021 | roof of mouth development | 0.028 | 0.028 | GLI3, CSRNP1 |
| GO:0097112 | gamma-aminobutyric acid receptor clustering | 0.030 | 0.030 | NRXN1 |
| GO:0071205 | protein localization to juxtaparanode region of axon | 0.030 | 0.030 | CNTNAP2 |
| GO:1904071 | presynaptic active zone assembly | 0.030 | 0.030 | NRXN1 |
| GO:0060996 | dendritic spine development | 0.030 | 0.030 | IL1RAPL1, CTNND2 |
| GO:0097114 | NMDA glutamate receptor clustering | 0.030 | 0.030 | NRXN1 |
| GO:0099553 | trans-synaptic signaling by endocannabinoid, modulating synaptic transmission | 0.030 | 0.030 | NRXN1 |
| GO:0045163 | clustering of voltage-gated potassium channels | 0.030 | 0.030 | CNTNAP2 |
| GO:0099552 | trans-synaptic signaling by lipid, modulating synaptic transmission | 0.030 | 0.030 | NRXN1 |
| GO:0060066 | oviduct development | 0.036 | 0.036 | CSMD1 |
| GO:2000821 | regulation of grooming behavior | 0.036 | 0.036 | NRXN1 |
| GO:0150099 | neuron-glial cell signaling | 0.036 | 0.036 | GABBR2 |
| GO:0051490 | negative regulation of filopodium assembly | 0.036 | 0.036 | NRXN1 |
| GO:0099545 | trans-synaptic signaling by trans-synaptic complex | 0.041 | 0.041 | IL1RAPL1 |
| GO:0038138 | ERBB4-ERBB4 signaling pathway | 0.041 | 0.041 | NRG3 |
| GO:0021631 | optic nerve morphogenesis | 0.041 | 0.041 | GLI3 |
| GO:1900019 | regulation of protein kinase C activity | 0.041 | 0.041 | NRXN1 |
| GO:1900020 | positive regulation of protein kinase C activity | 0.041 | 0.041 | NRXN1 |
| GO:1990708 | conditioned place preference | 0.041 | 0.041 | CSMD1 |
| GO:0021761 | limbic system development | 0.041 | 0.041 | CNTNAP2, GLI3 |
| GO:0060831 | smoothened signaling pathway involved in dorsal/ventral neural tube patterning | 0.041 | 0.041 | GLI3 |
| GO:0023041 | neuronal signal transduction | 0.041 | 0.041 | NRXN1 |
| GO:0021861 | forebrain radial glial cell differentiation | 0.041 | 0.041 | GLI3 |
| GO:0021521 | ventral spinal cord interneuron specification | 0.043 | 0.043 | GLI3 |
| GO:1903010 | regulation of bone development | 0.043 | 0.043 | GLI3 |
| GO:1903598 | positive regulation of gap junction assembly | 0.043 | 0.043 | CNTNAP2 |
| GO:0060573 | cell fate specification involved in pattern specification | 0.043 | 0.043 | GLI3 |
| GO:0007154 | cell communication | 0.043 | 0.043 | NRXN1, CNTNAP2, LRRC4C, IL1RAPL1, GABBR2, GLI3, NRG3, CSRNP1, DGKB, CTNND2 |
| GO:0007442 | hindgut morphogenesis | 0.043 | 0.043 | GLI3 |
| GO:0021830 | interneuron migration from the subpallium to the cortex | 0.043 | 0.043 | NRG3 |
| GO:0060745 | mammary gland branching involved in pregnancy | 0.043 | 0.043 | CSMD1 |
| GO:1990709 | presynaptic active zone organization | 0.043 | 0.043 | NRXN1 |
| GO:2001223 | negative regulation of neuron migration | 0.043 | 0.043 | NRG3 |
| GO:0021798 | forebrain dorsal/ventral pattern formation | 0.043 | 0.043 | GLI3 |
| GO:0097104 | postsynaptic membrane assembly | 0.043 | 0.043 | NRXN1 |
| GO:0021843 | substrate-independent telencephalic tangential interneuron migration | 0.043 | 0.043 | NRG3 |
| GO:0021826 | substrate-independent telencephalic tangential migration | 0.043 | 0.043 | NRG3 |
| GO:0030534 | adult behavior | 0.045 | 0.045 | NRXN1, CNTNAP2 |
| GO:0021707 | cerebellar granule cell differentiation | 0.045 | 0.045 | NRXN1 |
| GO:0045743 | positive regulation of fibroblast growth factor receptor signaling pathway | 0.045 | 0.045 | NRXN1 |
| GO:0021684 | cerebellar granular layer formation | 0.045 | 0.045 | NRXN1 |
| GO:0006924 | activation-induced cell death of T cells | 0.045 | 0.045 | TSC22D3 |
| GO:0048702 | embryonic neurocranium morphogenesis | 0.045 | 0.045 | GLI3 |
| GO:0048732 | gland development | 0.045 | 0.045 | CSMD1, GLI3, NRG3 |
| GO:0097119 | postsynaptic density protein 95 clustering | 0.045 | 0.045 | NRXN1 |
| GO:0090129 | positive regulation of synapse maturation | 0.045 | 0.045 | NRXN1 |
| GO:0048856 | anatomical structure development | 0.045 | 0.045 | NRXN1, CNTNAP2, LRRC4C, CSMD1, IL1RAPL1, FOXP2, GLI3, NRG3, CSRNP1, CTNND2 |
| GO:0051960 | regulation of nervous system development | 0.045 | 0.045 | NRXN1, IL1RAPL1, GLI3 |
| GO:0061525 | hindgut development | 0.045 | 0.045 | GLI3 |
| GO:0060872 | semicircular canal development | 0.045 | 0.045 | GLI3 |
| GO:0060363 | cranial suture morphogenesis | 0.045 | 0.045 | GLI3 |
| GO:0099541 | trans-synaptic signaling by lipid | 0.045 | 0.045 | NRXN1 |
| GO:0099542 | trans-synaptic signaling by endocannabinoid | 0.045 | 0.045 | NRXN1 |
| GO:0048813 | dendrite morphogenesis | 0.045 | 0.045 | IL1RAPL1, CTNND2 |
| GO:0048513 | animal organ development | 0.046 | 0.046 | NRXN1, CNTNAP2, CSMD1, FOXP2, GLI3, NRG3, CSRNP1 |
| GO:1903596 | regulation of gap junction assembly | 0.048 | 0.048 | CNTNAP2 |
| GO:0007409 | axonogenesis | 0.048 | 0.048 | NRXN1, LRRC4C, GLI3 |
| GO:0045060 | negative thymic T cell selection | 0.048 | 0.048 | GLI3 |
| GO:0046834 | lipid phosphorylation | 0.048 | 0.048 | DGKB |
| GO:0099612 | protein localization to axon | 0.048 | 0.048 | CNTNAP2 |
| GO:0038130 | ERBB4 signaling pathway | 0.048 | 0.048 | NRG3 |
| GO:0044089 | positive regulation of cellular component biogenesis | 0.049 | 0.049 | NRXN1, CNTNAP2, IL1RAPL1 |
| GO:0048754 | branching morphogenesis of an epithelial tube | 0.049 | 0.049 | CSMD1, GLI3 |
| GO:0097060 | synaptic membrane | <0.0001 | <0.0001 | NRXN1, CNTNAP2, LRRC4C, IL1RAPL1, GABBR2, DGKB |
| GO:0098978 | glutamatergic synapse | 0.001 | 0.001 | NRXN1, CNTNAP2, LRRC4C, IL1RAPL1, NRG3, DGKB |
| GO:0045202 | synapse | 0.001 | 0.001 | NRXN1, CNTNAP2, LRRC4C, IL1RAPL1, GABBR2, NRG3, DGKB, CTNND2 |
| GO:0098685 | Schaffer collateral - CA1 synapse | 0.004 | 0.004 | NRXN1, LRRC4C, DGKB |
| GO:0045211 | postsynaptic membrane | 0.004 | 0.004 | LRRC4C, IL1RAPL1, GABBR2, DGKB |
| GO:0030054 | cell junction | 0.008 | 0.008 | NRXN1, CNTNAP2, LRRC4C, IL1RAPL1, GABBR2, NRG3, DGKB, CTNND2 |
| GO:0098794 | postsynapse | 0.008 | 0.008 | LRRC4C, IL1RAPL1, GABBR2, DGKB, CTNND2 |
| GO:0098590 | plasma membrane region | 0.008 | 0.008 | NRXN1, CNTNAP2, LRRC4C, IL1RAPL1, GABBR2, DGKB |
| GO:1902712 | G protein-coupled GABA receptor complex | 0.028 | 0.028 | GABBR2 |
| GO:0038039 | G protein-coupled receptor heterodimeric complex | 0.038 | 0.038 | GABBR2 |
| GO:1990788 | GLI-SUFU complex | 0.042 | 0.042 | GLI3 |
| GO:0098635 | protein complex involved in cell-cell adhesion | 0.042 | 0.042 | NRXN1 |
| GO:0098982 | GABA-ergic synapse | 0.045 | 0.045 | NRXN1, CNTNAP2 |
| GO:0043005 | neuron projection | 0.047 | 0.047 | NRXN1, CNTNAP2, IL1RAPL1, GABBR2, CTNND2 |
| GO:0036477 | somatodendritic compartment | 0.049 | 0.049 | NRXN1, CNTNAP2, IL1RAPL1, CTNND2 |

**Table S2.** Gene ontology terms (GO.ID) significantly associated with down-regulated genes in the contrast between individuals in the local and foreign song treatments in the first trials. P (FDR) are P-values after false discovery rate correction.

| GO.ID | Description | P | P (FDR) | Genes |
| --- | --- | --- | --- | --- |
| GO:0001675 | acrosome assembly | 0.032 | 0.032 | ZPBP2 |
| GO:0035036 | sperm-egg recognition | 0.032 | 0.032 | ZPBP2 |
| GO:0002638 | negative regulation of immunoglobulin production | 0.032 | 0.032 | ZPBP2 |
| GO:0002701 | negative regulation of production of molecular mediator of immune response | 0.032 | 0.032 | ZPBP2 |
| GO:0007339 | binding of sperm to zona pellucida | 0.032 | 0.032 | ZPBP2 |
| GO:0009988 | cell-cell recognition | 0.035 | 0.035 | ZPBP2 |
| GO:0046466 | membrane lipid catabolic process | 0.035 | 0.035 | ZPBP2 |
| GO:0033363 | secretory granule organization | 0.037 | 0.037 | ZPBP2 |
| GO:0002637 | regulation of immunoglobulin production | 0.037 | 0.037 | ZPBP2 |
| GO:0002698 | negative regulation of immune effector process | 0.037 | 0.037 | ZPBP2 |
| GO:0032922 | circadian regulation of gene expression | 0.037 | 0.037 | ZPBP2 |
| GO:0032989 | cellular anatomical entity morphogenesis | 0.042 | 0.042 | ZPBP2 |
| GO:0010927 | cellular component assembly involved in morphogenesis | 0.042 | 0.042 | ZPBP2 |
| GO:0008037 | cell recognition | 0.042 | 0.042 | ZPBP2 |
| GO:0007338 | single fertilization | 0.042 | 0.042 | ZPBP2 |
| GO:0002377 | immunoglobulin production | 0.042 | 0.042 | ZPBP2 |
| GO:0009566 | fertilization | 0.042 | 0.042 | ZPBP2 |
| GO:0007286 | spermatid development | 0.042 | 0.042 | ZPBP2 |
| GO:0006665 | sphingolipid metabolic process | 0.042 | 0.042 | ZPBP2 |
| GO:0002700 | regulation of production of molecular mediator of immune response | 0.042 | 0.042 | ZPBP2 |
| GO:0048515 | spermatid differentiation | 0.043 | 0.043 | ZPBP2 |
| GO:0002440 | production of molecular mediator of immune response | 0.043 | 0.043 | ZPBP2 |
| GO:0006643 | membrane lipid metabolic process | 0.047 | 0.047 | ZPBP2 |
| GO:0007623 | circadian rhythm | 0.047 | 0.047 | ZPBP2 |
| GO:0002199 | zona pellucida receptor complex | 0.009 | 0.009 | ZPBP2 |

**Table S3.** Gene ontology terms (GO.ID) significantly associated with up-regulated genes in the contrast between individuals in the local song treatment in the first trials and baseline individuals. P (FDR) are P-values after false discovery rate correction.

| GO.ID | Description | P | P (FDR) | Genes |
| --- | --- | --- | --- | --- |
| CORUM:386 | Ubiquitin E3 ligase (UBADC1, RNF123) | 0.004 | 0.004 | UBAC1 |
| CORUM:680 | RANBMP-CD39 complex | 0.004 | 0.004 | ENTPD1 |
| CORUM:7344 | NLGN1-NRXN1 complex | 0.004 | 0.004 | NRXN1 |
| GO:0097090 | presynaptic membrane organization | 0.027 | 0.027 | NRXN1,IL1RAPL1 |
| GO:0097105 | presynaptic membrane assembly | 0.027 | 0.027 | NRXN1,IL1RAPL1 |
| GO:0010308 | acireductone dioxygenase (Ni2+-requiring) activity | 0.049 | 0.049 | ADI1 |
| GO:0010309 | acireductone dioxygenase [iron(II)-requiring] activity | 0.049 | 0.049 | ADI1 |
| GO:0016791 | phosphatase activity | 0.049 | 0.049 | PFKFB3,PPP1R14C,BCKDK,VCAN |
| GO:0047323 | 3-methyl-2-oxobutanoate dehydrogenase (acetyl-transferring) kinase activity | 0.049 | 0.049 | BCKDK |

**Table S4.** Gene ontology terms (GO.ID) significantly associated with down-regulated genes in the contrast between individuals in the local song treatment in the first trials and baseline individuals. P (FDR) are P-values after false discovery rate correction.

| GO.ID | Description | P | P (FDR) | Genes |
| --- | --- | --- | --- | --- |
| CORUM:240 | BRCA1-CTIP-ZBRK1 repressor complex | 0.003 | 0.003 | RBBP8 |
| CORUM:2214 | LMO4-BRCA1-CTIP-LDB1 complex | 0.003 | 0.003 | RBBP8 |
| CORUM:2215 | BRCA1-LMO4-CTIP complex | 0.003 | 0.003 | RBBP8 |
| CORUM:2787 | BRCA1 C complex | 0.003 | 0.003 | RBBP8 |
| CORUM:2819 | BRCA1-CtIP-CtBP complex | 0.003 | 0.003 | RBBP8 |
| CORUM:2923 | SHARP-CtBP1-CtIP complex | 0.003 | 0.003 | RBBP8 |
| CORUM:2930 | SHARP-CtIP-RBP-Jkappa complex | 0.003 | 0.003 | RBBP8 |
| CORUM:2931 | SHARP-CtBP1-CtIP-RBP-Jkappa corepressor complex | 0.003 | 0.003 | RBBP8 |
| CORUM:2815 | BRCA1-BARD1-BACH1-DNA damage complex II | 0.006 | 0.006 | RBBP8 |
| CORUM:5199 | Kinase maturation complex 1 | 0.010 | 0.010 | YWHAH |
| GO:0005272 | sodium channel activity | 0.021 | 0.021 | SHROOM2, YWHAH |
| GO:0031405 | lipoic acid binding | 0.021 | 0.021 | DBT |
| GO:0043754 | dihydrolipoyllysine-residue (2-methylpropanoyl)transferase activity | 0.021 | 0.021 | DBT |
| GO:0003779 | actin binding | 0.031 | 0.031 | SHROOM2, YWHAH, PXK |
| REAC:R-HSA-69473 | G2/M DNA damage checkpoint | 0.035 | 0.035 | YWHAH, RBBP8 |

**Table S5.** Gene ontology terms (GO.ID) significantly associated with down-regulated genes in the contrast between individuals in the foreign song treatment in the first trials and baseline individuals. There were no GO terms significantly associated with up-regulated genes in this comparison. P (FDR) are P-values after false discovery rate correction.

| GO.ID | Description | P | P (FDR) | Genes |
| --- | --- | --- | --- | --- |
| GO:0009081 | branched-chain amino acid metabolic process | 0.015 | 0.015 | DBT |
| GO:0009083 | branched-chain amino acid catabolic process | 0.015 | 0.015 | DBT |
| GO:0009063 | amino acid catabolic process | 0.040 | 0.040 | DBT |
| GO:0046395 | carboxylic acid catabolic process | 0.048 | 0.048 | DBT |
| GO:0016054 | organic acid catabolic process | 0.048 | 0.048 | DBT |
| GO:0044282 | small molecule catabolic process | 0.048 | 0.048 | DBT |
| GO:0006520 | amino acid metabolic process | 0.048 | 0.048 | DBT |
| GO:0160157 | branched-chain alpha-ketoacid dehydrogenase complex | 0.008 | 0.008 | DBT |
| GO:0045239 | tricarboxylic acid cycle heteromeric enzyme complex | 0.009 | 0.009 | DBT |
| GO:0045240 | alpha-ketoacid dehydrogenase complex | 0.009 | 0.009 | DBT |
| GO:0045252 | oxoglutarate dehydrogenase complex | 0.009 | 0.009 | DBT |
| GO:0009295 | nucleoid | 0.014 | 0.014 | DBT |
| GO:0042645 | mitochondrial nucleoid | 0.014 | 0.014 | DBT |
| GO:1990204 | oxidoreductase complex | 0.028 | 0.028 | DBT |
| GO:0031405 | lipoic acid binding | 0.001 | 0.001 | DBT |
| GO:0043754 | dihydrolipoyllysine-residue (2-methylpropanoyl)transferase activity | 0.001 | 0.001 | DBT |
| GO:0047101 | branched-chain alpha-keto acid dehydrogenase activity | 0.003 | 0.003 | DBT |
| GO:0005504 | fatty acid binding | 0.010 | 0.010 | DBT |
| GO:0016620 | oxidoreductase activity, acting on the aldehyde or oxo group of donors, NAD or NADP as acceptor | 0.010 | 0.010 | DBT |
| GO:0016903 | oxidoreductase activity, acting on the aldehyde or oxo group of donors | 0.010 | 0.010 | DBT |
| GO:0033293 | monocarboxylic acid binding | 0.012 | 0.012 | DBT |
| GO:0016407 | acetyltransferase activity | 0.019 | 0.019 | DBT |
| GO:0031406 | carboxylic acid binding | 0.025 | 0.025 | DBT |
| GO:0043177 | organic acid binding | 0.025 | 0.025 | DBT |
| GO:0016747 | acyltransferase activity, transferring groups other than amino-acyl groups | 0.032 | 0.032 | DBT |
| GO:1901681 | sulfur compound binding | 0.035 | 0.035 | DBT |
| GO:0044389 | ubiquitin-like protein ligase binding | 0.040 | 0.040 | DBT |
| GO:0031625 | ubiquitin protein ligase binding | 0.040 | 0.040 | DBT |
| HP:0002179 | Opisthotonus | 0.050 | 0.050 | DBT |
| HP:0001946 | Ketosis | 0.050 | 0.050 | DBT |
| HP:0001993 | Ketoacidosis | 0.050 | 0.050 | DBT |
| KEGG:01210 | 2-Oxocarboxylic acid metabolism | 0.004 | 0.004 | DBT |
| KEGG:00785 | Lipoic acid metabolism | 0.004 | 0.004 | DBT |
| KEGG:00640 | Propanoate metabolism | 0.004 | 0.004 | DBT |
| KEGG:00280 | Valine, leucine and isoleucine degradation | 0.005 | 0.005 | DBT |
| REAC:R-HSA-9857492 | Protein lipoylation | 0.010 | 0.010 | DBT |
| REAC:R-HSA-70895 | Branched-chain amino acid catabolism | 0.010 | 0.010 | DBT |
| REAC:R-HSA-9013407 | RHOH GTPase cycle | 0.011 | 0.011 | DBT |
| REAC:R-HSA-9837999 | Mitochondrial protein degradation | 0.020 | 0.020 | DBT |
| WP:WP4686 | Leucine isoleucine and valine metabolism | 0.003 | 0.003 | DBT |

**Table S6.** Gene ontology terms (GO.ID) significantly associated with the turquoise module in the contrast between individuals in the local and foreign song treatments in the first trials and baseline individuals.

| GO.ID | Description | P | P (FDR) | Genes |
| --- | --- | --- | --- | --- |
| CORUM:571 | p300-CBP-p270 complex | 0.005 | 0.005 | CREBBP, ARID1A, EP300 |
| CORUM:570 | p300-CBP-p270-SWI/SNF complex | 0.023 | 0.023 | CREBBP, ARID1A, EP300 |
| GO:0051254 | positive regulation of RNA metabolic process | 0.006 | 0.006 | ACVR2A, MAP2K7, CSRNP2, MED13, ARID3B, ATRX, MEF2A, ZBTB16, IKBKG, NR4A1, ASXL2, PPRC1, CREBBP, BTG2, KMT2E, TNRC6B, KMT2C, TBX4, KANSL2, SMAD4, MED1, RFX2, HAX1, ASH1L, ARID1A, RUNX1, TET3, EP300. RBPMS, HDAC4, BCAS3, DOT1L, ZMIZ2, NSD1, DYRK1A, PBX1, WNK1, NFIX, TRIM28, ZFP36L2, ARID5A, SERPINF2, TESC, EP400. HCFC1, PMF1, LMX1B, IL4I1, PLCB1, FOXK2, HELZ2, RERE, AIRE, CRTC1, POU2F1, KMT2A, SMARCD3, SOX12, POGZ, E2F4, TNRC6C, TCF7L2, NR4A2, GALR3 |
| GO:1902680 | positive regulation of RNA biosynthetic process | 0.006 | 0.006 | ACVR2A, MAP2K7, CSRNP2, MED13, ARID3B, ATRX, MEF2A, ZBTB16, IKBKG, NR4A1, ASXL2, PPRC1, CREBBP, KMT2E, KMT2C, TBX4, KANSL2, SMAD4, MED1, RFX2, HAX1, ASH1L, ARID1A, RUNX1, TET3, EP300. RBPMS, HDAC4, BCAS3, DOT1L, ZMIZ2, NSD1, DYRK1A, PBX1, WNK1, NFIX, TRIM28, ARID5A, SERPINF2, TESC, EP400. HCFC1, PMF1, LMX1B, IL4I1, PLCB1, FOXK2, HELZ2, RERE, AIRE, CRTC1, POU2F1, KMT2A, SMARCD3, SOX12, POGZ, E2F4, TCF7L2, NR4A2, GALR3 |
| GO:0045935 | positive regulation of nucleobase-containing compound metabolic process | 0.006 | 0.006 | ACVR2A, MAP2K7, CSRNP2, MED13, ARID3B, ATRX, MEF2A, ZBTB16, IKBKG, NR4A1, ASXL2, PPRC1, CREBBP, BTG2, KMT2E, TNRC6B, KMT2C, TBX4, KANSL2, SMAD4, MED1, RFX2, HAX1, ASH1L, PFKFB1, ARID1A, RUNX1, TET3, EP300. WIZ, RBPMS, HDAC4, BCAS3, DOT1L, ZMIZ2, NSD1, DYRK1A, PBX1, WNK1, NFIX, GPD1, TRIM28, ZFP36L2, ARID5A, SERPINF2, TESC, EP400. HCFC1, PMF1, LMX1B, IL4I1, PLCB1, FOXK2, HELZ2, RERE, AIRE, FGFR4, CRTC1, POU2F1, KMT2A, SMARCD3, POLG2, SOX12, POGZ, E2F4, TNRC6C, TCF7L2, NR4A2, GALR3 |
| GO:0045893 | positive regulation of DNA-templated transcription | 0.006 | 0.006 | ACVR2A, MAP2K7, CSRNP2, MED13, ARID3B, ATRX, MEF2A, ZBTB16, IKBKG, NR4A1, ASXL2, PPRC1, CREBBP, KMT2E, KMT2C, TBX4, KANSL2, SMAD4, MED1, RFX2, HAX1, ASH1L, ARID1A, RUNX1, TET3, EP300. RBPMS, HDAC4, BCAS3, DOT1L, ZMIZ2, NSD1, DYRK1A, PBX1, WNK1, NFIX, TRIM28, ARID5A, SERPINF2, TESC, EP400. HCFC1, PMF1, LMX1B, IL4I1, PLCB1, FOXK2, HELZ2, RERE, AIRE, CRTC1, POU2F1, KMT2A, SMARCD3, SOX12, POGZ, E2F4, TCF7L2, NR4A2, GALR3 |
| GO:0051252 | regulation of RNA metabolic process | 0.046 | 0.046 | ACVR2A, MAP2K7, RBM25, CSRNP2, MN1, MED13, ARID3B, ATRX, MEF2A, ZBTB16, NCOR1, IKBKG, NR4A1, ASXL2, PPRC1, CHD5, CREBBP, ZNF687, BTG2, SPEN, SON, KMT2E, TNRC6B, KMT2C, TBX4, KANSL2, SMAD4, MED1, RFX2, HAX1, ASH1L, ARID1A, CEP350. RUNX1, TET3, DACH1, MED13L, EP300. WIZ, RBPMS, HDAC4, BCAS3, MXD1, DOT1L, DACH2, ZMIZ2, NSD1, DYRK1A, PBX1, TENM2, WNK1, NFIX, MPHOSPH8, TRIM28, ZFP36L2, ARID5A, SERPINF2, TESC, EP400. HCFC1, SP2, PMF1, LMX1B, ZNF469, IL4I1, PLCB1, FOXK2, SMAD7, HELZ2, RERE, AIRE, L3MBTL1, CRTC1, POU2F1, ELP3, KMT2A, SMARCD3, CARD9, SOX12, POGZ, E2F4, PURG, TNRC6C, TCF7L2, ZNF618, LARP1, CELF5, NR4A2, GOLGB1, GALR3 |
| GO:2001141 | regulation of RNA biosynthetic process | 0.047 | 0.047 | ACVR2A, MAP2K7, CSRNP2, MN1, MED13, ARID3B, ATRX, MEF2A, ZBTB16, NCOR1, IKBKG, NR4A1, ASXL2, PPRC1, CHD5, CREBBP, ZNF687, BTG2, SPEN, KMT2E, KMT2C, TBX4, KANSL2, SMAD4, MED1, RFX2, HAX1, ASH1L, ARID1A, CEP350. RUNX1, TET3, DACH1, MED13L, EP300. WIZ, RBPMS, HDAC4, BCAS3, MXD1, DOT1L, DACH2, ZMIZ2, NSD1, DYRK1A, PBX1, TENM2, WNK1, NFIX, MPHOSPH8, TRIM28, ARID5A, SERPINF2, TESC, EP400. HCFC1, SP2, PMF1, LMX1B, ZNF469, IL4I1, PLCB1, FOXK2, SMAD7, HELZ2, RERE, AIRE, L3MBTL1, CRTC1, POU2F1, ELP3, KMT2A, SMARCD3, CARD9, SOX12, POGZ, E2F4, PURG, TCF7L2, ZNF618, NR4A2, GOLGB1, GALR3 |
| GO:0019219 | regulation of nucleobase-containing compound metabolic process | 0.047 | 0.047 | ACVR2A, MAP2K7, RBM25, CSRNP2, MN1, MED13, ARID3B, ATRX, MEF2A, ZBTB16, NCOR1, IKBKG, NR4A1, ASXL2, PPRC1, CHD5, CREBBP, ZNF687, BTG2, SPEN, SON, KMT2E, TNRC6B, KMT2C, TBX4, KANSL2, SMAD4, MED1, RFX2, HAX1, ASH1L, PFKFB1, ARID1A, CEP350. RUNX1, TET3, DACH1, MED13L, EP300. WIZ, RBPMS, HDAC4, BCAS3, MXD1, DOT1L, DACH2, ZMIZ2, NSD1, DYRK1A, PBX1, TENM2, WNK1, NFIX, GPD1, MPHOSPH8, TRIM28, ZFP36L2, ANKRD17, ARID5A, SERPINF2, TESC, EP400. CDC6, HCFC1, SP2, PMF1, LMX1B, ZNF469, IL4I1, PLCB1, FOXK2, SMAD7, HELZ2, RERE, AIRE, L3MBTL1, FGFR4, CRTC1, POU2F1, ELP3, KMT2A, SMARCD3, POLG2, CARD9, SOX12, POGZ, E2F4, PURG, TNRC6C, TCF7L2, ZNF618, LARP1, CELF5, NR4A2, GOLGB1, GALR3 |
| GO:0016070 | RNA metabolic process | 0.047 | 0.047 | ACVR2A, MAP2K7, RBM25, CSRNP2, ZC3H13, MN1, MED13, ARID3B, ATRX, MEF2A, ZBTB16, SNRPA, NCOR1, IKBKG, NR4A1, ASXL2, PPRC1, CHD5, CREBBP, ZNF687, BTG2, SPEN, SON, KMT2E, TNRC6B, KMT2C, TBX4, KANSL2, SMAD4, MED1, RFX2, HAX1, AHNAK2, SNRNP200. ASH1L, ARID1A, CEP350. RUNX1, SNRNP35, TET3, DACH1, MED13L, EP300. WIZ, RBPMS, HDAC4, BCAS3, ZNF638, MXD1, DOT1L, HIPK3, DACH2, ZMIZ2, NSD1, PPAN, DYRK1A, PBX1, TENM2, WNK1, NFIX, MPHOSPH8, WDR6, TRIM28, ZFP36L2, CLK2, ARID5A, PDCD11, SERPINF2, TESC, EP400. HCFC1, SP2, GTF3C2, PMF1, LMX1B, PRPF8, ZNF469, IL4I1, PLCB1, FOXK2, SMAD7, HELZ2, RERE, AIRE, L3MBTL1, CRTC1, ADARB2, POU2F1, ELP3, KMT2A, NF1, SMARCD3, XAB2, CARD9, SOX12, POGZ, E2F4, VARS2, PURG, TNRC6C, TCF7L2, ZNF618, LARP1, CELF5, RBMS2, NR4A2, GOLGB1, GALR3 |
| GO:0045944 | positive regulation of transcription by RNA polymerase II | 0.050 | 0.050 | ACVR2A, CSRNP2, MED13, ARID3B, ATRX, MEF2A, ZBTB16, IKBKG, NR4A1, ASXL2, PPRC1, CREBBP, KMT2C, SMAD4, MED1, RFX2, HAX1, ASH1L, RUNX1, TET3, EP300. HDAC4, BCAS3, DOT1L, ZMIZ2, PBX1, NFIX, ARID5A, SERPINF2, HCFC1, LMX1B, IL4I1, FOXK2, HELZ2, AIRE, CRTC1, POU2F1, KMT2A, SOX12, POGZ, E2F4, TCF7L2, NR4A2, GALR3 |
| GO:0006355 | regulation of DNA-templated transcription | 0.050 | 0.050 | ACVR2A, MAP2K7, CSRNP2, MN1, MED13, ARID3B, ATRX, MEF2A, ZBTB16, NCOR1, IKBKG, NR4A1, ASXL2, PPRC1, CHD5, CREBBP, ZNF687, BTG2, SPEN, KMT2E, KMT2C, TBX4, KANSL2, SMAD4, MED1, RFX2, HAX1, ASH1L, ARID1A, CEP350. RUNX1, TET3, DACH1, MED13L, EP300. WIZ, RBPMS, HDAC4, BCAS3, MXD1, DOT1L, DACH2, ZMIZ2, NSD1, DYRK1A, PBX1, TENM2, WNK1, NFIX, TRIM28, ARID5A, SERPINF2, TESC, EP400. HCFC1, SP2, PMF1, LMX1B, ZNF469, IL4I1, PLCB1, FOXK2, SMAD7, HELZ2, RERE, AIRE, L3MBTL1, CRTC1, POU2F1, ELP3, KMT2A, SMARCD3, CARD9, SOX12, POGZ, E2F4, PURG, TCF7L2, ZNF618, NR4A2, GOLGB1, GALR3 |
| GO:0006351 | DNA-templated transcription | 0.050 | 0.050 | ACVR2A, MAP2K7, CSRNP2, MN1, MED13, ARID3B, ATRX, MEF2A, ZBTB16, NCOR1, IKBKG, NR4A1, ASXL2, PPRC1, CHD5, CREBBP, ZNF687, BTG2, SPEN, KMT2E, KMT2C, TBX4, KANSL2, SMAD4, MED1, RFX2, HAX1, ASH1L, ARID1A, CEP350. RUNX1, TET3, DACH1, MED13L, EP300. WIZ, RBPMS, HDAC4, BCAS3, MXD1, DOT1L, HIPK3, DACH2, ZMIZ2, NSD1, DYRK1A, PBX1, TENM2, WNK1, NFIX, TRIM28, ARID5A, SERPINF2, TESC, EP400. HCFC1, SP2, GTF3C2, PMF1, LMX1B, ZNF469, IL4I1, PLCB1, FOXK2, SMAD7, HELZ2, RERE, AIRE, L3MBTL1, CRTC1, POU2F1, ELP3, KMT2A, SMARCD3, XAB2, CARD9, SOX12, POGZ, E2F4, PURG, TCF7L2, ZNF618, NR4A2, GOLGB1, GALR3 |
| GO:0141187 | nucleic acid biosynthetic process | 0.050 | 0.050 | ACVR2A, MAP2K7, RBM25, CSRNP2, ZC3H13, MN1, MED13, ARID3B, ATRX, MEF2A, ZBTB16, SNRPA, NCOR1, IKBKG, NR4A1, ASXL2, PPRC1, CHD5, CREBBP, ZNF687, BTG2, SPEN, SON, KMT2E, KMT2C, TBX4, KANSL2, SMAD4, MED1, RFX2, HAX1, AHNAK2, SNRNP200. ASH1L, HSP90AB1, ARID1A, CEP350. RUNX1, SNRNP35, TET3, DACH1, MED13L, EP300. WIZ, RBPMS, HDAC4, BCAS3, ZNF638, MXD1, DOT1L, HIPK3, DACH2, ZMIZ2, NSD1, PPAN, DYRK1A, PBX1, TENM2, WNK1, NFIX, MPHOSPH8, WDR6, TRIM28, CLK2, ARID5A, PDCD11, SERPINF2, TESC, EP400. HCFC1, SP2, GTF3C2, PMF1, LMX1B, PRPF8, ZNF469, IL4I1, PLCB1, FOXK2, SMAD7, HELZ2, RERE, AIRE, L3MBTL1, FGFR4, CRTC1, ADARB2, POU2F1, ELP3, KMT2A, SMARCD3, XAB2, POLG2, CARD9, SOX12, POGZ, E2F4, PURG, TCF7L2, ZNF618, CELF5, RBMS2, NR4A2, GOLGB1, GALR3 |
| GO:0090304 | nucleic acid metabolic process | 0.050 | 0.050 | ACVR2A, MAP2K7, RBM25, CSRNP2, ZC3H13, MN1, MED13, ARID3B, ATRX, MEF2A, ZBTB16, SNRPA, NCOR1, IKBKG, NR4A1, ASXL2, PPRC1, CHD5, CREBBP, HERC2, ZNF687, BTG2, SPEN, SON, KMT2E, TNRC6B, KMT2C, TBX4, KANSL2, SMAD4, MED1, RFX2, HAX1, AHNAK2, SNRNP200. ASH1L, HSP90AB1, ARID1A, CEP350. RUNX1, SNRNP35, TET3, DACH1, MED13L, EP300. WIZ, RBPMS, HDAC4, BCAS3, ZNF638, MXD1, DOT1L, HIPK3, FANCG, DACH2, ZMIZ2, NSD1, PPAN, DYRK1A, PBX1, TENM2, WNK1, NFIX, MPHOSPH8, WDR6, TRIM28, ZFP36L2, CLK2, ANKRD17, ARID5A, PDCD11, SERPINF2, TESC, EP400. CDC6, HCFC1, SP2, GTF3C2, PMF1, LMX1B, PRPF8, ZNF469, IL4I1, PLCB1, FOXK2, SMAD7, HELZ2, ZSWIM7, RERE, AIRE, L3MBTL1, TEX11, FGFR4, GINS4, CRTC1, ADARB2, POU2F1, ELP3, KMT2A, FANCD2, NF1, SMARCD3, XAB2, POLG2, CARD9, SOX12, POGZ, E2F4, VARS2, PURG, TNRC6C, TCF7L2, ZNF618, LARP1, CELF5, RBMS2, NR4A2, GOLGB1, GALR3 |
| GO:0005667 | transcription regulator complex | 0.013 | 0.013 | MEF2A, ZBTB16, NCOR1, NR4A1, CHD5, CREBBP, SPEN, SMAD4, MED1, HAX1, RUNX1, DACH1, EP300. HDAC4, MXD1, DACH2, PBX1, TRIM28, ARID5A, GTF3C2, PMF1, SMAD7, POU2F1, E2F4, TCF7L2, NR4A2 |
| GO:0003712 | transcription coregulator activity | 0.007 | 0.007 | MED13, NCOR1, PPRC1, CREBBP, BTG2, SPEN, KMT2E, KMT2C, MED1, ARID1A, MED13L, EP300. RBPMS, HDAC4, DOT1L, NSD1, DYRK1A, TRIM28, ARID5A, HCFC1, PMF1, SMAD7, RERE, SMARCD3, TCERG1L |
| GO:0060090 | molecular adaptor activity | 0.007 | 0.007 | MED13, NCOR1, IKBKG, PPRC1, CHD5, CREBBP, BTG2, SPEN, KMT2E, ZMYND10. KMT2C, MED1, HAX1, ARID1A, GAB3, MED13L, EP300. MAGI2, RBPMS, HDAC4, DOT1L, NSD1, DYRK1A, WNK1, BBS4, MPHOSPH8, TRIM28, ARID5A, NAPB, HCFC1, RAB3A, DAB1, PMF1, SMAD7, RERE, ITSN2, SURF6, SMARCD3, CARD9, PCNT, UBE2Z, SNAP47, ANKRD13D, TCERG1L |
| GO:0030674 | protein-macromolecule adaptor activity | 0.007 | 0.007 | MED13, NCOR1, IKBKG, PPRC1, CHD5, CREBBP, BTG2, SPEN, KMT2E, KMT2C, MED1, HAX1, ARID1A, GAB3, MED13L, EP300. MAGI2, RBPMS, HDAC4, DOT1L, NSD1, DYRK1A, WNK1, BBS4, MPHOSPH8, TRIM28, ARID5A, NAPB, HCFC1, RAB3A, DAB1, PMF1, SMAD7, RERE, SURF6, SMARCD3, CARD9, SNAP47, ANKRD13D, TCERG1L |
| GO:0061629 | RNA polymerase II-specific DNA-binding transcription factor binding | 0.007 | 0.007 | MED13, MEF2A, NCOR1, NR4A1, CREBBP, SPEN, SMAD4, MED1, ARID1A, EP300. HDAC4, BCAS3, DOT1L, NSD1, BBS4, ARID5A, PDCD11, SMARCD3, TCF7L2, NR4A2 |
| GO:0016922 | nuclear receptor binding | 0.007 | 0.007 | MED13, NCOR1, NR4A1, MED1, ARID1A, EP300. BCAS3, NSD1, ARID5A, SMARCD3, TCF7L2, NR4A2 |
| GO:0003682 | chromatin binding | 0.007 | 0.007 | ATRX, MEF2A, ASXL2, CHD5, CREBBP, SMAD4, MED1, ASH1L, ARID1A, EP300. HDAC4, BCAS3, NSD1, MPHOSPH8, TRIM28, TNRC18, ANKRD17, ARID5A, EP400. HCFC1, RERE, AIRE, L3MBTL1, MBD5, KMT2A, SMARCD3, E2F4, TCF7L2 |
| GO:0008134 | transcription factor binding | 0.007 | 0.007 | MED13, MEF2A, ZBTB16, NCOR1, NR4A1, PPRC1, CREBBP, SPEN, SMAD4, MED1, ARID1A, RUNX1, EP300. WIZ, HDAC4, BCAS3, DOT1L, NSD1, PBX1, BBS4, ARID5A, PDCD11, HCFC1, CRTC1, SMARCD3, TCF7L2, ZNF618, NR4A2 |
| GO:0140297 | DNA-binding transcription factor binding | 0.007 | 0.007 | MED13, MEF2A, NCOR1, NR4A1, CREBBP, SPEN, SMAD4, MED1, ARID1A, RUNX1, EP300. HDAC4, BCAS3, DOT1L, NSD1, PBX1, BBS4, ARID5A, PDCD11, HCFC1, CRTC1, SMARCD3, TCF7L2, NR4A2 |
| GO:0140938 | histone H3 methyltransferase activity | 0.007 | 0.007 | KMT2C, ASH1L, DOT1L, NSD1, KMT2A, SETD1B |
| GO:0140110 | transcription regulator activity | 0.007 | 0.007 | CSRNP2, MED13, MEF2A, ZBTB16, NCOR1, NR4A1, PPRC1, CREBBP, ZNF687, BTG2, SPEN, KMT2E, KMT2C, TBX4, SMAD4, MED1, RFX2, ARID1A, RUNX1, DACH1, MED13L, EP300. WIZ, RBPMS, HDAC4, MXD1, DOT1L, DACH2, NSD1, DYRK1A, PBX1, NFIX, TRIM28, ARID5A, HCFC1, SP2, PMF1, LMX1B, ZNF469, FOXK2, SMAD7, RERE, POU2F1, SMARCD3, SOX12, E2F4, PURG, TCF7L2, ZNF618, NR4A2, TCERG1L |
| GO:0140945 | histone H3K4 monomethyltransferase activity | 0.012 | 0.012 | KMT2C, KMT2A, SETD1B |
| GO:0003713 | transcription coactivator activity | 0.013 | 0.013 | MED13, CREBBP, KMT2E, KMT2C, MED1, ARID1A, EP300. RBPMS, DOT1L, DYRK1A, TRIM28, HCFC1, PMF1, RERE, SMARCD3 |
| GO:0001221 | transcription coregulator binding | 0.014 | 0.014 | ZBTB16, CREBBP, SMAD4, MED1, RUNX1, EP300. WIZ, PBX1, SMARCD3, ZNF618 |
| GO:0042800 | histone H3K4 methyltransferase activity | 0.014 | 0.014 | KMT2C, ASH1L, KMT2A, SETD1B |
| GO:0046966 | nuclear thyroid hormone receptor binding | 0.016 | 0.016 | MED13, NCOR1, MED1, NSD1, ARID5A |
| GO:0140999 | histone H3K4 trimethyltransferase activity | 0.016 | 0.016 | KMT2C, KMT2A, SETD1B |
| GO:0003677 | DNA binding | 0.018 | 0.018 | CSRNP2, ARID3B, ATRX, MEF2A, ZBTB16, SNRPA, NCOR1, NR4A1, ASXL2, CHD5, CREBBP, ZNF687, SPEN, SON, KMT2C, TBX4, SMAD4, MED1, RFX2, ASH1L, ARID1A, CEP350. RUNX1, TET3, DACH1, EP300. WIZ, HDAC4, ZNF638, MXD1, DOT1L, FANCG, DACH2, NSD1, PBX1, NFIX, TRIM28, ARID5A, EP400. SP2, GTF3C2, LMX1B, ZNF469, FOXK2, HELZ2, RERE, AIRE, POU2F1, MBD5, SURF6, KMT2A, POLG2, SOX12, POGZ, E2F4, PURG, TCF7L2, ZNF618, NR4A2, GOLGB1 |
| GO:0015211 | purine nucleoside transmembrane transporter activity | 0.027 | 0.027 | SLC29A1, SLC25A26 |
| GO:0120300 | peptide lactyltransferase (CoA-dependent) activity | 0.027 | 0.027 | CREBBP, EP300 |
| GO:0044017 | histone H3K27 acetyltransferase activity | 0.027 | 0.027 | CREBBP, EP300 |
| GO:0003676 | nucleic acid binding | 0.032 | 0.032 | RBM25, CSRNP2, ZC3H13, ARID3B, ATRX, MEF2A, ZBTB16, EIF4G3, SNRPA, NCOR1, NR4A1, ASXL2, PPRC1, CHD5, CREBBP, ZNF687, SPEN, MACF1, SON, TNRC6B, KMT2C, TBX4, SMAD4, MED1, PRRC2C, RFX2, UBAP2L, SNRNP200. ANKHD1, ASH1L, HSP90AB1, TNS1, ARID1A, CEP350. RUNX1, SNRNP35, TET3, DACH1, EP300. WIZ, RBPMS, HDAC4, ZNF638, RAVER1, MXD1, DOT1L, FANCG, DACH2, NSD1, PPAN, PBX1, NFIX, WDR6, TRIM28, EIF5B, ZFP36L2, ANKRD17, ARID5A, PDCD11, EP400. SP2, GTF3C2, LMX1B, DYNC1H1, PRPF8, ZNF469, FOXK2, HELZ2, RERE, AIRE, MAP4, GPATCH8, ADARB2, POU2F1, ELP3, MBD5, SURF6, KMT2A, POLG2, SOX12, POGZ, E2F4, SETD1B, PURG, TNRC6C, TCF7L2, ZNF618, LARP1, CELF5, RBMS2, NR4A2, GOLGB1 |
| GO:0043167 | ion binding | 0.039 | 0.039 | BSN, ACVR2A, EFCAB5, GK5, MAP2K7, ZC3H13, ATRX, ZBTB16, UBR4, IKBKG, NR4A1, ASXL2, CHD5, CREBBP, HERC2, ZNF687, MACF1, PAK4, KMT2E, ZMYND10. KMT2C, DOCK7, SMAD4, SNRNP200. GALK1, PDE6B, DTNB, MYO15A, HERC1, ZC3HC1, ASH1L, HSP90AB1, PFKFB1, TNS1, RUNX1, TET3, MAPK13, ITGB4, ARHGEF12, EP300. ATP4A, WIZ, AK7, HDAC4, CAMK2B, FRAS1, ZNF638, RAB5B, TNS2, SBF1, HIPK3, ESYT1, SYVN1, NUDT8, D2HGDH, ZMIZ2, GSR, NSD1, CALN1, CUTA, DYRK1A, ABCC2, CARNS1, ZMYND19, TENM2, WNK1, REM1, TRIM28, HCN3, EIF5B, ZFP36L2, CLK2, CELSR3, RAPGEFL1, TESC, CPXM1, EP400. CDC6, RAB3A, SP2, LMX1B, SLIT1, SIL1, ACTG2, CABP4, HSPG2, DYNC1H1, GRIN3B, ZNF469, PLCB1, NCS1, RASSF1, FOXK2, SMAD7, HELZ2, ZSWIM7, RERE, AIRE, NECAB2, L3MBTL1, TRIM54, DNAH17, FGFR4, ITSN2, RHO, RPS6KB2, PLEK2, GPATCH8, ADARB2, PTK7, STAB1, ELP3, RNFT2, KMT2A, NF1, CARD9, POGZ, MDN1, SYT12, UBE2Z, VARS2, KIF13B, HAO1, DOCK1, TAT, PHYHD1, ZNF618, PDE10A, GCGR, DNER, LPP, NR4A2, CPNE7, MYLK2, RNF213, KIF17 |
| GO:0036094 | small molecule binding | 0.041 | 0.041 | BSN, ACVR2A, EFCAB5, GK5, MAP2K7, ZC3H13, ATRX, ZBTB16, UBR4, IKBKG, NR4A1, ASXL2, CHD5, CREBBP, HERC2, ZNF687, MACF1, PAK4, KMT2E, ZMYND10. KMT2C, DOCK7, SMAD4, SNRNP200. GALK1, PDE6B, DTNB, MYO15A, HERC1, ZC3HC1, ASH1L, HSP90AB1, PFKFB1, TNS1, RUNX1, TET3, MAPK13, ITGB4, ARHGEF12, EP300. ATP4A, WIZ, AK7, HDAC4, CAMK2B, FRAS1, ZNF638, RAB5B, GGCX, TNS2, SBF1, HIPK3, ESYT1, SYVN1, NUDT8, D2HGDH, ZMIZ2, GSR, NSD1, CALN1, CUTA, DYRK1A, ABCC2, CARNS1, ZMYND19, TENM2, WNK1, REM1, GPD1, TRIM28, HCN3, EIF5B, ZFP36L2, CLK2, CELSR3, RAPGEFL1, TESC, CPXM1, EP400. CDC6, RAB3A, SP2, LMX1B, SLIT1, SIL1, ACTG2, CABP4, HSPG2, DYNC1H1, GRIN3B, ZNF469, PLCB1, NCS1, RASSF1, FOXK2, SMAD7, HELZ2, ZSWIM7, RERE, AIRE, NECAB2, L3MBTL1, TRIM54, DNAH17, FGFR4, ITSN2, RHO, RPS6KB2, PLEK2, GPATCH8, ADARB2, PTK7, STAB1, ELP3, RNFT2, KMT2A, NF1, CARD9, POGZ, MDN1, SYT12, CHRNA2, UBE2Z, VARS2, KIF13B, HAO1, DOCK1, TAT, PHYHD1, ZNF618, PDE10A, GCGR, DNER, LPP, NR4A2, CPNE7, MYLK2, RNF213, KIF17 |
| GO:0042974 | nuclear retinoic acid receptor binding | 0.041 | 0.041 | MED1, NSD1, ARID5A, NR4A2 |
| GO:0042054 | histone methyltransferase activity | 0.041 | 0.041 | KMT2C, ASH1L, DOT1L, NSD1, KMT2A, SETD1B |
| GO:0016279 | protein-lysine N-methyltransferase activity | 0.044 | 0.044 | KMT2C, ASH1L, DOT1L, NSD1, KMT2A, SETD1B |
| GO:0016278 | lysine N-methyltransferase activity | 0.047 | 0.047 | KMT2C, ASH1L, DOT1L, NSD1, KMT2A, SETD1B |
| HP:0000752 | Hyperactivity | <0.0001 | <0.0001 | MED13, ATRX, IKBKG, CHD5, CREBBP, HERC2, SPEN, TNRC6B, SMAD4, UBAP2L, ASH1L, ARID1A, TET3, ADSL, DISP1, MED13L, EP300. ANKRD11, HDAC4, APC2, SZT2, SLC6A8, NSD1, DYRK1A, NFIX, ANKRD17, CABP4, HSPG2, RERE, DPP6, MBD5, KMT2A, FANCD2, NF1, PCNT, POGZ, CHRNA2, KCNT1 |
| HP:0000729 | Autistic behavior | <0.0001 | <0.0001 | MED13, ATRX, CHD5, CREBBP, HERC2, SPEN, SON, TNRC6B, KMT2C, DOCK7, SMAD4, UBAP2L, ASH1L, ARID1A, TET3, ADSL, MED13L, EP300. ANKRD11, NPRL2, HDAC4, APC2, CAMK2B, SZT2, SLC6A8, NSD1, NLGN3, DYRK1A, NFIX, BBS4, ANKRD17, HSPG2, RERE, MBD5, KMT2A, NF1, HECTD4, POGZ, SETD1B, NR4A2 |
| HP:0025783 | Diagnostic behavioral phenotype | <0.0001 | <0.0001 | MED13, ATRX, CHD5, CREBBP, HERC2, SPEN, SON, TNRC6B, KMT2C, DOCK7, SMAD4, UBAP2L, ASH1L, ARID1A, TET3, ADSL, MED13L, EP300. ANKRD11, NPRL2, HDAC4, APC2, CAMK2B, SZT2, SLC6A8, NSD1, NLGN3, DYRK1A, NFIX, BBS4, ANKRD17, HSPG2, RERE, MBD5, KMT2A, NF1, POLG2, HECTD4, POGZ, SETD1B, NR4A2 |
| HP:0000734 | Disinhibition | <0.0001 | <0.0001 | MED13, ATRX, IKBKG, CHD5, CREBBP, HERC2, SPEN, KMT2E, TNRC6B, SMAD4, UBAP2L, ASH1L, ARID1A, DCTN1, TET3, ADSL, DISP1, MED13L, EP300. ANKRD11, HDAC4, APC2, CAMK2B, SZT2, SLC6A8, NSD1, NLGN3, DYRK1A, NFIX, ANKRD17, CABP4, HSPG2, RERE, DPP6, MBD5, KMT2A, FANCD2, NF1, PCNT, HECTD4, POGZ, CHRNA2, KCNT1 |
| HP:5200263 | Abnormally increased volition | <0.0001 | <0.0001 | MED13, ATRX, IKBKG, CHD5, CREBBP, HERC2, SPEN, KMT2E, TNRC6B, SMAD4, UBAP2L, ASH1L, ARID1A, DCTN1, TET3, ADSL, DISP1, MED13L, EP300. ANKRD11, HDAC4, APC2, CAMK2B, SZT2, SLC6A8, NSD1, NLGN3, DYRK1A, NFIX, ANKRD17, CABP4, HSPG2, RERE, DPP6, MBD5, KMT2A, FANCD2, NF1, PCNT, HECTD4, POGZ, CHRNA2, KCNT1 |
| HP:0000540 | Hypermetropia | <0.0001 | <0.0001 | CREBBP, HERC2, SPEN, SON, KMT2C, SMAD4, UBAP2L, PDE6B, ASH1L, MED13L, EP300. APC2, SLC6A8, NSD1, DYRK1A, PBX1, NFIX, CABP4, HSPG2, RERE, RHO, MBD5, PCNT, POGZ |
| HP:5200241 | Recurrent maladaptive behavior | <0.0001 | <0.0001 | MN1, MED13, ATRX, ZBTB16, IKBKG, CHD5, CREBBP, HERC2, SPEN, MACF1, KMT2E, TNRC6B, KMT2C, DOCK7, SMAD4, UBAP2L, ASH1L, ARID1A, DCTN1, RUNX1, TET3, ADSL, DISP1, MED13L, EP300. ANKRD11, HDAC4, APC2, CAMK2B, SZT2, SLC6A8, NSD1, NLGN3, DYRK1A, NFIX, ANKRD17, NAPB, CABP4, HSPG2, RERE, DPP6, MBD5, KMT2A, FANCD2, NF1, PCNT, HECTD4, POGZ, CHRNA2, KCNT1, NR4A2 |
| HP:0000733 | Motor stereotypy | <0.0001 | <0.0001 | ATRX, CREBBP, HERC2, SPEN, MACF1, DOCK7, UBAP2L, ASH1L, EP300. HDAC4, CAMK2B, SLC6A8, NLGN3, DYRK1A, NAPB, CABP4, HSPG2, RERE, MBD5, KMT2A, POGZ, CHRNA2, KCNT1 |
| HP:0000534 | Abnormal eyebrow morphology | <0.0001 | <0.0001 | MN1, MED13, ASXL2, CHD5, CREBBP, SPEN, SON, KMT2C, DOCK7, TBX4, SMAD4, UBAP2L, HERC1, ASH1L, CEP41, ARID1A, MED13L, EP300. ANKRD11, HDAC4, BCAS3, FRAS1, SZT2, NUP188, NSD1, NFIX, BBS4, LMX1B, HSPG2, SV2A, RERE, MBD5, KMT2A, NF1, PCNT, HECTD4, POGZ, SETD1B |
| HP:0000708 | Atypical behavior | <0.0001 | <0.0001 | MN1, MED13, ATRX, ZBTB16, IKBKG, CHD5, CREBBP, HERC2, SPEN, MACF1, SON, KMT2E, TNRC6B, KMT2C, DOCK7, TBX4, SMAD4, UBAP2L, SNRNP200. PDE6B, HERC1, ASH1L, ARID1A, DCTN1, RUNX1, TET3, ADSL, DISP1, ITGB4, MED13L, EP300. ANKRD11, NPRL2, HDAC4, APC2, CAMK2B, SZT2, SLC6A8, NSD1, NLGN3, DYRK1A, NFIX, BBS4, ANKRD17, NAPB, LMX1B, CABP4, HSPG2, PRPF8, RERE, DPP6, AIRE, RHO, MBD5, KMT2A, FANCD2, NF1, POLG2, PCNT, HECTD4, POGZ, CHRNA2, SETD1B, KCNT1, TAT, NR4A2 |
| HP:0031432 | Restricted or repetitive behaviors or interests | <0.0001 | <0.0001 | ATRX, CREBBP, HERC2, SPEN, MACF1, DOCK7, UBAP2L, ASH1L, EP300. HDAC4, CAMK2B, SLC6A8, NLGN3, DYRK1A, NAPB, CABP4, HSPG2, RERE, MBD5, KMT2A, POGZ, CHRNA2, KCNT1 |
| HP:0000736 | Short attention span | <0.0001 | <0.0001 | MED13, IKBKG, CHD5, CREBBP, HERC2, SPEN, TNRC6B, UBAP2L, TET3, DISP1, EP300. ANKRD11, HDAC4, APC2, SZT2, SLC6A8, NSD1, DYRK1A, BBS4, ANKRD17, CABP4, HSPG2, RERE, MBD5, KMT2A, FANCD2, NF1, PCNT, POGZ, CHRNA2, KCNT1 |
| HP:0000549 | Abnormal conjugate eye movement | <0.0001 | <0.0001 | MN1, MED13, ATRX, IKBKG, CREBBP, HERC2, SPEN, MACF1, SON, TBX4, SMAD4, UBAP2L, PDE6B, ASH1L, CEP41, ARID1A, DCTN1, TET3, ADSL, DISP1, MED13L, EP300. ANKRD11, APC2, BCAS3, CAMK2B, GGCX, SBF1, FANCG, SLC6A8, NSD1, DYRK1A, PBX1, NFIX, BBS4, DHODH, ANKRD17, SLC37A4, DAB1, LMX1B, SIL1, CABP4, HSPG2, RERE, DPP6, WDR81, RHO, MBD5, KMT2A, FANCD2, NF1, CTSK, POGZ, CLDN19 |
| HP:5200044 | Reduced attention regulation | <0.0001 | <0.0001 | MED13, IKBKG, CHD5, CREBBP, HERC2, SPEN, TNRC6B, UBAP2L, TET3, DISP1, EP300. ANKRD11, HDAC4, APC2, SZT2, SLC6A8, NSD1, DYRK1A, BBS4, ANKRD17, CABP4, HSPG2, RERE, MBD5, KMT2A, FANCD2, NF1, PCNT, POGZ, CHRNA2, KCNT1 |
| HP:0010720 | Abnormal hair pattern | <0.0001 | <0.0001 | IKBKG, CREBBP, HERC2, SPEN, DOCK7, UBAP2L, ASH1L, ARID1A, ITGB4, MED13L, EP300. ANKRD11, HDAC4, APC2, FRAS1, NSD1, NFIX, ANKRD17, LMX1B, HSPG2, RERE, AIRE, MBD5, KMT2A, NF1 |
| HP:0000486 | Strabismus | <0.0001 | <0.0001 | MN1, MED13, ATRX, IKBKG, CREBBP, HERC2, SPEN, MACF1, SON, TBX4, SMAD4, UBAP2L, PDE6B, ASH1L, CEP41, ARID1A, TET3, ADSL, DISP1, MED13L, EP300. ANKRD11, APC2, BCAS3, CAMK2B, GGCX, SBF1, FANCG, SLC6A8, NSD1, DYRK1A, PBX1, NFIX, BBS4, DHODH, ANKRD17, SLC37A4, LMX1B, SIL1, CABP4, HSPG2, RERE, DPP6, WDR81, RHO, MBD5, KMT2A, FANCD2, NF1, CTSK, POGZ, CLDN19 |
| HP:0002683 | Abnormal calvaria morphology | <0.0001 | <0.0001 | MN1, ATRX, IKBKG, CHD5, CREBBP, HERC2, SPEN, SON, KMT2E, KMT2C, TBX4, SMAD4, HERC1, ASH1L, CEP41, TET3, ADSL, ITGB4, MED13L, EP300. ANKRD11, HDAC4, APC2, FRAS1, NUP188, D2HGDH, FANCG, NSD1, DYRK1A, NFIX, ANKRD17, CDC6, HCFC1, LMX1B, HSPG2, DYNC1H1, EXOC6B, RERE, MBD5, KMT2A, FANCD2, CTSK, PCNT, HECTD4, POGZ, SETD1B |
| HP:0000539 | Abnormality of refraction | <0.0001 | <0.0001 | ATRX, CREBBP, HERC2, SPEN, SON, KMT2C, SMAD4, UBAP2L, PDE6B, HERC1, ASH1L, ARID1A, MED13L, EP300. ANKRD11, HDAC4, APC2, CAMK2B, FANCG, SLC6A8, NSD1, DYRK1A, PBX1, NFIX, BBS4, LMX1B, CABP4, HSPG2, ZNF469, RERE, RHO, MBD5, FANCD2, CTSK, PCNT, POGZ, CLDN19 |
| HP:0007018 | Attention deficit hyperactivity disorder | <0.0001 | <0.0001 | MED13, IKBKG, CHD5, HERC2, SPEN, TNRC6B, UBAP2L, TET3, DISP1, ANKRD11, HDAC4, APC2, SZT2, SLC6A8, NSD1, DYRK1A, ANKRD17, CABP4, HSPG2, RERE, FANCD2, NF1, PCNT, POGZ, CHRNA2, KCNT1 |
| HP:5200230 | Maladaptive fear-related cognitions | <0.0001 | <0.0001 | ATRX, CREBBP, KMT2E, UBAP2L, ASH1L, DCTN1, TET3, DISP1, EP300. APC2, NSD1, DYRK1A, NFIX, BBS4, CABP4, HSPG2, MBD5, KMT2A, NF1, POLG2, HECTD4, POGZ, CHRNA2, SETD1B, KCNT1, PDE10A, NR4A2 |
| HP:0000483 | Astigmatism | <0.0001 | <0.0001 | UBAP2L, ASH1L, ANKRD11, HDAC4, APC2, CAMK2B, FANCG, NSD1, DYRK1A, PBX1, NFIX, BBS4, LMX1B, RERE, MBD5, FANCD2, POGZ, CLDN19 |
| HP:0008551 | Microtia | <0.0001 | <0.0001 | ATRX, CHD5, SPEN, TBX4, SMAD4, UBAP2L, ITGB4, FRAS1, PBX1, NFIX, DHODH, CDC6, HSPG2, RERE, MBD5, PCNT, HECTD4 |
| HP:0000739 | Anxiety | <0.0001 | <0.0001 | ATRX, CREBBP, KMT2E, UBAP2L, ASH1L, DCTN1, TET3, DISP1, EP300. APC2, NSD1, DYRK1A, NFIX, BBS4, CABP4, HSPG2, MBD5, KMT2A, NF1, POLG2, HECTD4, POGZ, CHRNA2, SETD1B, KCNT1, PDE10A, NR4A2 |
| HP:0002463 | Language impairment | <0.0001 | <0.0001 | MN1, MED13, ASXL2, CHD5, CREBBP, HERC2, SPEN, MACF1, KMT2E, ZMYND10. TNRC6B, KMT2C, DOCK7, TBX4, SMAD4, UBAP2L, HERC1, ASH1L, ARID1A, DCTN1, TET3, ADSL, DISP1, MED13L, EP300. ANKRD11, HDAC4, APC2, BCAS3, CAMK2B, SZT2, SLC6A8, NSD1, NLGN3, DYRK1A, PBX1, NFIX, BBS4, ANKRD17, LMX1B, HSPG2, DYNC1H1, RERE, WDR81, MBD5, KMT2A, NF1, POGZ, SETD1B, KCNT1, NR4A2 |
| HP:0100691 | Abnormality of the curvature of the cornea | <0.0001 | <0.0001 | UBAP2L, SNRNP200. PDE6B, ASH1L, ANKRD11, HDAC4, APC2, CAMK2B, FANCG, NSD1, DYRK1A, PBX1, NFIX, BBS4, LMX1B, PRPF8, ZNF469, RERE, RHO, MBD5, FANCD2, POGZ, CLDN19 |
| HP:5200423 | Abnormal experience of reality | <0.0001 | <0.0001 | ATRX, CREBBP, HERC2, SON, KMT2E, SMAD4, UBAP2L, ASH1L, DCTN1, TET3, DISP1, EP300. NPRL2, APC2, NSD1, DYRK1A, NFIX, BBS4, CABP4, HSPG2, MBD5, KMT2A, NF1, POLG2, HECTD4, POGZ, CHRNA2, SETD1B, KCNT1, PDE10A, NR4A2 |
| HP:0002553 | Highly arched eyebrow | <0.0001 | <0.0001 | MN1, ASXL2, CREBBP, SPEN, TBX4, SMAD4, CEP41, EP300. ANKRD11, HDAC4, SZT2, NUP188, NFIX, LMX1B, SV2A, MBD5, KMT2A, POGZ |
| HP:0000718 | Aggressive behavior | <0.0001 | <0.0001 | CHD5, CREBBP, HERC2, SPEN, KMT2E, UBAP2L, ARID1A, ADSL, MED13L, EP300. HDAC4, APC2, CAMK2B, SLC6A8, NSD1, DYRK1A, MBD5, KMT2A, HECTD4, POGZ, KCNT1 |
| HP:0000750 | Delayed speech and language development | <0.0001 | <0.0001 | MN1, MED13, ASXL2, CHD5, CREBBP, HERC2, SPEN, MACF1, KMT2E, ZMYND10. TNRC6B, KMT2C, DOCK7, TBX4, UBAP2L, HERC1, ASH1L, ARID1A, TET3, ADSL, DISP1, MED13L, EP300. ANKRD11, HDAC4, APC2, BCAS3, CAMK2B, SZT2, SLC6A8, NSD1, NLGN3, DYRK1A, PBX1, NFIX, BBS4, ANKRD17, LMX1B, HSPG2, DYNC1H1, RERE, WDR81, MBD5, KMT2A, NF1, POGZ, SETD1B, KCNT1, NR4A2 |
| HP:0011361 | Congenital abnormal hair pattern | <0.0001 | <0.0001 | CREBBP, HERC2, SPEN, DOCK7, UBAP2L, ASH1L, ARID1A, MED13L, EP300. ANKRD11, HDAC4, APC2, FRAS1, NSD1, NFIX, ANKRD17, LMX1B, HSPG2, RERE, MBD5, KMT2A, NF1 |
| HP:0008771 | Aplasia/Hypoplasia of the ear | <0.0001 | <0.0001 | ATRX, CHD5, SPEN, TBX4, SMAD4, UBAP2L, ITGB4, FRAS1, PBX1, NFIX, DHODH, CDC6, HSPG2, RERE, GREB1L, MBD5, PCNT, HECTD4 |
| HP:0001250 | Seizure | <0.0001 | <0.0001 | MN1, MED13, ATRX, IKBKG, ASXL2, CHD5, CREBBP, HERC2, SPEN, MACF1, SON, KMT2E, TNRC6B, KMT2C, DOCK7, SMAD4, UBAP2L, HAX1, GALK1, HERC1, ASH1L, CEP41, ARID1A, TET3, NUP214, ADSL, DISP1, EP300. ANKRD11, NPRL2, HDAC4, APC2, BCAS3, CAMK2B, FRAS1, SZT2, D2HGDH, SLC6A8, NSD1, NLGN3, DYRK1A, NFIX, BBS4, ANKRD17, NAPB, HCFC1, SLC37A4, LMX1B, ST3GAL3, CABP4, HSPG2, DYNC1H1, PLCB1, PTPN23, SV2A, RERE, AIRE, WDR81, MBD5, KMT2A, NF1, POLG2, PCNT, HECTD4, POGZ, CHRNA2, SETD1B, VARS2, KCNT1, TAT, PDE10A, NR4A2, RNF213 |
| HP:5200401 | Abnormal judgment | <0.0001 | <0.0001 | ATRX, CREBBP, KMT2E, UBAP2L, ASH1L, DCTN1, TET3, DISP1, EP300. APC2, NSD1, DYRK1A, NFIX, BBS4, CABP4, HSPG2, MBD5, KMT2A, NF1, POLG2, HECTD4, POGZ, CHRNA2, SETD1B, KCNT1, PDE10A, NR4A2 |
| HP:0000717 | Autism | <0.0001 | <0.0001 | ATRX, CREBBP, HERC2, SPEN, DOCK7, SMAD4, ADSL, MED13L, EP300. ANKRD11, HDAC4, SZT2, NLGN3, DYRK1A, BBS4, HSPG2, RERE, KMT2A, HECTD4 |
| HP:0008772 | Aplasia/Hypoplasia of the external ear | 0.001 | 0.001 | ATRX, CHD5, SPEN, TBX4, SMAD4, UBAP2L, ITGB4, FRAS1, PBX1, NFIX, DHODH, CDC6, HSPG2, RERE, MBD5, PCNT, HECTD4 |
| HP:0009553 | Abnormality of the hairline | 0.001 | 0.001 | CREBBP, SPEN, DOCK7, UBAP2L, ASH1L, ARID1A, MED13L, EP300. ANKRD11, APC2, FRAS1, NSD1, NFIX, ANKRD17, LMX1B, HSPG2, RERE, MBD5, KMT2A, NF1 |
| HP:0000240 | Abnormality of skull size | 0.001 | 0.001 | ATRX, IKBKG, ASXL2, CREBBP, SPEN, MACF1, SON, KMT2E, TNRC6B, KMT2C, TBX4, SMAD4, GALK1, HERC1, ASH1L, ARID1A, TET3, NUP214, ADSL, DISP1, EP300. ANKRD11, HDAC4, APC2, BCAS3, CAMK2B, FRAS1, SZT2, SBF1, NUP188, D2HGDH, FANCG, SLC6A8, NSD1, DYRK1A, NFIX, ANKRD17, NAPB, CDC6, HCFC1, LMX1B, SIL1, ACTG2, HSPG2, DYNC1H1, ZNF469, PLCB1, PTPN23, ZSWIM7, SV2A, RERE, DPP6, MBD5, KMT2A, FANCD2, NF1, PCNT, HECTD4, POGZ, VARS2, KCNT1, TAT |
| HP:0000252 | Microcephaly | 0.001 | 0.001 | ATRX, IKBKG, CREBBP, SPEN, MACF1, TNRC6B, KMT2C, TBX4, SMAD4, GALK1, ASH1L, ARID1A, TET3, NUP214, ADSL, DISP1, EP300. ANKRD11, HDAC4, BCAS3, CAMK2B, FRAS1, SZT2, SBF1, NUP188, FANCG, SLC6A8, NSD1, DYRK1A, NFIX, ANKRD17, NAPB, CDC6, HCFC1, LMX1B, SIL1, ACTG2, HSPG2, DYNC1H1, PLCB1, PTPN23, ZSWIM7, SV2A, RERE, DPP6, MBD5, KMT2A, FANCD2, NF1, PCNT, POGZ, VARS2, KCNT1, TAT |
| HP:0025732 | Abnormal social development | 0.001 | 0.001 | ATRX, CREBBP, HERC1, DCTN1, TET3, ADSL, MED13L, EP300. APC2, SLC6A8, DYRK1A, MBD5, NF1, POGZ |
| HP:0100716 | Self-injurious behavior | 0.001 | 0.001 | ATRX, CHD5, CREBBP, HERC2, SPEN, KMT2E, KMT2C, UBAP2L, ADSL, EP300. HDAC4, SLC6A8, NFIX, HSPG2, RERE, MBD5, POGZ |
| HP:0007364 | Aplasia/Hypoplasia of the cerebrum | 0.001 | 0.001 | ATRX, IKBKG, CREBBP, SPEN, MACF1, SON, TNRC6B, KMT2C, DOCK7, TBX4, SMAD4, GALK1, ASH1L, CEP41, ARID1A, TET3, NUP214, ADSL, DISP1, EP300. ANKRD11, HDAC4, APC2, BCAS3, CAMK2B, FRAS1, SZT2, SBF1, NUP188, FANCG, SLC6A8, NSD1, DYRK1A, NFIX, ANKRD17, NAPB, CDC6, HCFC1, LMX1B, SIL1, ACTG2, HSPG2, DYNC1H1, PLCB1, PTPN23, ZSWIM7, SV2A, RERE, DPP6, WDR81, MBD5, KMT2A, FANCD2, NF1, POLG2, PCNT, HECTD4, POGZ, VARS2, KCNT1, TAT |
| HP:0003272 | Abnormal hip bone morphology | 0.001 | 0.001 | CREBBP, HERC2, SPEN, MACF1, TBX4, SMAD4, TET3, DISP1, EP300. ANKRD11, HDAC4, APC2, FRAS1, FANCG, NSD1, WNK1, NFIX, DHODH, PHLDB1, LMX1B, SIL1, HSPG2, EXOC6B, ZNF469, RERE, SERPINF1, MBD5, FANCD2, CTSK, PCNT |
| HP:0000826 | Precocious puberty | 0.001 | 0.001 | CREBBP, HERC2, SPEN, TNRC6B, KMT2C, SMAD4, DISP1, EP300. DYRK1A, NFIX, PLCB1, NF1, PCNT, KCNT1 |
| HP:0040195 | Decreased head circumference | 0.001 | 0.001 | ATRX, IKBKG, CREBBP, SPEN, MACF1, TNRC6B, KMT2C, TBX4, SMAD4, GALK1, ASH1L, ARID1A, TET3, NUP214, ADSL, DISP1, EP300. ANKRD11, HDAC4, BCAS3, CAMK2B, FRAS1, SZT2, SBF1, NUP188, FANCG, SLC6A8, NSD1, DYRK1A, NFIX, ANKRD17, NAPB, CDC6, HCFC1, LMX1B, SIL1, ACTG2, HSPG2, DYNC1H1, PLCB1, PTPN23, ZSWIM7, SV2A, RERE, DPP6, MBD5, KMT2A, FANCD2, NF1, PCNT, POGZ, VARS2, KCNT1, TAT |
| HP:0000574 | Thick eyebrow | 0.001 | 0.001 | MN1, CREBBP, SPEN, KMT2C, DOCK7, SMAD4, ARID1A, EP300. ANKRD11, BCAS3, NFIX, MBD5, KMT2A, SETD1B |
| HP:0025780 | Abnormal volitional state | 0.001 | 0.001 | MED13, ATRX, IKBKG, CHD5, CREBBP, HERC2, SPEN, KMT2E, TNRC6B, SMAD4, UBAP2L, ASH1L, ARID1A, DCTN1, TET3, ADSL, DISP1, MED13L, EP300. ANKRD11, HDAC4, APC2, CAMK2B, SZT2, SLC6A8, NSD1, NLGN3, DYRK1A, PBX1, NFIX, BBS4, ANKRD17, CABP4, HSPG2, RERE, DPP6, MBD5, KMT2A, FANCD2, NF1, POLG2, PCNT, HECTD4, POGZ, CHRNA2, KCNT1, NR4A2 |
| HP:0100000 | Early onset of sexual maturation | 0.001 | 0.001 | CREBBP, HERC2, SPEN, TNRC6B, KMT2C, SMAD4, DISP1, EP300. DYRK1A, NFIX, PLCB1, NF1, PCNT, KCNT1 |
| HP:0002236 | Frontal upsweep of hair | 0.001 | 0.001 | CREBBP, HERC2, UBAP2L, MED13L, EP300. HDAC4 |
| HP:0031703 | Abnormal ear morphology | 0.001 | 0.001 | MN1, ATRX, IKBKG, ASXL2, CHD5, CREBBP, SPEN, MACF1, SON, ZMYND10. TNRC6B, KMT2C, DOCK7, TBX4, SMAD4, UBAP2L, HAX1, SNRNP200. PDE6B, HERC1, ASH1L, CEP41, ARID1A, TET3, ADSL, ITGB4, MED13L, EP300. ANKRD11, HDAC4, APC2, FRAS1, NUP188, FANCG, SLC6A8, NSD1, DYRK1A, PBX1, NFIX, BBS4, DHODH, CDC6, SLC37A4, LMX1B, ACTG2, HSPG2, PRPF8, ZNF469, RERE, DPP6, AIRE, GREB1L, RHO, MBD5, KMT2A, FANCD2, NF1, CTSK, PCNT, HECTD4, POGZ, NR4A2 |
| HP:0100037 | Abnormality of the scalp hair | 0.001 | 0.001 | ATRX, CREBBP, SPEN, DOCK7, UBAP2L, ASH1L, ARID1A, ITGB4, MED13L, EP300. ANKRD11, HDAC4, APC2, FRAS1, NSD1, NFIX, ANKRD17, LMX1B, HSPG2, RERE, MBD5, KMT2A, NF1, PCNT |
| HP:0010674 | Abnormal curvature of the vertebral column | 0.001 | 0.001 | ATRX, IKBKG, ASXL2, CREBBP, HERC2, SPEN, SON, KMT2E, KMT2C, TBX4, SMAD4, UBAP2L, HERC1, ASH1L, CEP41, ARID1A, DISP1, EP300. ANKRD11, HDAC4, APC2, SBF1, FANCG, NSD1, DYRK1A, WNK1, NFIX, ANKRD17, SLC37A4, LMX1B, SIL1, HSPG2, EXOC6B, ZNF469, PLCB1, RERE, DPP6, WDR81, PHKA1, MBD5, KMT2A, FANCD2, NF1, CTSK, PCNT, HECTD4, POGZ, KCNT1, NR4A2 |
| HP:0008050 | Abnormality of the palpebral fissures | 0.001 | 0.001 | MN1, MED13, ATRX, CHD5, CREBBP, HERC2, SPEN, SON, KMT2E, TNRC6B, KMT2C, SMAD4, UBAP2L, HERC1, MED13L, EP300. ANKRD11, HDAC4, APC2, SZT2, FANCG, NSD1, DYRK1A, NFIX, BBS4, DHODH, ANKRD17, LMX1B, HSPG2, DYNC1H1, RERE, KMT2A, FANCD2, NF1, PCNT, HECTD4, POGZ, SETD1B, NR4A2 |
| HP:0001999 | Abnormal facial shape | 0.001 | 0.001 | MN1, ATRX, ASXL2, CREBBP, SPEN, MACF1, SON, KMT2E, TNRC6B, KMT2C, UBAP2L, HERC1, CEP41, ARID1A, TET3, ADSL, MED13L, EP300. ANKRD11, HDAC4, APC2, BCAS3, FRAS1, FANCG, SLC6A8, NSD1, DYRK1A, PBX1, NFIX, ANKRD17, CDC6, SLC37A4, LMX1B, ACTG2, HSPG2, RERE, WDR81, MBD5, KMT2A, FANCD2, NF1, PCNT, HECTD4, POGZ, SETD1B, NR4A2 |

**Table S7.** Gene ontology terms (GO.ID) significantly associated with up-regulated genes in the contrast between individuals in the foreign song treatment in the paired trials. There were no down-regulated differentially expressed genes in this comparison. P (FDR) are P-values after false discovery rate correction.

| GO.ID | Description | P | P (FDR) | Genes |
| --- | --- | --- | --- | --- |
| GO:0007156 | homophilic cell adhesion via plasma membrane adhesion molecules | 0.035 | 0.035 | CDH18, PCDH7, CADM2 |
| REAC:R-HSA-418990 | Adherens junctions interactions | 0.046 | 0.046 | CDH18, CADM2 |

**Table S8.** Gene ontology terms (GO.ID) significantly associated with the darkorange2 module in the contrast between individuals in the local and foreign song treatments in the paired trials.

| GO.ID | Description | P | P (FDR) | Genes |
| --- | --- | --- | --- | --- |
| GO:0034330 | cell junction organization | <0.0001 | <0.0001 | CDH18, PTPRF, CNTN5, GRID1, GRID2, CDH2, WNT7A, ELMO1, DNM3, CLSTN2, TENM4, IL1RAPL1, CTTNBP2, CD2AP, NTRK2, SHISA6, NBEA, CSMD2, ROBO2, LRTM2, CDH12, SDK1, DAB1, RAB3A, PKHD1, PTPRO, PRRT1, KIF1A, CTNND2, PTPRT, CNKSR2, CDH13, ANK2, MARVELD3, GRIN2B, BCR, TANC2, ROR2, GABRB3, RELN, CACNB2, DLG5, DBN1, DISC1, SYNGAP1, NRXN1, DNER, ZDHHC15, RTN4R, PARD3, PLXND1, CTNNA2, CDKL5, TBX5, CAMK1, EPHA7, MTSS1, BSN, DST, EPHB2, EPHB1, GABRA2, EFNB2, TNS1 |
| GO:0050808 | synapse organization | <0.0001 | <0.0001 | PTPRF, CNTN5, GRID1, GRID2, CDH2, WNT7A, ELMO1, DNM3, CLSTN2, TENM4, IL1RAPL1, CTTNBP2, CD2AP, NTRK2, SHISA6, NBEA, CSMD2, ROBO2, LRTM2, SDK1, DAB1, RAB3A, PTPRO, PRRT1, KIF1A, CTNND2, PTPRT, CNKSR2, GRIN2B, TANC2, ROR2, GABRB3, RELN, CACNB2, DLG5, DBN1, DISC1, SYNGAP1, NRXN1, DNER, ZDHHC15, RTN4R, PLXND1, CTNNA2, CDKL5, CAMK1, EPHA7, BSN, EPHB2, EPHB1, GABRA2 |
| GO:0007399 | nervous system development | <0.0001 | <0.0001 | PTPRF, FOXP2, MYO6, CNTN5, GRID2, FGF14, PLXDC1, CDH2, CAMK1D, MDGA2, WNT7A, NAV3, TENM1, ELMO1, DNM3, NOVA1, ZMIZ1, SH2B2, MACROD2, USH2A, CLSTN2, TENM4, ULK4, RBFOX1, IL1RAPL1, SOX21, CD2AP, NTRK2, CSMD2, AFF2, ROBO2, LRTM2, MAP1S, ZFP36L1, SDK1, DAB1, MSI2, DMD, MAGI2, KIFAP3, TP63, RAB3A, PREX2, PTPRG, PAX5, ASTN2, PTPRO, CRIM1, CNTN1, KIF1A, CTNND2, NCAM1, ARHGEF10. ANK2, DAAM2, GRIN2B, BCR, TANC2, SLIT3, PKD1, IRX1, ROR2, SEMA6A, CHL1, TRPV1, NOTCH2, GABRB3, GABRB1, RELN, DLG5, DBN1, ZNF521, GAL3ST1, SDCCAG8, HSPG2, EPHA6, DISC1, SYNGAP1, NRXN1, AKT3, PROX1, DNER, ZDHHC15, BCL11A, ACAP3, ZFHX3, BCL11B, RTN4R, PARD3, CEP85L, CHD7, SPTBN1, PLXND1, MXRA8, CTNNA2, VAX1, SKI, CCDC88A, RAPGEFL1, SIM2, CDKL5, AUTS2, CAMK1, EPHA7, APBA1, FZD3, CELSR1, PPP3CA, BMPR1B, ERBB3, BSN, CPLX2, MNAT1, PBX1, SCARF1, EPHB2, EPHB1, ATL1, GABRA2, NHEJ1, MAP2, SAMD14, EFNB2, NKX2-2, ADORA2A, KCNIP2, RAPGEF5 |
| GO:0048731 | system development | <0.0001 | <0.0001 | PTPRF, FOXP2, MYO6, CNTN5, CSMD1, GRID2, FGF14, PLXDC1, CDH2, CAMK1D, MDGA2, WNT7A, NAV3, TENM1, ELMO1, DNM3, NOVA1, ZMIZ1, SH2B2, MACROD2, USH2A, CLSTN2, DCN, TENM4, ULK4, RBFOX1, IL1RAPL1, SOX21, CD2AP, NTRK2, E2F7, CSMD2, TCOF1, AFF2, ROBO2, LRTM2, MAP1S, ZFP36L1, SDK1, TCF7L2, DAB1, MSI2, DMD, CPE, MAGI2, ADAMTS2, A2M, ABI3BP, ARHGAP24, KIFAP3, TP63, WT1, RAB3A, PREX2, PTPRG, GLUL, CLMP, PAX5, PKHD1, PPARGC1B, ASTN2, PTPRO, CRIM1, CNTN1, KIF1A, CTNND2, NCAM1, CBLB, MAP2K6, CDH13, ARHGEF10. ANK2, LAMA4, CADM1, DAAM2, GRIN2B, BCR, EMCN, TANC2, SLIT3, PKD1, GCNT4, IRX1, ROR2, SEMA6A, CHST11, CHL1, TRPV1, NOTCH2, GABRB3, GABRB1, RELN, COL1A1, DLG5, DBN1, ZNF521, GAL3ST1, SDCCAG8, HSPG2, EPHA6, DISC1, SYNGAP1, NRXN1, AKT3, PROX1, DNER, ZDHHC15, BCL11A, CPAMD8, ACAP3, RORC, ZFHX3, BCL11B, RTN4R, PARD3, CEP85L, CHD7, SPTBN1, PLXND1, FOXL1, MXRA8, FBN2, CTNNA2, VAX1, SKI, CCDC88A, TRPS1, RAPGEFL1, SIM2, CDKL5, PDGFD, LDB3, AUTS2, TBX5, NEBL, CAMK1, EPHA7, APBA1, FZD3, CELSR1, PPP3CA, CACNA1G, BMPR1B, MTSS1, ADAM8, ERBB3, BSN, CPLX2, MNAT1, PBX1, COL4A2, SCARF1, EPHB2, EPHB1, ANGPT2, ADAM12, ATL1, GABRA2, NHEJ1, MAP2, SAMD14, TSPAN18, EFNB2, NKX2-2, ADORA2A, KCNIP2, ZFPM2, NSDHL, RAPGEF5, STAB2 |
| GO:0007155 | cell adhesion | <0.0001 | <0.0001 | CDH18, PTPRF, CNTN5, PCDH7, GRID2, CDH2, TENM1, PARVG, ZMIZ1, CLSTN2, TENM4, IL1RAPL1, CD2AP, ROBO2, CDH12, ZFP36L1, SDK1, DAB1, DMD, ABI3BP, EDIL3, KIFAP3, SPON1, PKHD1, ASTN2, PTPRO, CNTN1, CTNND2, PTPRT, CTNNAL1, JAK3, NCAM1, CBLB, CDH13, LAMA4, CADM1, BCR, EMCN, PKD1, NID2, PODXL2, SEMA6A, CHL1, RELN, COL1A1, DLG5, FCHO1, HSPG2, DISC1, NRXN1, ZFHX3, PARD3, PLXND1, MXRA8, DOCK5, CTNNA2, CTNNA3, LRRN2, LCK, UTRN, EPHA7, APBA1, CELSR1, PPP3CA, MTSS1, ADAM8, ERBB3, SCARF1, DST, EPHB2, MYH10. EPHB1, ANGPT2, ADAM12, EFNB2, ADORA2A, STAB2 |
| GO:0050803 | regulation of synapse structure or activity | <0.0001 | <0.0001 | GRID1, GRID2, CDH2, WNT7A, ELMO1, DNM3, CLSTN2, IL1RAPL1, CTTNBP2, NTRK2, ROBO2, LRTM2, DAB1, PTPRO, KIF1A, PTPRT, GRIN2B, TANC2, ROR2, RELN, DLG5, DBN1, DISC1, SYNGAP1, NRXN1, ZDHHC15, RTN4R, CTNNA2, CDKL5, CAMK1, EPHA7, EPHB2, EPHB1 |
| GO:0050807 | regulation of synapse organization | <0.0001 | <0.0001 | GRID1, GRID2, CDH2, WNT7A, ELMO1, DNM3, CLSTN2, IL1RAPL1, CTTNBP2, NTRK2, ROBO2, LRTM2, DAB1, PTPRO, KIF1A, PTPRT, GRIN2B, TANC2, ROR2, RELN, DLG5, DBN1, DISC1, NRXN1, ZDHHC15, RTN4R, CTNNA2, CDKL5, CAMK1, EPHA7, EPHB2, EPHB1 |
| GO:0007275 | multicellular organism development | <0.0001 | <0.0001 | PTPRF, FOXP2, MYO6, CNTN5, CSMD1, GRID2, FGF14, PLXDC1, CDH2, CAMK1D, MDGA2, WNT7A, NAV3, TENM1, ELMO1, DNM3, NOVA1, ZMIZ1, SH2B2, MACROD2, USH2A, CLSTN2, DCN, TENM4, ULK4, RBFOX1, IL1RAPL1, SOX21, CD2AP, RAD51B, NTRK2, E2F7, CSMD2, DACH1, TCOF1, AFF2, ROBO2, LRTM2, MAP1S, ZFP36L1, SDK1, TCF7L2, DAB1, MSI2, DMD, CPE, TTC39C, MAGI2, ADAMTS2, A2M, ABI3BP, ARHGAP24, KIFAP3, TP63, ADCYAP1R1, WT1, RAB3A, PREX2, PTPRG, GLUL, CLMP, PAX5, PKHD1, PPARGC1B, ASTN2, PTPRO, CRIM1, CNTN1, KIF1A, CTNND2, JAK3, NCAM1, CBLB, MAP2K6, CDH13, ARHGEF10. ANK2, ASPH, LAMA4, CADM1, DAAM2, GRIN2B, BCR, EMCN, TANC2, SLIT3, PKD1, GCNT4, IRX1, STOX2, ROR2, SEMA6A, CHST11, CHL1, TRPV1, NOTCH2, GABRB3, GABRB1, RELN, TCF7, COL1A1, DLG5, DBN1, ZNF521, GAL3ST1, SDCCAG8, HSPG2, EPHA6, DISC1, SYNGAP1, NRXN1, AKT3, PROX1, DNER, ZDHHC15, BCL11A, CPAMD8, ACAP3, RORC, ZFHX3, BCL11B, RTN4R, PARD3, CEP85L, CHD7, SPTBN1, PLXND1, FOXL1, MXRA8, FBN2, CTNNA2, VAX1, SKI, CCDC88A, TRPS1, RAPGEFL1, SIM2, CDKL5, PDGFD, LDB3, AUTS2, TBX5, NEBL, CAMK1, EPHA7, APBA1, FZD3, CELSR1, PPP3CA, CACNA1G, BMPR1B, MTSS1, ADAM8, ERBB3, BSN, CPLX2, MNAT1, PBX1, COL4A2, SCARF1, EPHB2, EPHB1, ANGPT2, ADAM12, ATL1, GABRA2, NHEJ1, MAP2, SAMD14, TSPAN18, EFNB2, NKX2-2, ADORA2A, KCNIP2, GAL, ZFPM2, SH3PXD2A, NSDHL, RAPGEF5, STAB2, MLLT3 |
| GO:0048667 | cell morphogenesis involved in neuron differentiation | <0.0001 | <0.0001 | CNTN5, WNT7A, DNM3, IL1RAPL1, CD2AP, NTRK2, ROBO2, LRTM2, MAP1S, DAB1, RAB3A, PREX2, PTPRO, CNTN1, KIF1A, CTNND2, NCAM1, TANC2, SLIT3, SEMA6A, CHL1, NOTCH2, RELN, DBN1, EPHA6, DISC1, SYNGAP1, NRXN1, ZDHHC15, BCL11A, BCL11B, RTN4R, PARD3, PLXND1, CTNNA2, VAX1, CDKL5, AUTS2, EPHA7, FZD3, PPP3CA, BMPR1B, EPHB2, EPHB1, ATL1, MAP2, EFNB2 |
| GO:0048699 | generation of neurons | <0.0001 | <0.0001 | PTPRF, MYO6, CNTN5, GRID2, CAMK1D, MDGA2, WNT7A, TENM1, DNM3, ZMIZ1, USH2A, TENM4, ULK4, IL1RAPL1, SOX21, CD2AP, NTRK2, ROBO2, LRTM2, MAP1S, SDK1, DAB1, DMD, MAGI2, KIFAP3, RAB3A, PREX2, PTPRG, ASTN2, PTPRO, CNTN1, KIF1A, CTNND2, NCAM1, TANC2, SLIT3, IRX1, ROR2, SEMA6A, CHL1, NOTCH2, GABRB1, RELN, DLG5, DBN1, ZNF521, SDCCAG8, EPHA6, DISC1, SYNGAP1, NRXN1, PROX1, DNER, ZDHHC15, BCL11A, ACAP3, ZFHX3, BCL11B, RTN4R, PARD3, CEP85L, PLXND1, CTNNA2, VAX1, CCDC88A, CDKL5, AUTS2, CAMK1, EPHA7, FZD3, CELSR1, PPP3CA, BMPR1B, PBX1, SCARF1, EPHB2, EPHB1, ATL1, MAP2, SAMD14, EFNB2, NKX2-2, ADORA2A, KCNIP2 |
| GO:0022008 | neurogenesis | <0.0001 | <0.0001 | PTPRF, MYO6, CNTN5, GRID2, CDH2, CAMK1D, MDGA2, WNT7A, NAV3, TENM1, DNM3, ZMIZ1, USH2A, TENM4, ULK4, IL1RAPL1, SOX21, CD2AP, NTRK2, ROBO2, LRTM2, MAP1S, SDK1, DAB1, DMD, MAGI2, KIFAP3, RAB3A, PREX2, PTPRG, ASTN2, PTPRO, CNTN1, KIF1A, CTNND2, NCAM1, ARHGEF10. DAAM2, TANC2, SLIT3, IRX1, ROR2, SEMA6A, CHL1, NOTCH2, GABRB1, RELN, DLG5, DBN1, ZNF521, SDCCAG8, EPHA6, DISC1, SYNGAP1, NRXN1, PROX1, DNER, ZDHHC15, BCL11A, ACAP3, ZFHX3, BCL11B, RTN4R, PARD3, CEP85L, CHD7, PLXND1, MXRA8, CTNNA2, VAX1, SKI, CCDC88A, CDKL5, AUTS2, CAMK1, EPHA7, FZD3, CELSR1, PPP3CA, BMPR1B, ERBB3, PBX1, SCARF1, EPHB2, EPHB1, ATL1, MAP2, SAMD14, EFNB2, NKX2-2, ADORA2A, KCNIP2 |
| GO:0099173 | postsynapse organization | <0.0001 | <0.0001 | GRID1, GRID2, CDH2, WNT7A, ELMO1, DNM3, IL1RAPL1, SHISA6, CSMD2, KIF1A, CTNND2, CNKSR2, GRIN2B, TANC2, ROR2, RELN, DLG5, DBN1, DISC1, SYNGAP1, NRXN1, ZDHHC15, RTN4R, CDKL5, EPHA7, EPHB2, EPHB1 |
| GO:0031175 | neuron projection development | <0.0001 | <0.0001 | PTPRF, CNTN5, GRID2, CAMK1D, WNT7A, DNM3, ULK4, IL1RAPL1, CD2AP, NTRK2, ROBO2, LRTM2, MAP1S, SDK1, DAB1, DMD, MAGI2, RAB3A, PREX2, PTPRG, PTPRO, CNTN1, KIF1A, CTNND2, NCAM1, TANC2, SLIT3, ROR2, SEMA6A, CHL1, NOTCH2, RELN, DLG5, DBN1, EPHA6, DISC1, SYNGAP1, NRXN1, ZDHHC15, BCL11A, ACAP3, BCL11B, RTN4R, PARD3, PLXND1, CTNNA2, VAX1, CCDC88A, CDKL5, AUTS2, CAMK1, EPHA7, FZD3, PPP3CA, BMPR1B, SCARF1, EPHB2, EPHB1, ATL1, MAP2, SAMD14, EFNB2, ADORA2A |
| GO:0048666 | neuron development | <0.0001 | <0.0001 | PTPRF, CNTN5, GRID2, CAMK1D, WNT7A, TENM1, DNM3, ZMIZ1, TENM4, ULK4, IL1RAPL1, CD2AP, NTRK2, ROBO2, LRTM2, MAP1S, SDK1, DAB1, DMD, MAGI2, RAB3A, PREX2, PTPRG, PTPRO, CNTN1, KIF1A, CTNND2, NCAM1, TANC2, SLIT3, ROR2, SEMA6A, CHL1, NOTCH2, GABRB1, RELN, DLG5, DBN1, EPHA6, DISC1, SYNGAP1, NRXN1, ZDHHC15, BCL11A, ACAP3, BCL11B, RTN4R, PARD3, PLXND1, CTNNA2, VAX1, CCDC88A, CDKL5, AUTS2, CAMK1, EPHA7, FZD3, PPP3CA, BMPR1B, PBX1, SCARF1, EPHB2, EPHB1, ATL1, MAP2, SAMD14, EFNB2, ADORA2A, KCNIP2 |
| GO:0048812 | neuron projection morphogenesis | <0.0001 | <0.0001 | CNTN5, WNT7A, DNM3, IL1RAPL1, CD2AP, NTRK2, ROBO2, LRTM2, MAP1S, DAB1, RAB3A, PREX2, PTPRO, CNTN1, KIF1A, CTNND2, NCAM1, TANC2, SLIT3, SEMA6A, CHL1, NOTCH2, RELN, DBN1, EPHA6, DISC1, SYNGAP1, NRXN1, ZDHHC15, BCL11A, BCL11B, RTN4R, PARD3, PLXND1, CTNNA2, VAX1, CDKL5, AUTS2, EPHA7, FZD3, PPP3CA, BMPR1B, EPHB2, EPHB1, ATL1, MAP2, EFNB2, ADORA2A |
| GO:0010975 | regulation of neuron projection development | <0.0001 | <0.0001 | PTPRF, GRID2, CAMK1D, WNT7A, DNM3, ULK4, IL1RAPL1, NTRK2, ROBO2, DAB1, DMD, MAGI2, PTPRG, PTPRO, CNTN1, KIF1A, TANC2, ROR2, RELN, DBN1, DISC1, SYNGAP1, ZDHHC15, BCL11A, ACAP3, RTN4R, PLXND1, CTNNA2, CCDC88A, CDKL5, CAMK1, EPHA7, PPP3CA, SCARF1, EPHB2, MAP2, EFNB2 |
| GO:0016358 | dendrite development | <0.0001 | <0.0001 | CAMK1D, WNT7A, DNM3, IL1RAPL1, MAP1S, SDK1, DAB1, PREX2, KIF1A, CTNND2, TANC2, RELN, DLG5, DBN1, DISC1, SYNGAP1, ZDHHC15, BCL11A, CTNNA2, CDKL5, CAMK1, PPP3CA, SCARF1, EPHB2, EPHB1, MAP2 |
| GO:0120039 | plasma membrane bounded cell projection morphogenesis | <0.0001 | <0.0001 | CNTN5, WNT7A, DNM3, IL1RAPL1, CD2AP, NTRK2, ROBO2, LRTM2, MAP1S, DAB1, RAB3A, PREX2, PTPRO, CNTN1, KIF1A, CTNND2, NCAM1, TANC2, SLIT3, SEMA6A, CHL1, NOTCH2, RELN, DBN1, EPHA6, DISC1, SYNGAP1, NRXN1, ZDHHC15, BCL11A, BCL11B, RTN4R, PARD3, PLXND1, CTNNA2, VAX1, CDKL5, AUTS2, EPHA7, FZD3, PPP3CA, BMPR1B, EPHB2, EPHB1, ATL1, MAP2, EFNB2, ADORA2A |
| GO:0032502 | developmental process | <0.0001 | <0.0001 | CDH18, PTPRF, LRGUK, FOXP2, MYO6, CNTN5, CSMD1, GRID2, FGF14, WIPF3, PLXDC1, CDH2, PRDM16, CAMK1D, MDGA2, WNT7A, NAV3, TENM1, ELMO1, DNM3, NOVA1, PANX3, ZMIZ1, SH2B2, MACROD2, USH2A, CLSTN2, FCER1G, DCN, TENM4, ULK4, PPARGC1A, RBFOX1, IL1RAPL1, SOX21, CD2AP, RAD51B, NTRK2, SHISA6, E2F7, CSMD2, DACH1, TCOF1, AFF2, ROBO2, LRTM2, MAP1S, CDH12, TDRD9, ZFP36L1, SDK1, TCF7L2, MBNL3, DAB1, MSI2, DMD, CPE, TTC39C, RBM47, MAGI2, ADAMTS2, A2M, ABI3BP, ARHGAP24, FLT3, KIFAP3, TP63, ADCYAP1R1, WT1, RAB3A, PREX2, PTPRG, GLUL, CLMP, PAX5, PKHD1, PPARGC1B, ASTN2, PTPRO, CRIM1, CNTN1, KIF1A, CTNND2, JAK3, NCAM1, SLC4A11, CBLB, MAP2K6, KATNAL1, CDH13, ARHGEF10. ANK2, ASPH, LAMA4, CADM1, DAAM2, GRIN2B, BCR, EMCN, TANC2, SLIT3, PKD1, GCNT4, IRX1, STOX2, ROR2, SEMA6A, CHST11, CHL1, TRPV1, NOTCH2, GABRB3, GABRB1, RELN, TCF7, COL1A1, DLG5, DBN1, ZNF521, GAL3ST1, SDCCAG8, HSPG2, EPHA6, DISC1, SYNGAP1, NRXN1, AKT3, TWIST2, PROX1, DNER, ZDHHC15, BCL11A, CPAMD8, ACAP3, RORC, ZFHX3, BCL11B, RTN4R, PARD3, CEP85L, CHD7, SPTBN1, PLXND1, FKBP6, FOXL1, MXRA8, FBN2, DOCK5, CTNNA2, VAX1, FGD4, FGF22, SKI, CCDC88A, TRPS1, MTA3, RAPGEFL1, SIM2, CDKL5, WWC2, PDGFD, LDB3, AUTS2, LCK, UTRN, TBX5, NEBL, CAMK1, EPHA7, APBA1, SGCA, KAZN, FZD3, CELSR1, PPP3CA, CACNA1G, BMPR1B, MTSS1, ADAM8, ERBB3, BSN, CPLX2, MNAT1, PBX1, COL4A2, SCARF1, BICD1, EPHB2, TUFT1, MYH10. EPHB1, ANGPT2, ADAM12, ATL1, ZMYND12, MRC2, GABRA2, NHEJ1, MAP2, ZBTB40. CHD2, SAMD14, TSPAN18, EFNB2, NKX2-2, ADORA2A, NBEAL2, KCNIP2, GAL, ZFPM2, SH3PXD2A, NSDHL, RAPGEF5, STAB2, MLLT3 |
| GO:0007409 | axonogenesis | <0.0001 | <0.0001 | CNTN5, WNT7A, CD2AP, NTRK2, ROBO2, LRTM2, MAP1S, DAB1, RAB3A, PTPRO, CNTN1, NCAM1, SLIT3, SEMA6A, CHL1, NOTCH2, RELN, EPHA6, DISC1, SYNGAP1, NRXN1, BCL11A, BCL11B, RTN4R, PARD3, PLXND1, CTNNA2, VAX1, CDKL5, AUTS2, EPHA7, FZD3, BMPR1B, EPHB2, EPHB1, ATL1, MAP2, EFNB2 |
| GO:0048858 | cell projection morphogenesis | <0.0001 | <0.0001 | CNTN5, WNT7A, DNM3, IL1RAPL1, CD2AP, NTRK2, ROBO2, LRTM2, MAP1S, DAB1, RAB3A, PREX2, PTPRO, CNTN1, KIF1A, CTNND2, NCAM1, TANC2, SLIT3, SEMA6A, CHL1, NOTCH2, RELN, DBN1, EPHA6, DISC1, SYNGAP1, NRXN1, ZDHHC15, BCL11A, BCL11B, RTN4R, PARD3, PLXND1, CTNNA2, VAX1, CDKL5, AUTS2, EPHA7, FZD3, PPP3CA, BMPR1B, EPHB2, EPHB1, ATL1, MAP2, EFNB2, ADORA2A |
| GO:0030182 | neuron differentiation | <0.0001 | <0.0001 | PTPRF, MYO6, CNTN5, GRID2, CAMK1D, MDGA2, WNT7A, TENM1, DNM3, ZMIZ1, USH2A, TENM4, ULK4, IL1RAPL1, SOX21, CD2AP, NTRK2, ROBO2, LRTM2, MAP1S, SDK1, DAB1, DMD, MAGI2, RAB3A, PREX2, PTPRG, PTPRO, CNTN1, KIF1A, CTNND2, NCAM1, TANC2, SLIT3, IRX1, ROR2, SEMA6A, CHL1, NOTCH2, GABRB1, RELN, DLG5, DBN1, ZNF521, EPHA6, DISC1, SYNGAP1, NRXN1, PROX1, ZDHHC15, BCL11A, ACAP3, ZFHX3, BCL11B, RTN4R, PARD3, PLXND1, CTNNA2, VAX1, CCDC88A, CDKL5, AUTS2, CAMK1, EPHA7, FZD3, PPP3CA, BMPR1B, PBX1, SCARF1, EPHB2, EPHB1, ATL1, MAP2, SAMD14, EFNB2, NKX2-2, ADORA2A, KCNIP2 |
| GO:0099175 | regulation of postsynapse organization | <0.0001 | <0.0001 | GRID1, GRID2, CDH2, WNT7A, ELMO1, DNM3, IL1RAPL1, KIF1A, GRIN2B, TANC2, ROR2, RELN, DBN1, DISC1, NRXN1, ZDHHC15, RTN4R, CDKL5, EPHA7, EPHB2 |
| GO:0032501 | multicellular organismal process | <0.0001 | <0.0001 | PTPRF, FOXP2, MYO6, CNTN5, CSMD1, GRID1, GRID2, FGF14, PLXDC1, CDH2, PRDM16, CAMK1D, MDGA2, WNT7A, NAV3, TENM1, ELMO1, DNM3, NOVA1, PANX3, ZMIZ1, P3H4, PLCL1, SH2B2, MACROD2, USH2A, CHRM5, CLSTN2, FCER1G, DCN, TENM4, ULK4, PPARGC1A, RBFOX1, IL1RAPL1, CUBN, SOX21, CD2AP, RAD51B, NTRK2, NBEA, E2F7, CSMD2, ESRRG, DACH1, TCOF1, NLRX1, SLC4A3, AFF2, ROBO2, SLCO3A1, LRTM2, MAP1S, ZFP36L1, SDK1, TCF7L2, DAB1, MSI2, DMD, CPE, TTC39C, RBM47, MAGI2, ADAMTS2, A2M, ABI3BP, ARHGAP24, FLT3, KIFAP3, TP63, ADCYAP1R1, WT1, RAB3A, PREX2, PTPRG, GLUL, CLMP, PAX5, PKHD1, PPARGC1B, ASTN2, PTPRO, CRIM1, PRRT1, CNTN1, KIF1A, DGKI, CTNND2, JAK3, NCAM1, CBLB, MAP2K6, MUC6, CDH13, ARHGEF10. ANK2, MARVELD3, ASPH, LAMA4, CADM1, DAAM2, GRIN2B, BCR, EMCN, TANC2, SLIT3, PKD1, MZB1, GCNT4, IRX1, STOX2, ROR2, SEMA6A, CHST11, CHL1, TRPV1, NOTCH2, GABRB3, GABRB1, RELN, TCF7, CACNB2, COL1A1, DLG5, DBN1, FCHO1, ZNF521, GAL3ST1, SDCCAG8, HSPG2, EPHA6, DISC1, SYNGAP1, NRXN1, AKT3, TWIST2, PROX1, DNER, ZDHHC15, BCL11A, CPAMD8, ACAP3, RORC, ZFHX3, BCL11B, RTN4R, PARD3, CEP85L, CHD7, SPTBN1, PLXND1, FOXL1, MXRA8, FBN2, DOCK5, CTNNA2, VAX1, RRH, SKI, CCDC88A, TRPS1, MGLL, RAPGEFL1, SIM2, PLCB2, CDKL5, WWC2, PDGFD, CTNNA3, LDB3, AUTS2, GLRA1, LCK, UTRN, TBX5, NEBL, CAMK1, EPHA7, APBA1, SGCA, KAZN, FZD3, CELSR1, TNNC2, PPP3CA, CACNA1G, BMPR1B, MTSS1, ADAM8, ERBB3, ADCY5, BSN, CPLX2, MNAT1, PBX1, COL4A2, SCARF1, EPHB2, REM1, TUFT1, EPHB1, ANGPT2, ADAM12, ATL1, MRC2, GABRA2, NHEJ1, MAP2, ZBTB40. SAMD14, TSPAN18, EFNB2, NKX2-2, ADORA2A, KCNIP2, GAL, ZFPM2, SH3PXD2A, NSDHL, RAPGEF5, STAB2, MLLT3 |
| GO:0048856 | anatomical structure development | <0.0001 | <0.0001 | CDH18, PTPRF, FOXP2, MYO6, CNTN5, CSMD1, GRID2, FGF14, PLXDC1, CDH2, CAMK1D, MDGA2, WNT7A, NAV3, TENM1, ELMO1, DNM3, NOVA1, ZMIZ1, SH2B2, MACROD2, USH2A, CLSTN2, FCER1G, DCN, TENM4, ULK4, PPARGC1A, RBFOX1, IL1RAPL1, SOX21, CD2AP, RAD51B, NTRK2, E2F7, CSMD2, DACH1, TCOF1, AFF2, ROBO2, LRTM2, MAP1S, CDH12, ZFP36L1, SDK1, TCF7L2, MBNL3, DAB1, MSI2, DMD, CPE, TTC39C, RBM47, MAGI2, ADAMTS2, A2M, ABI3BP, ARHGAP24, FLT3, KIFAP3, TP63, ADCYAP1R1, WT1, RAB3A, PREX2, PTPRG, GLUL, CLMP, PAX5, PKHD1, PPARGC1B, ASTN2, PTPRO, CRIM1, CNTN1, KIF1A, CTNND2, JAK3, NCAM1, CBLB, MAP2K6, CDH13, ARHGEF10. ANK2, ASPH, LAMA4, CADM1, DAAM2, GRIN2B, BCR, EMCN, TANC2, SLIT3, PKD1, GCNT4, IRX1, STOX2, ROR2, SEMA6A, CHST11, CHL1, TRPV1, NOTCH2, GABRB3, GABRB1, RELN, TCF7, COL1A1, DLG5, DBN1, ZNF521, GAL3ST1, SDCCAG8, HSPG2, EPHA6, DISC1, SYNGAP1, NRXN1, AKT3, PROX1, DNER, ZDHHC15, BCL11A, CPAMD8, ACAP3, RORC, ZFHX3, BCL11B, RTN4R, PARD3, CEP85L, CHD7, SPTBN1, PLXND1, FOXL1, MXRA8, FBN2, DOCK5, CTNNA2, VAX1, FGD4, FGF22, SKI, CCDC88A, TRPS1, RAPGEFL1, SIM2, CDKL5, PDGFD, LDB3, AUTS2, LCK, UTRN, TBX5, NEBL, CAMK1, EPHA7, APBA1, SGCA, KAZN, FZD3, CELSR1, PPP3CA, CACNA1G, BMPR1B, MTSS1, ADAM8, ERBB3, BSN, CPLX2, MNAT1, PBX1, COL4A2, SCARF1, BICD1, EPHB2, TUFT1, MYH10. EPHB1, ANGPT2, ADAM12, ATL1, ZMYND12, GABRA2, NHEJ1, MAP2, ZBTB40. CHD2, SAMD14, TSPAN18, EFNB2, NKX2-2, ADORA2A, NBEAL2, KCNIP2, GAL, ZFPM2, SH3PXD2A, NSDHL, RAPGEF5, STAB2, MLLT3 |
| GO:0048869 | cellular developmental process | <0.0001 | <0.0001 | PTPRF, LRGUK, MYO6, CNTN5, GRID2, WIPF3, CDH2, PRDM16, CAMK1D, MDGA2, WNT7A, NAV3, TENM1, DNM3, PANX3, ZMIZ1, SH2B2, USH2A, FCER1G, TENM4, ULK4, PPARGC1A, IL1RAPL1, SOX21, CD2AP, NTRK2, E2F7, TCOF1, ROBO2, LRTM2, MAP1S, TDRD9, ZFP36L1, SDK1, TCF7L2, MBNL3, DAB1, MSI2, DMD, RBM47, MAGI2, A2M, ABI3BP, ARHGAP24, FLT3, KIFAP3, TP63, ADCYAP1R1, WT1, RAB3A, PREX2, PTPRG, PAX5, PKHD1, PPARGC1B, ASTN2, PTPRO, CRIM1, CNTN1, KIF1A, CTNND2, JAK3, NCAM1, SLC4A11, MAP2K6, ARHGEF10. ANK2, LAMA4, CADM1, DAAM2, BCR, TANC2, SLIT3, IRX1, ROR2, SEMA6A, CHST11, CHL1, TRPV1, NOTCH2, GABRB1, RELN, TCF7, COL1A1, DLG5, DBN1, ZNF521, SDCCAG8, HSPG2, EPHA6, DISC1, SYNGAP1, NRXN1, TWIST2, PROX1, DNER, ZDHHC15, BCL11A, ACAP3, RORC, ZFHX3, BCL11B, RTN4R, PARD3, CEP85L, CHD7, PLXND1, FKBP6, FOXL1, MXRA8, FBN2, DOCK5, CTNNA2, VAX1, FGF22, SKI, CCDC88A, TRPS1, MTA3, SIM2, CDKL5, LDB3, AUTS2, LCK, TBX5, NEBL, CAMK1, EPHA7, KAZN, FZD3, CELSR1, PPP3CA, BMPR1B, MTSS1, ADAM8, ERBB3, CPLX2, PBX1, COL4A2, SCARF1, EPHB2, EPHB1, ANGPT2, ADAM12, ATL1, ZMYND12, MRC2, NHEJ1, MAP2, CHD2, SAMD14, EFNB2, NKX2-2, ADORA2A, NBEAL2, KCNIP2, ZFPM2, SH3PXD2A, MLLT3 |
| GO:0030154 | cell differentiation | <0.0001 | <0.0001 | PTPRF, LRGUK, MYO6, CNTN5, GRID2, WIPF3, CDH2, PRDM16, CAMK1D, MDGA2, WNT7A, NAV3, TENM1, DNM3, PANX3, ZMIZ1, SH2B2, USH2A, FCER1G, TENM4, ULK4, PPARGC1A, IL1RAPL1, SOX21, CD2AP, NTRK2, E2F7, TCOF1, ROBO2, LRTM2, MAP1S, TDRD9, ZFP36L1, SDK1, TCF7L2, MBNL3, DAB1, MSI2, DMD, RBM47, MAGI2, A2M, ABI3BP, ARHGAP24, FLT3, KIFAP3, TP63, ADCYAP1R1, WT1, RAB3A, PREX2, PTPRG, PAX5, PKHD1, PPARGC1B, ASTN2, PTPRO, CRIM1, CNTN1, KIF1A, CTNND2, JAK3, NCAM1, SLC4A11, MAP2K6, ARHGEF10. ANK2, LAMA4, CADM1, DAAM2, BCR, TANC2, SLIT3, IRX1, ROR2, SEMA6A, CHST11, CHL1, TRPV1, NOTCH2, GABRB1, RELN, TCF7, COL1A1, DLG5, DBN1, ZNF521, SDCCAG8, HSPG2, EPHA6, DISC1, SYNGAP1, NRXN1, TWIST2, PROX1, DNER, ZDHHC15, BCL11A, ACAP3, RORC, ZFHX3, BCL11B, RTN4R, PARD3, CEP85L, CHD7, PLXND1, FKBP6, FOXL1, MXRA8, FBN2, DOCK5, CTNNA2, VAX1, FGF22, SKI, CCDC88A, TRPS1, MTA3, SIM2, CDKL5, LDB3, AUTS2, LCK, TBX5, NEBL, CAMK1, EPHA7, KAZN, FZD3, CELSR1, PPP3CA, BMPR1B, MTSS1, ADAM8, ERBB3, CPLX2, PBX1, COL4A2, SCARF1, EPHB2, EPHB1, ANGPT2, ADAM12, ATL1, ZMYND12, MRC2, NHEJ1, MAP2, CHD2, SAMD14, EFNB2, NKX2-2, ADORA2A, NBEAL2, KCNIP2, ZFPM2, SH3PXD2A, MLLT3 |
| GO:0061564 | axon development | <0.0001 | <0.0001 | PTPRF, CNTN5, WNT7A, CD2AP, NTRK2, ROBO2, LRTM2, MAP1S, DAB1, RAB3A, PTPRO, CNTN1, NCAM1, SLIT3, SEMA6A, CHL1, NOTCH2, RELN, EPHA6, DISC1, SYNGAP1, NRXN1, BCL11A, BCL11B, RTN4R, PARD3, PLXND1, CTNNA2, VAX1, CDKL5, AUTS2, EPHA7, FZD3, BMPR1B, SCARF1, EPHB2, EPHB1, ATL1, MAP2, EFNB2 |
| GO:0099536 | synaptic signaling | <0.0001 | <0.0001 | GABBR2, GRID1, DLGAP1, GRID2, WNT7A, PLCL1, PTPRN2, CHRM5, CLSTN2, IL1RAPL1, CD2AP, NTRK2, SHISA6, CACNA1B, DLGAP2, RPS6KA2, DMD, RAB3A, CLMP, PRRT1, DGKI, RASGRF2, GRIN2B, BCR, ROR2, TRPV1, GABRB3, GABRB1, RELN, CACNB2, DBN1, DISC1, SYNGAP1, NRXN1, CDKL5, HTR1F, GLRA1, UTRN, EPHA7, APBA1, PPP3CA, CACNA1G, BSN, CPLX2, EPHB2, EPHB1, GABRA2, ADORA2A, KCNIP2 |
| GO:0060996 | dendritic spine development | <0.0001 | <0.0001 | WNT7A, DNM3, IL1RAPL1, SDK1, KIF1A, CTNND2, TANC2, RELN, DLG5, DBN1, DISC1, ZDHHC15, CDKL5, CAMK1, EPHB2, EPHB1 |
| GO:0099537 | trans-synaptic signaling | <0.0001 | <0.0001 | GABBR2, GRID1, DLGAP1, GRID2, WNT7A, PLCL1, PTPRN2, CHRM5, CLSTN2, IL1RAPL1, CD2AP, NTRK2, SHISA6, CACNA1B, DLGAP2, RPS6KA2, DMD, RAB3A, CLMP, PRRT1, DGKI, RASGRF2, GRIN2B, BCR, ROR2, TRPV1, GABRB3, GABRB1, RELN, CACNB2, DBN1, DISC1, SYNGAP1, NRXN1, CDKL5, HTR1F, GLRA1, EPHA7, APBA1, PPP3CA, CACNA1G, BSN, CPLX2, EPHB2, EPHB1, GABRA2, ADORA2A, KCNIP2 |
| GO:0120036 | plasma membrane bounded cell projection organization | <0.0001 | <0.0001 | PTPRF, LRGUK, CNTN5, GRID2, CAMK1D, WNT7A, TENM1, DNM3, ULK4, IL1RAPL1, CD2AP, NTRK2, ROBO2, LRTM2, MAP1S, SDK1, DAB1, DMD, TTC39C, MAGI2, ARHGAP24, KIFAP3, RAB3A, PREX2, PTPRG, PKHD1, PTPRO, CNTN1, KIF1A, IFT43, CTNND2, NCAM1, CDH13, DAAM2, GRIN2B, TANC2, SLIT3, ROR2, SEMA6A, CCDC13, CHL1, NOTCH2, RELN, DLG5, DBN1, SDCCAG8, EPHA6, DISC1, SYNGAP1, NRXN1, ZDHHC15, BCL11A, ACAP3, BCL11B, RTN4R, PARD3, PLXND1, CTNNA2, VAX1, FGD4, CCDC88A, RILP, CDKL5, AUTS2, CAMK1, EPHA7, FZD3, PPP3CA, BMPR1B, MTSS1, PCNT, SCARF1, EPHB2, EPHB1, ATL1, ZMYND12, LRRC23, MAP2, SAMD14, EFNB2, ADORA2A |
| GO:0007416 | synapse assembly | <0.0001 | <0.0001 | CNTN5, GRID2, CDH2, WNT7A, ELMO1, DNM3, CLSTN2, IL1RAPL1, NTRK2, CSMD2, ROBO2, LRTM2, SDK1, GABRB3, RELN, DLG5, NRXN1, DNER, RTN4R, PLXND1, EPHA7, BSN, EPHB2, EPHB1, GABRA2 |
| GO:0120035 | regulation of plasma membrane bounded cell projection organization | <0.0001 | <0.0001 | PTPRF, GRID2, CAMK1D, WNT7A, TENM1, DNM3, ULK4, IL1RAPL1, NTRK2, ROBO2, DAB1, DMD, MAGI2, ARHGAP24, PTPRG, PTPRO, CNTN1, KIF1A, DAAM2, GRIN2B, TANC2, ROR2, RELN, DBN1, SDCCAG8, DISC1, SYNGAP1, NRXN1, ZDHHC15, BCL11A, ACAP3, RTN4R, PLXND1, CTNNA2, CCDC88A, CDKL5, AUTS2, CAMK1, EPHA7, PPP3CA, SCARF1, EPHB2, MAP2, EFNB2 |
| GO:0030030 | cell projection organization | <0.0001 | <0.0001 | PTPRF, LRGUK, CNTN5, GRID2, CAMK1D, WNT7A, TENM1, DNM3, PARVG, ULK4, IL1RAPL1, CD2AP, NTRK2, ROBO2, LRTM2, MAP1S, SDK1, DAB1, DMD, TTC39C, MAGI2, ARHGAP24, KIFAP3, RAB3A, PREX2, PTPRG, PKHD1, PTPRO, CNTN1, KIF1A, IFT43, CTNND2, NCAM1, CDH13, DAAM2, GRIN2B, TANC2, SLIT3, ROR2, SEMA6A, CCDC13, CHL1, NOTCH2, RELN, DLG5, DBN1, SDCCAG8, EPHA6, DISC1, SYNGAP1, NRXN1, ZDHHC15, BCL11A, ACAP3, BCL11B, RTN4R, PARD3, PLXND1, CTNNA2, VAX1, FGD4, CCDC88A, RILP, CDKL5, AUTS2, CAMK1, EPHA7, FZD3, PPP3CA, BMPR1B, MTSS1, PCNT, SCARF1, EPHB2, EPHB1, ATL1, ZMYND12, LRRC23, MAP2, SAMD14, EFNB2, ADORA2A |
| GO:0007268 | chemical synaptic transmission | <0.0001 | <0.0001 | GABBR2, GRID1, DLGAP1, GRID2, WNT7A, PLCL1, PTPRN2, CHRM5, CLSTN2, CD2AP, NTRK2, SHISA6, CACNA1B, DLGAP2, RPS6KA2, DMD, RAB3A, CLMP, PRRT1, DGKI, RASGRF2, GRIN2B, BCR, ROR2, TRPV1, GABRB3, GABRB1, RELN, CACNB2, DBN1, DISC1, SYNGAP1, NRXN1, CDKL5, HTR1F, GLRA1, EPHA7, APBA1, PPP3CA, CACNA1G, BSN, CPLX2, EPHB2, EPHB1, GABRA2, ADORA2A, KCNIP2 |
| GO:0098916 | anterograde trans-synaptic signaling | <0.0001 | <0.0001 | GABBR2, GRID1, DLGAP1, GRID2, WNT7A, PLCL1, PTPRN2, CHRM5, CLSTN2, CD2AP, NTRK2, SHISA6, CACNA1B, DLGAP2, RPS6KA2, DMD, RAB3A, CLMP, PRRT1, DGKI, RASGRF2, GRIN2B, BCR, ROR2, TRPV1, GABRB3, GABRB1, RELN, CACNB2, DBN1, DISC1, SYNGAP1, NRXN1, CDKL5, HTR1F, GLRA1, EPHA7, APBA1, PPP3CA, CACNA1G, BSN, CPLX2, EPHB2, EPHB1, GABRA2, ADORA2A, KCNIP2 |
| GO:0009653 | anatomical structure morphogenesis | <0.0001 | <0.0001 | CDH18, MYO6, CNTN5, CSMD1, GRID2, PLXDC1, CDH2, WNT7A, DNM3, ZMIZ1, DCN, TENM4, IL1RAPL1, CD2AP, NTRK2, E2F7, TCOF1, ROBO2, LRTM2, MAP1S, CDH12, ZFP36L1, SDK1, DAB1, DMD, CPE, TTC39C, ARHGAP24, TP63, WT1, RAB3A, PREX2, GLUL, PAX5, PKHD1, PPARGC1B, ASTN2, PTPRO, CNTN1, KIF1A, CTNND2, NCAM1, CDH13, ANK2, ASPH, BCR, EMCN, TANC2, SLIT3, PKD1, GCNT4, IRX1, SEMA6A, CHST11, CHL1, NOTCH2, RELN, COL1A1, DLG5, DBN1, SDCCAG8, HSPG2, EPHA6, DISC1, SYNGAP1, NRXN1, AKT3, PROX1, ZDHHC15, BCL11A, BCL11B, RTN4R, PARD3, CHD7, SPTBN1, PLXND1, FOXL1, FBN2, DOCK5, CTNNA2, VAX1, FGD4, FGF22, SKI, CDKL5, LDB3, AUTS2, TBX5, NEBL, CAMK1, EPHA7, FZD3, CELSR1, PPP3CA, BMPR1B, MTSS1, ADAM8, PBX1, COL4A2, BICD1, EPHB2, TUFT1, MYH10. EPHB1, ANGPT2, ADAM12, ATL1, MAP2, TSPAN18, EFNB2, ADORA2A, NBEAL2, ZFPM2, SH3PXD2A, STAB2 |
| GO:0000902 | cell morphogenesis | <0.0001 | <0.0001 | CDH18, CNTN5, CDH2, WNT7A, DNM3, IL1RAPL1, CD2AP, NTRK2, ROBO2, LRTM2, MAP1S, CDH12, DAB1, RAB3A, PREX2, PKHD1, PTPRO, CNTN1, KIF1A, CTNND2, NCAM1, CDH13, TANC2, SLIT3, SEMA6A, CHL1, NOTCH2, RELN, DBN1, EPHA6, DISC1, SYNGAP1, NRXN1, PROX1, ZDHHC15, BCL11A, BCL11B, RTN4R, PARD3, PLXND1, CTNNA2, VAX1, FGD4, CDKL5, AUTS2, EPHA7, FZD3, PPP3CA, BMPR1B, ADAM8, EPHB2, MYH10. EPHB1, ATL1, MAP2, EFNB2, ADORA2A, NBEAL2 |
| GO:0031344 | regulation of cell projection organization | <0.0001 | <0.0001 | PTPRF, GRID2, CAMK1D, WNT7A, TENM1, DNM3, ULK4, IL1RAPL1, NTRK2, ROBO2, DAB1, DMD, MAGI2, ARHGAP24, PTPRG, PTPRO, CNTN1, KIF1A, DAAM2, GRIN2B, TANC2, ROR2, RELN, DBN1, SDCCAG8, DISC1, SYNGAP1, NRXN1, ZDHHC15, BCL11A, ACAP3, RTN4R, PLXND1, CTNNA2, CCDC88A, CDKL5, AUTS2, CAMK1, EPHA7, PPP3CA, SCARF1, EPHB2, MAP2, EFNB2 |
| GO:0034329 | cell junction assembly | <0.0001 | <0.0001 | CDH18, CNTN5, GRID2, CDH2, WNT7A, ELMO1, DNM3, CLSTN2, IL1RAPL1, NTRK2, CSMD2, ROBO2, LRTM2, CDH12, SDK1, PTPRO, CDH13, ANK2, MARVELD3, BCR, GABRB3, RELN, DLG5, NRXN1, DNER, RTN4R, PARD3, PLXND1, TBX5, EPHA7, BSN, DST, EPHB2, EPHB1, GABRA2, TNS1 |
| GO:0060998 | regulation of dendritic spine development | <0.0001 | <0.0001 | DNM3, IL1RAPL1, SDK1, KIF1A, TANC2, RELN, DLG5, DBN1, DISC1, CDKL5, CAMK1, EPHB2 |
| GO:0098742 | cell-cell adhesion via plasma-membrane adhesion molecules | <0.0001 | <0.0001 | CDH18, PTPRF, PCDH7, GRID2, CDH2, TENM1, CLSTN2, TENM4, IL1RAPL1, ROBO2, CDH12, SDK1, DAB1, KIFAP3, PTPRT, CDH13, CADM1, PKD1, NRXN1, CELSR1, SCARF1 |
| GO:0051960 | regulation of nervous system development | <0.0001 | <0.0001 | GRID2, WNT7A, CLSTN2, TENM4, IL1RAPL1, NTRK2, ROBO2, LRTM2, DAB1, KIFAP3, DAAM2, RELN, DLG5, DBN1, DISC1, SYNGAP1, NRXN1, PROX1, BCL11A, RTN4R, PARD3, CHD7, PLXND1, VAX1, SKI, CDKL5, EPHA7, FZD3, PPP3CA, EPHB2, EPHB1, MAP2, NKX2-2 |
| GO:0007267 | cell-cell signaling | <0.0001 | <0.0001 | GABBR2, GRID1, DLGAP1, GRID2, FGF14, WNT7A, PANX3, PLCL1, PTPRN2, CHRM5, CLSTN2, IL1RAPL1, CD2AP, NTRK2, SHISA6, CACNA1B, DLGAP2, RPS6KA2, TCF7L2, DMD, CPE, TP63, RAB3A, CLMP, PRRT1, DGKI, RASGRF2, MAP2K6, ANK2, GRIN2B, BCR, ROR2, PFKL, TRPV1, GABRB3, GABRB1, RELN, CACNB2, DBN1, DISC1, SYNGAP1, NRXN1, CHD7, FOXL1, CDKL5, HTR1F, GLRA1, UTRN, TBX5, EPHA7, APBA1, PPP3CA, CACNA1G, ADAM8, ADCY5, BSN, CPLX2, EPHB2, EPHB1, GABRA2, EFNB2, ADORA2A, KCNIP2, GAL |
| GO:0060997 | dendritic spine morphogenesis | <0.0001 | <0.0001 | WNT7A, DNM3, IL1RAPL1, KIF1A, CTNND2, TANC2, RELN, DBN1, ZDHHC15, EPHB2, EPHB1 |
| GO:0007417 | central nervous system development | <0.0001 | <0.0001 | FOXP2, CNTN5, GRID2, PLXDC1, CDH2, MDGA2, WNT7A, ZMIZ1, MACROD2, TENM4, SOX21, NTRK2, AFF2, ROBO2, MAP1S, DAB1, MSI2, DMD, PAX5, CNTN1, DAAM2, GRIN2B, BCR, PKD1, ROR2, TRPV1, GABRB1, RELN, DLG5, HSPG2, DISC1, NRXN1, AKT3, PROX1, DNER, BCL11B, RTN4R, CHD7, SPTBN1, MXRA8, CTNNA2, VAX1, SKI, EPHA7, FZD3, CELSR1, BMPR1B, MNAT1, PBX1, EPHB2, EPHB1, NHEJ1, MAP2, NKX2-2, ADORA2A |
| GO:0001764 | neuron migration | <0.0001 | <0.0001 | ZMIZ1, ULK4, NTRK2, DAB1, ASTN2, SEMA6A, CHL1, RELN, SDCCAG8, DISC1, DNER, ACAP3, CEP85L, CTNNA2, VAX1, CDKL5, AUTS2, FZD3, CELSR1 |
| GO:0048468 | cell development | <0.0001 | <0.0001 | PTPRF, CNTN5, GRID2, CDH2, CAMK1D, WNT7A, TENM1, DNM3, ZMIZ1, FCER1G, TENM4, ULK4, IL1RAPL1, CD2AP, NTRK2, TCOF1, ROBO2, LRTM2, MAP1S, ZFP36L1, SDK1, DAB1, MSI2, DMD, RBM47, MAGI2, FLT3, KIFAP3, TP63, WT1, RAB3A, PREX2, PTPRG, PAX5, PKHD1, PPARGC1B, PTPRO, CNTN1, KIF1A, CTNND2, JAK3, NCAM1, ARHGEF10. ANK2, DAAM2, BCR, TANC2, SLIT3, IRX1, ROR2, SEMA6A, CHST11, CHL1, TRPV1, NOTCH2, GABRB1, RELN, TCF7, DLG5, DBN1, EPHA6, DISC1, SYNGAP1, NRXN1, PROX1, DNER, ZDHHC15, BCL11A, ACAP3, RORC, BCL11B, RTN4R, PARD3, CHD7, PLXND1, MXRA8, CTNNA2, VAX1, SKI, CCDC88A, CDKL5, LDB3, AUTS2, LCK, TBX5, NEBL, CAMK1, EPHA7, FZD3, PPP3CA, BMPR1B, ADAM8, ERBB3, PBX1, SCARF1, EPHB2, EPHB1, ANGPT2, ATL1, ZMYND12, NHEJ1, MAP2, CHD2, SAMD14, EFNB2, NKX2-2, ADORA2A, NBEAL2, KCNIP2, SH3PXD2A, MLLT3 |
| GO:0016477 | cell migration | <0.0001 | <0.0001 | CDH18, PTPRF, CDH2, CAMK1D, WNT7A, NAV3, ZMIZ1, FCER1G, DCN, ULK4, CD2AP, NTRK2, DACH1, CDH12, DAB1, MAGI2, ARHGAP24, PTPRG, GLUL, ASTN2, PTPRO, PTPRT, CDH13, MARVELD3, LAMA4, DAAM2, BCR, PODXL2, ROR2, SEMA6A, CHL1, RELN, COL1A1, DLG5, SDCCAG8, DISC1, AKT3, TWIST2, PROX1, DNER, ACAP3, BCL11B, CEP85L, PLXND1, CCR10. DOCK5, CTNNA2, VAX1, FGF22, CCDC88A, CDKL5, WWC2, PDGFD, CTNNA3, AUTS2, LCK, TBX5, FZD3, CELSR1, PPP3CA, ADAM8, EPHB2, EPHB1, ANGPT2, LRCH1, EFNB2, PIK3C2B, TNS1 |
| GO:0007154 | cell communication | <0.0001 | <0.0001 | PTPRF, MYO6, GABBR2, GRID1, ARHGEF10L, DLGAP1, GRID2, FGF14, CDH2, PRDM16, WNT7A, TENM1, ELMO1, PANX3, ZMIZ1, PLCL1, SH2B2, PTPRN2, RCAN2, CHRM5, CLSTN2, FCER1G, RBMS3, DCN, TENM4, ULK4, IL1RAPL1, INCENP, CD2AP, NTRK2, SHISA6, ESRRG, NLRX1, CACNA1B, ROBO2, DLGAP2, ERG, ZFP36L1, RPS6KA2, TCF7L2, PIK3R5, DAB1, DMD, CPE, RBM47, MAGI2, PDE10A, ARHGAP24, FLT3, KIFAP3, TP63, ADCYAP1R1, RASSF8, RAB3A, PREX2, PTPRG, CLMP, PKHD1, PPARGC1B, PLCD1, PTPRO, CRIM1, PRRT1, CNTN1, LTBP1, DGKI, CTNND2, PTPRT, ARHGAP19, RASGRF2, CTNNAL1, JAK3, NCAM1, CBLB, MAP2K6, CNKSR2, PEX5L, CDH13, ARHGEF10. ANK2, MARVELD3, ASPH, CADM1, DAAM2, GRIN2B, BCR, SLIT3, PKD1, MZB1, ROR2, SEMA6A, PFKL, CHST11, CHL1, TRPV1, NOTCH2, GABRB3, GABRB1, RELN, TCF7, CACNB2, COL1A1, DLG5, DBN1, FCHO1, HSPG2, NRG4, EPHA6, DISC1, SYNGAP1, NRXN1, AKT3, MAP3K12, PROX1, DNER, RGL1, PIK3C2G, RORC, RTN4R, PARD3, CHD7, SPTBN1, PLXND1, CDT1, FOXL1, CCR10. FBN2, DOCK5, FMOD, FGD4, RRH, FGF22, SKI, CCDC88A, FNIP2, MGLL, RAPGEFL1, PLCB2, CDKL5, WWC2, PDGFD, CTNNA3, HTR1F, VIPR1, FRMPD1, LRRN2, AUTS2, GLRA1, LCK, UTRN, TBX5, CAMK1, EPHA7, APBA1, FZD3, CELSR1, PPP3CA, CACNA1G, BMPR1B, MTSS1, ADAM8, ERBB3, ADCY5, SH2D3C, BSN, CPLX2, PCNT, COL4A2, BICD1, DST, EPHB2, REM1, TUFT1, EPHB1, ANGPT2, PRKAG2, ADAM12, GABRA2, ZNF385B, SAMD14, EFNB2, NKX2-2, ADORA2A, KCNIP2, GAL, PIK3C2B, CAMKMT, NSDHL, RAPGEF5, MLLT3 |
| GO:0097061 | dendritic spine organization | <0.0001 | <0.0001 | WNT7A, DNM3, IL1RAPL1, KIF1A, CTNND2, GRIN2B, TANC2, RELN, DBN1, ZDHHC15, EPHB2, EPHB1 |
| GO:0023052 | signaling | <0.0001 | <0.0001 | PTPRF, MYO6, GABBR2, GRID1, ARHGEF10L, DLGAP1, GRID2, FGF14, CDH2, PRDM16, WNT7A, TENM1, ELMO1, PANX3, ZMIZ1, PLCL1, SH2B2, PTPRN2, RCAN2, CHRM5, CLSTN2, FCER1G, RBMS3, DCN, TENM4, ULK4, IL1RAPL1, INCENP, CD2AP, NTRK2, SHISA6, ESRRG, NLRX1, CACNA1B, ROBO2, DLGAP2, ERG, ZFP36L1, RPS6KA2, TCF7L2, PIK3R5, DAB1, DMD, CPE, RBM47, MAGI2, PDE10A, ARHGAP24, FLT3, KIFAP3, TP63, ADCYAP1R1, RASSF8, RAB3A, PREX2, PTPRG, CLMP, PKHD1, PPARGC1B, PLCD1, PTPRO, CRIM1, PRRT1, CNTN1, LTBP1, DGKI, CTNND2, PTPRT, ARHGAP19, RASGRF2, CTNNAL1, JAK3, NCAM1, CBLB, MAP2K6, CNKSR2, PEX5L, CDH13, ARHGEF10. ANK2, MARVELD3, ASPH, CADM1, DAAM2, GRIN2B, BCR, SLIT3, PKD1, MZB1, ROR2, SEMA6A, PFKL, CHST11, CHL1, TRPV1, NOTCH2, GABRB3, GABRB1, RELN, TCF7, CACNB2, COL1A1, DLG5, DBN1, FCHO1, HSPG2, NRG4, EPHA6, DISC1, SYNGAP1, NRXN1, AKT3, MAP3K12, PROX1, DNER, RGL1, PIK3C2G, RORC, RTN4R, PARD3, CHD7, SPTBN1, PLXND1, CDT1, FOXL1, CCR10. FBN2, DOCK5, FMOD, FGD4, RRH, FGF22, SKI, CCDC88A, FNIP2, MGLL, RAPGEFL1, PLCB2, CDKL5, WWC2, PDGFD, HTR1F, VIPR1, FRMPD1, LRRN2, AUTS2, GLRA1, LCK, UTRN, TBX5, CAMK1, EPHA7, APBA1, FZD3, CELSR1, PPP3CA, CACNA1G, BMPR1B, MTSS1, ADAM8, ERBB3, ADCY5, SH2D3C, BSN, CPLX2, PCNT, COL4A2, BICD1, DST, EPHB2, REM1, TUFT1, EPHB1, ANGPT2, PRKAG2, ADAM12, GABRA2, ZNF385B, SAMD14, EFNB2, NKX2-2, ADORA2A, KCNIP2, GAL, PIK3C2B, CAMKMT, NSDHL, RAPGEF5, MLLT3 |
| GO:0048813 | dendrite morphogenesis | <0.0001 | <0.0001 | WNT7A, DNM3, IL1RAPL1, PREX2, KIF1A, CTNND2, TANC2, RELN, DBN1, ZDHHC15, CTNNA2, CDKL5, PPP3CA, EPHB2, EPHB1, MAP2 |
| GO:0048870 | cell motility | <0.0001 | <0.0001 | CDH18, PTPRF, CDH2, CAMK1D, WNT7A, NAV3, ELMO1, ZMIZ1, FCER1G, DCN, ULK4, CD2AP, NTRK2, DACH1, CDH12, DAB1, MAGI2, ARHGAP24, PTPRG, GLUL, ASTN2, PTPRO, PTPRT, CDH13, MARVELD3, LAMA4, DAAM2, BCR, PODXL2, ROR2, SEMA6A, CHL1, RELN, COL1A1, DLG5, SDCCAG8, DISC1, AKT3, TWIST2, PROX1, DNER, ACAP3, BCL11B, CEP85L, PLXND1, CCR10. DOCK5, CTNNA2, VAX1, FGF22, SKI, CCDC88A, CDKL5, WWC2, PDGFD, CTNNA3, AUTS2, LCK, TBX5, FZD3, CELSR1, PPP3CA, ADAM8, DST, EPHB2, EPHB1, ANGPT2, ZMYND12, LRCH1, LRRC23, EFNB2, PIK3C2B, TNS1 |
| GO:0051961 | negative regulation of nervous system development | 0.001 | 0.001 | WNT7A, ROBO2, DAB1, KIFAP3, DAAM2, SYNGAP1, PROX1, BCL11A, RTN4R, VAX1, SKI, EPHA7, PPP3CA, EPHB2, MAP2 |
| GO:0070243 | regulation of thymocyte apoptotic process | 0.001 | 0.001 | KIFAP3, JAK3, RORC, BCL11B, ADAM8 |
| GO:0007411 | axon guidance | 0.001 | 0.001 | CNTN5, ROBO2, LRTM2, PTPRO, CNTN1, NCAM1, SLIT3, SEMA6A, CHL1, NOTCH2, RELN, EPHA6, NRXN1, BCL11B, VAX1, EPHA7, FZD3, BMPR1B, EPHB2, EPHB1, EFNB2 |
| GO:0097485 | neuron projection guidance | 0.001 | 0.001 | CNTN5, ROBO2, LRTM2, PTPRO, CNTN1, NCAM1, SLIT3, SEMA6A, CHL1, NOTCH2, RELN, EPHA6, NRXN1, BCL11B, VAX1, EPHA7, FZD3, BMPR1B, EPHB2, EPHB1, EFNB2 |
| GO:0051965 | positive regulation of synapse assembly | 0.001 | 0.001 | GRID2, WNT7A, CLSTN2, IL1RAPL1, NTRK2, LRTM2, DLG5, NRXN1, EPHB2, EPHB1 |
| GO:0098609 | cell-cell adhesion | 0.001 | 0.001 | CDH18, PTPRF, CNTN5, PCDH7, GRID2, CDH2, TENM1, ZMIZ1, CLSTN2, TENM4, IL1RAPL1, CD2AP, ROBO2, CDH12, ZFP36L1, SDK1, DAB1, KIFAP3, PKHD1, ASTN2, CNTN1, CTNND2, PTPRT, JAK3, CBLB, CDH13, CADM1, EMCN, PKD1, PODXL2, DLG5, FCHO1, NRXN1, CTNNA2, CTNNA3, LCK, EPHA7, CELSR1, PPP3CA, ADAM8, SCARF1, EFNB2, ADORA2A |
| GO:0051239 | regulation of multicellular organismal process | 0.001 | 0.001 | GRID2, PRDM16, WNT7A, NAV3, NOVA1, PANX3, ZMIZ1, PLCL1, CLSTN2, FCER1G, DCN, TENM4, PPARGC1A, IL1RAPL1, CD2AP, NTRK2, ESRRG, NLRX1, SLC4A3, ROBO2, LRTM2, ZFP36L1, TCF7L2, DAB1, DMD, RBM47, ABI3BP, KIFAP3, TP63, WT1, PTPRG, GLUL, PPARGC1B, PTPRO, JAK3, CBLB, MAP2K6, ANK2, MARVELD3, ASPH, LAMA4, CADM1, DAAM2, GRIN2B, BCR, MZB1, ROR2, SEMA6A, TRPV1, NOTCH2, RELN, TCF7, CACNB2, COL1A1, DLG5, DBN1, FCHO1, HSPG2, DISC1, SYNGAP1, NRXN1, AKT3, PROX1, BCL11A, ZFHX3, RTN4R, PARD3, CHD7, SPTBN1, PLXND1, FBN2, DOCK5, VAX1, SKI, TRPS1, MGLL, CDKL5, WWC2, CTNNA3, GLRA1, LCK, TBX5, EPHA7, SGCA, FZD3, TNNC2, PPP3CA, CACNA1G, BMPR1B, ADAM8, ERBB3, PBX1, COL4A2, EPHB2, REM1, EPHB1, ANGPT2, ADAM12, MAP2, TSPAN18, EFNB2, NKX2-2, ADORA2A, KCNIP2, GAL, ZFPM2 |
| GO:0099177 | regulation of trans-synaptic signaling | 0.001 | 0.001 | GRID1, GRID2, WNT7A, PLCL1, CLSTN2, CD2AP, NTRK2, SHISA6, CACNA1B, RAB3A, CLMP, PRRT1, DGKI, RASGRF2, GRIN2B, BCR, ROR2, RELN, DBN1, DISC1, SYNGAP1, NRXN1, CDKL5, EPHA7, APBA1, PPP3CA, BSN, CPLX2, EPHB2, EPHB1, ADORA2A |
| GO:0050804 | modulation of chemical synaptic transmission | 0.001 | 0.001 | GRID1, GRID2, WNT7A, PLCL1, CLSTN2, CD2AP, NTRK2, SHISA6, CACNA1B, RAB3A, CLMP, PRRT1, DGKI, RASGRF2, GRIN2B, BCR, ROR2, RELN, DBN1, DISC1, SYNGAP1, NRXN1, CDKL5, EPHA7, APBA1, PPP3CA, BSN, CPLX2, EPHB2, EPHB1, ADORA2A |
| GO:0106027 | neuron projection organization | 0.001 | 0.001 | WNT7A, DNM3, IL1RAPL1, KIF1A, CTNND2, GRIN2B, TANC2, RELN, DBN1, ZDHHC15, EPHB2, EPHB1 |
| GO:0050768 | negative regulation of neurogenesis | 0.001 | 0.001 | WNT7A, DAB1, KIFAP3, DAAM2, SYNGAP1, PROX1, BCL11A, RTN4R, VAX1, SKI, EPHA7, PPP3CA, EPHB2, MAP2 |
| GO:0050767 | regulation of neurogenesis | 0.001 | 0.001 | WNT7A, TENM4, IL1RAPL1, NTRK2, ROBO2, DAB1, KIFAP3, DAAM2, RELN, DBN1, DISC1, SYNGAP1, PROX1, BCL11A, RTN4R, CHD7, PLXND1, VAX1, SKI, CDKL5, EPHA7, FZD3, PPP3CA, EPHB2, MAP2, NKX2-2 |
| GO:0010976 | positive regulation of neuron projection development | 0.002 | 0.002 | CAMK1D, IL1RAPL1, NTRK2, DMD, MAGI2, CNTN1, ROR2, RELN, DBN1, DISC1, ZDHHC15, BCL11A, CAMK1, SCARF1, EPHB2 |
| GO:0001568 | blood vessel development | 0.002 | 0.002 | PLXDC1, CDH2, WNT7A, ZMIZ1, DCN, NTRK2, E2F7, ROBO2, ZFP36L1, TCF7L2, ARHGAP24, WT1, GLUL, CDH13, LAMA4, EMCN, PKD1, SEMA6A, NOTCH2, COL1A1, HSPG2, NRXN1, AKT3, PROX1, CHD7, PLXND1, PDGFD, TBX5, ADAM8, COL4A2, EPHB2, EPHB1, ANGPT2, ADAM12, TSPAN18, EFNB2, ZFPM2, NSDHL, STAB2 |
| GO:0007156 | homophilic cell adhesion via plasma membrane adhesion molecules | 0.002 | 0.002 | CDH18, PCDH7, CDH2, CLSTN2, ROBO2, CDH12, SDK1, PTPRT, CDH13, CADM1, PKD1, CELSR1 |
| GO:0035239 | tube morphogenesis | 0.002 | 0.002 | CSMD1, PLXDC1, CDH2, WNT7A, ZMIZ1, DCN, NTRK2, E2F7, ZFP36L1, ARHGAP24, TP63, WT1, GLUL, PKHD1, CDH13, EMCN, PKD1, IRX1, SEMA6A, NOTCH2, DLG5, SDCCAG8, HSPG2, NRXN1, AKT3, PROX1, CHD7, PLXND1, SKI, TBX5, EPHA7, FZD3, CELSR1, MTSS1, ADAM8, PBX1, COL4A2, EPHB2, EPHB1, ANGPT2, ADAM12, TSPAN18, EFNB2, ZFPM2, STAB2 |
| GO:0061001 | regulation of dendritic spine morphogenesis | 0.003 | 0.003 | DNM3, IL1RAPL1, KIF1A, TANC2, RELN, DBN1, ZDHHC15, EPHB2 |
| GO:0031290 | retinal ganglion cell axon guidance | 0.003 | 0.003 | ROBO2, PTPRO, EPHA7, BMPR1B, EPHB2, EPHB1 |
| GO:0007165 | signal transduction | 0.003 | 0.003 | PTPRF, MYO6, GABBR2, GRID1, ARHGEF10L, GRID2, FGF14, CDH2, PRDM16, WNT7A, TENM1, ELMO1, ZMIZ1, PLCL1, SH2B2, RCAN2, CHRM5, FCER1G, RBMS3, DCN, TENM4, ULK4, IL1RAPL1, INCENP, CD2AP, NTRK2, SHISA6, ESRRG, NLRX1, ROBO2, ERG, ZFP36L1, RPS6KA2, TCF7L2, PIK3R5, DAB1, DMD, CPE, RBM47, MAGI2, PDE10A, ARHGAP24, FLT3, KIFAP3, TP63, ADCYAP1R1, RASSF8, PREX2, PTPRG, PKHD1, PPARGC1B, PLCD1, PTPRO, CRIM1, PRRT1, CNTN1, LTBP1, DGKI, CTNND2, PTPRT, ARHGAP19, RASGRF2, CTNNAL1, JAK3, NCAM1, CBLB, MAP2K6, CNKSR2, PEX5L, CDH13, ARHGEF10. ANK2, MARVELD3, ASPH, CADM1, DAAM2, GRIN2B, BCR, SLIT3, PKD1, MZB1, ROR2, SEMA6A, CHST11, CHL1, TRPV1, NOTCH2, GABRB3, GABRB1, RELN, TCF7, COL1A1, DLG5, FCHO1, HSPG2, NRG4, EPHA6, DISC1, SYNGAP1, NRXN1, AKT3, MAP3K12, PROX1, DNER, RGL1, PIK3C2G, RORC, RTN4R, PARD3, SPTBN1, PLXND1, CDT1, FOXL1, CCR10. FBN2, DOCK5, FMOD, FGD4, RRH, FGF22, SKI, CCDC88A, FNIP2, MGLL, RAPGEFL1, PLCB2, WWC2, PDGFD, HTR1F, VIPR1, FRMPD1, LRRN2, AUTS2, GLRA1, LCK, CAMK1, EPHA7, FZD3, CELSR1, PPP3CA, BMPR1B, MTSS1, ADAM8, ERBB3, ADCY5, SH2D3C, PCNT, COL4A2, BICD1, DST, EPHB2, REM1, TUFT1, EPHB1, ANGPT2, PRKAG2, ADAM12, ZNF385B, SAMD14, EFNB2, NKX2-2, ADORA2A, KCNIP2, GAL, PIK3C2B, CAMKMT, NSDHL, RAPGEF5, MLLT3 |
| GO:0035295 | tube development | 0.004 | 0.004 | CSMD1, PLXDC1, CDH2, WNT7A, ZMIZ1, DCN, NTRK2, E2F7, ROBO2, ZFP36L1, ADAMTS2, ARHGAP24, TP63, WT1, RAB3A, GLUL, CLMP, PKHD1, CDH13, EMCN, PKD1, IRX1, SEMA6A, NOTCH2, DLG5, SDCCAG8, HSPG2, NRXN1, AKT3, PROX1, CHD7, PLXND1, FOXL1, SKI, SIM2, TBX5, EPHA7, FZD3, CELSR1, MTSS1, ADAM8, PBX1, COL4A2, EPHB2, EPHB1, ANGPT2, ADAM12, TSPAN18, EFNB2, NKX2-2, ZFPM2, STAB2 |
| GO:0070244 | negative regulation of thymocyte apoptotic process | 0.004 | 0.004 | KIFAP3, JAK3, RORC, BCL11B |
| GO:0021684 | cerebellar granular layer formation | 0.004 | 0.004 | GRID2, WNT7A, NRXN1, PROX1 |
| GO:0021707 | cerebellar granule cell differentiation | 0.004 | 0.004 | GRID2, WNT7A, NRXN1, PROX1 |
| GO:0001944 | vasculature development | 0.004 | 0.004 | PLXDC1, CDH2, WNT7A, ZMIZ1, DCN, NTRK2, E2F7, ROBO2, ZFP36L1, TCF7L2, ARHGAP24, WT1, GLUL, CDH13, LAMA4, EMCN, PKD1, SEMA6A, NOTCH2, COL1A1, HSPG2, NRXN1, AKT3, PROX1, CHD7, PLXND1, PDGFD, TBX5, ADAM8, COL4A2, EPHB2, EPHB1, ANGPT2, ADAM12, TSPAN18, EFNB2, ZFPM2, NSDHL, STAB2 |
| GO:0001964 | startle response | 0.005 | 0.005 | CSMD1, GRID2, NRXN1, CTNNA2, GLRA1, ADORA2A |
| GO:0016339 | calcium-dependent cell-cell adhesion via plasma membrane cell adhesion molecules | 0.005 | 0.005 | CDH18, CDH2, CDH12, KIFAP3, CDH13, NRXN1 |
| GO:2000026 | regulation of multicellular organismal development | 0.005 | 0.005 | GRID2, WNT7A, ZMIZ1, CLSTN2, DCN, TENM4, IL1RAPL1, NTRK2, ROBO2, LRTM2, ZFP36L1, DAB1, KIFAP3, TP63, WT1, GLUL, PPARGC1B, JAK3, LAMA4, DAAM2, ROR2, SEMA6A, NOTCH2, RELN, TCF7, DLG5, DBN1, HSPG2, DISC1, SYNGAP1, NRXN1, AKT3, PROX1, BCL11A, RTN4R, PARD3, CHD7, PLXND1, FBN2, VAX1, SKI, TRPS1, CDKL5, TBX5, EPHA7, FZD3, PPP3CA, BMPR1B, ADAM8, COL4A2, EPHB2, EPHB1, ANGPT2, ADAM12, MAP2, TSPAN18, NKX2-2, GAL, ZFPM2 |
| GO:0048513 | animal organ development | 0.005 | 0.005 | FOXP2, MYO6, CNTN5, CSMD1, GRID2, CDH2, WNT7A, ZMIZ1, MACROD2, USH2A, DCN, TENM4, PPARGC1A, SOX21, CD2AP, NTRK2, E2F7, TCOF1, AFF2, ROBO2, MAP1S, ZFP36L1, SDK1, TCF7L2, DAB1, DMD, CPE, TTC39C, MAGI2, ADAMTS2, A2M, ABI3BP, FLT3, TP63, WT1, RAB3A, PAX5, PKHD1, PPARGC1B, PTPRO, CNTN1, NCAM1, MAP2K6, ANK2, CADM1, GRIN2B, BCR, SLIT3, PKD1, GCNT4, IRX1, STOX2, ROR2, SEMA6A, CHST11, NOTCH2, RELN, COL1A1, DLG5, HSPG2, DISC1, NRXN1, AKT3, PROX1, DNER, CPAMD8, RORC, ZFHX3, BCL11B, RTN4R, CHD7, PLXND1, FOXL1, FBN2, CTNNA2, VAX1, FGF22, SKI, TRPS1, SIM2, PDGFD, LDB3, UTRN, TBX5, NEBL, EPHA7, SGCA, KAZN, FZD3, CELSR1, PPP3CA, CACNA1G, BMPR1B, MTSS1, ADAM8, ERBB3, MNAT1, PBX1, EPHB2, TUFT1, EPHB1, ANGPT2, ZBTB40. CHD2, EFNB2, NKX2-2, GAL, ZFPM2, NSDHL |
| GO:0007264 | small GTPase-mediated signal transduction | 0.005 | 0.005 | ARHGEF10L, ELMO1, SH2B2, CD2AP, DAB1, ARHGAP24, ADCYAP1R1, PREX2, DGKI, ARHGAP19, RASGRF2, CTNNAL1, CDH13, ARHGEF10. BCR, NOTCH2, RELN, SYNGAP1, RGL1, RTN4R, DOCK5, FGD4, CCDC88A, RAPGEFL1, AUTS2, CELSR1, SH2D3C, EPHB2, RAPGEF5 |
| GO:0007157 | heterophilic cell-cell adhesion via plasma membrane cell adhesion molecules | 0.005 | 0.005 | GRID2, CDH2, TENM1, TENM4, CADM1, NRXN1, SCARF1 |
| GO:0070242 | thymocyte apoptotic process | 0.006 | 0.006 | KIFAP3, JAK3, RORC, BCL11B, ADAM8 |
| GO:0060078 | regulation of postsynaptic membrane potential | 0.006 | 0.006 | GRID1, GRID2, WNT7A, DGKI, GRIN2B, TRPV1, GABRB1, RELN, NRXN1, GLRA1, PPP3CA, GABRA2, ADORA2A |
| GO:0031345 | negative regulation of cell projection organization | 0.007 | 0.007 | DNM3, DAB1, ARHGAP24, PTPRG, PTPRO, GRIN2B, SYNGAP1, NRXN1, BCL11A, RTN4R, EPHA7, PPP3CA, EPHB2, MAP2, EFNB2 |
| GO:0060999 | positive regulation of dendritic spine development | 0.008 | 0.008 | IL1RAPL1, RELN, DLG5, DBN1, CDKL5, CAMK1, EPHB2 |
| GO:0050789 | regulation of biological process | 0.008 | 0.008 | PTPRF, FOXP2, MYO6, GABBR2, GRID1, ARHGEF10L, DLGAP1, GRID2, FGF14, CDH2, PRDM16, CAMK1D, WNT7A, NAV3, SUPT3H, KLF12, TENM1, ELMO1, DNM3, NOVA1, PANX3, ZMIZ1, KCNIP4, PLCL1, SH2B2, ZNF608, PTPRN2, USH2A, RCAN2, CHRM5, CLSTN2, FCER1G, RBMS3, DCN, TENM4, ULK4, PPARGC1A, RBFOX1, IL1RAPL1, CTTNBP2, CHTF18, SOX21, INCENP, CD2AP, RAD51B, NTRK2, SHISA6, NBEA, E2F7, DPP10. ESRRG, DACH1, TCOF1, TTC39B, NLRX1, CACNA1B, SLC4A3, AFF2, CREB5, ROBO2, DLGAP2, SLCO3A1, LRTM2, MAP1S, NKAIN2, ERG, TDRD9, ZNF362, ZFP36L1, SDK1, RPS6KA2, TCF7L2, PIK3R5, MBNL3, DAB1, MSI2, DMD, CPE, RBM47, MAGI2, A2M, ABI3BP, PDE10A, EDIL3, ARHGAP24, FLT3, KIFAP3, TP63, ADCYAP1R1, SPON1, RASSF8, WT1, RAB3A, PREX2, PTPRG, GLUL, CLMP, PAX5, TOX2, PKHD1, SHMT1, MRPL12, PPARGC1B, PLCD1, ASTN2, PTPRO, CRIM1, PRRT1, CNTN1, KIF1A, LTBP1, DGKI, CTNND2, PTPRT, ARHGAP19, RASGRF2, CTNNAL1, JAK3, NCAM1, SLC4A11, CBLB, MAP2K6, CNKSR2, PEX5L, EBF1, KIF18B, CDH13, ARHGEF10. ANK2, MARVELD3, ASPH, LAMA4, CADM1, DAAM2, GRIN2B, BCR, EMCN, TANC2, SLIT3, PKD1, MZB1, IRX1, STOX2, ROR2, SEMA6A, PFKL, CHST11, CHL1, TRPV1, NOTCH2, GABRB3, GABRB1, RELN, TCF7, STX18, CACNB2, COL1A1, DLG5, DBN1, FCHO1, ZNF521, SDCCAG8, HSPG2, NRG4, CERS1, EPHA6, DISC1, SYNGAP1, NRXN1, AKT3, TWIST2, MAP3K12, PROX1, DNER, ZDHHC15, BCL11A, RGL1, PIK3C2G, ACAP3, RORC, ZFHX3, ELOVL5, BCL11B, RTN4R, PARD3, TNKS1BP1, CHD7, SPTBN1, PLXND1, CDT1, FKBP6, KIF13A, PKNOX2, FOXL1, CCR10. FBN2, DOCK5, XPO4, FMOD, CTNNA2, VAX1, FGD4, RRH, FGF22, SKI, CCDC88A, FNIP2, ATP13A2, TRPS1, MTA3, LSM11, RILP, MGLL, RAPGEFL1, ZNF618, SIM2, PLCB2, CDKL5, WWC2, PDGFD, CTNNA3, HTR1F, VIPR1, FRMPD1, LRRN2, AUTS2, GLRA1, LCK, UTRN, TBX5, CAMK1, EPHA7, APBA1, SGCA, FZD3, CELSR1, TNNC2, PPP3CA, NSUN3, ONECUT3, CACNA1G, BMPR1B, MTSS1, ADAM8, ERBB3, ADCY5, SH2D3C, BSN, CPLX2, MNAT1, PCNT, PBX1, COL4A2, SCARF1, BICD1, DST, EPHB2, REM1, TUFT1, MYH10. FRMD4A, EPHB1, ANGPT2, PRKAG2, ADAM12, SACS, GABRA2, RIOK1, LRCH1, MAP2, ZBTB40. ZNF385B, SAMD14, TSPAN18, UBE2Z, EFNB2, NKX2-2, ADORA2A, RBBP5, KCNIP2, GAL, ZFPM2, ZFHX4, PIK3C2B, CAMKMT, NSDHL, ABCG4, RAPGEF5, MLLT3 |
| GO:0141124 | intracellular signaling cassette | 0.008 | 0.008 | ARHGEF10L, FGF14, CDH2, WNT7A, TENM1, ELMO1, ZMIZ1, SH2B2, RCAN2, DCN, ULK4, CD2AP, NTRK2, NLRX1, ZFP36L1, TCF7L2, PIK3R5, DAB1, DMD, MAGI2, PDE10A, ARHGAP24, FLT3, ADCYAP1R1, PREX2, PKHD1, DGKI, ARHGAP19, RASGRF2, CTNNAL1, MAP2K6, PEX5L, CDH13, ARHGEF10. ANK2, MARVELD3, ASPH, GRIN2B, BCR, ROR2, SEMA6A, TRPV1, NOTCH2, RELN, SYNGAP1, NRXN1, MAP3K12, RGL1, PIK3C2G, RTN4R, SPTBN1, CCR10. DOCK5, FGD4, FGF22, SKI, CCDC88A, RAPGEFL1, PDGFD, AUTS2, EPHA7, CELSR1, PPP3CA, ADAM8, ERBB3, SH2D3C, EPHB2, REM1, EPHB1, SAMD14, GAL, PIK3C2B, RAPGEF5 |
| GO:0042391 | regulation of membrane potential | 0.009 | 0.009 | GRID1, KCNK17, GRID2, WNT7A, DCN, NTRK2, SLC4A3, DMD, DGKI, SLC4A11, ANK2, GRIN2B, TRPV1, GABRB3, GABRB1, RELN, CACNB2, NRXN1, CTNNA3, GLRA1, TBX5, PPP3CA, CACNA1G, GABRA2, ADORA2A, KCNIP2 |
| GO:0050793 | regulation of developmental process | 0.009 | 0.009 | PTPRF, GRID2, CAMK1D, WNT7A, DNM3, ZMIZ1, USH2A, CLSTN2, DCN, TENM4, IL1RAPL1, NTRK2, ROBO2, LRTM2, ZFP36L1, SDK1, TCF7L2, MBNL3, DAB1, DMD, ABI3BP, KIFAP3, TP63, WT1, GLUL, PKHD1, PPARGC1B, CRIM1, KIF1A, JAK3, SLC4A11, LAMA4, DAAM2, BCR, TANC2, ROR2, SEMA6A, TRPV1, NOTCH2, RELN, TCF7, COL1A1, DLG5, DBN1, HSPG2, DISC1, SYNGAP1, NRXN1, AKT3, TWIST2, PROX1, ZDHHC15, BCL11A, RORC, ZFHX3, BCL11B, RTN4R, PARD3, CHD7, PLXND1, FBN2, VAX1, FGD4, SKI, TRPS1, MTA3, CDKL5, WWC2, TBX5, CAMK1, EPHA7, FZD3, PPP3CA, BMPR1B, ADAM8, ERBB3, PBX1, COL4A2, SCARF1, EPHB2, MYH10. EPHB1, ANGPT2, ADAM12, MAP2, TSPAN18, EFNB2, NKX2-2, GAL, ZFPM2 |
| GO:0032835 | glomerulus development | 0.009 | 0.009 | CD2AP, MAGI2, WT1, PTPRO, NOTCH2, PDGFD, PPP3CA, MTSS1, ANGPT2 |
| GO:0021683 | cerebellar granular layer morphogenesis | 0.009 | 0.009 | GRID2, WNT7A, NRXN1, PROX1 |
| GO:0051963 | regulation of synapse assembly | 0.010 | 0.010 | GRID2, WNT7A, ELMO1, CLSTN2, IL1RAPL1, NTRK2, ROBO2, LRTM2, DLG5, NRXN1, RTN4R, EPHA7, EPHB2, EPHB1 |
| GO:0050770 | regulation of axonogenesis | 0.011 | 0.011 | WNT7A, NTRK2, ROBO2, DAB1, DISC1, SYNGAP1, BCL11A, RTN4R, PLXND1, CDKL5, EPHA7, EPHB2, MAP2 |
| GO:0051962 | positive regulation of nervous system development | 0.011 | 0.011 | GRID2, WNT7A, CLSTN2, TENM4, IL1RAPL1, NTRK2, ROBO2, LRTM2, RELN, DLG5, DBN1, DISC1, NRXN1, BCL11A, PLXND1, CDKL5, FZD3, EPHB2, EPHB1, NKX2-2 |
| GO:1901890 | positive regulation of cell junction assembly | 0.011 | 0.011 | GRID2, WNT7A, CLSTN2, IL1RAPL1, NTRK2, LRTM2, DLG5, NRXN1, TBX5, EPHB2, EPHB1 |
| GO:0021782 | glial cell development | 0.012 | 0.012 | TENM4, NTRK2, DMD, CNTN1, ARHGEF10. ROR2, PARD3, MXRA8, SKI, ERBB3, NKX2-2, ADORA2A |
| GO:0061318 | renal filtration cell differentiation | 0.012 | 0.012 | CD2AP, MAGI2, WT1, PTPRO, NOTCH2 |
| GO:0072112 | podocyte differentiation | 0.012 | 0.012 | CD2AP, MAGI2, WT1, PTPRO, NOTCH2 |
| GO:0051241 | negative regulation of multicellular organismal process | 0.012 | 0.012 | WNT7A, NAV3, NOVA1, PLCL1, DCN, CD2AP, NLRX1, ROBO2, DAB1, ABI3BP, KIFAP3, TP63, WT1, PTPRG, PTPRO, JAK3, CBLB, MAP2K6, MARVELD3, LAMA4, DAAM2, BCR, SEMA6A, TRPV1, DLG5, HSPG2, SYNGAP1, PROX1, BCL11A, RTN4R, DOCK5, VAX1, SKI, WWC2, GLRA1, TBX5, EPHA7, PPP3CA, COL4A2, EPHB2, ANGPT2, MAP2, ADORA2A, GAL, ZFPM2 |
| GO:0060322 | head development | 0.013 | 0.013 | FOXP2, CNTN5, GRID2, CDH2, WNT7A, ZMIZ1, MACROD2, SOX21, NTRK2, AFF2, ROBO2, MAP1S, DAB1, PAX5, CNTN1, ASPH, GRIN2B, BCR, RELN, COL1A1, DLG5, HSPG2, DISC1, NRXN1, AKT3, PROX1, BCL11B, RTN4R, CHD7, CTNNA2, VAX1, SKI, EPHA7, FZD3, MNAT1, PBX1, EPHB2, EPHB1, NKX2-2 |
| GO:0065007 | biological regulation | 0.014 | 0.014 | PTPRF, FOXP2, MYO6, GABBR2, GRID1, ARHGEF10L, KCNK17, DLGAP1, GRID2, FGF14, CDH2, PRDM16, CAMK1D, WNT7A, NAV3, SUPT3H, KLF12, TENM1, ELMO1, DNM3, NOVA1, PANX3, ZMIZ1, KCNIP4, PLCL1, SH2B2, ZNF608, PTPRN2, USH2A, RCAN2, CHRM5, CLSTN2, FCER1G, RBMS3, DCN, TENM4, ULK4, PPARGC1A, RBFOX1, IL1RAPL1, CTTNBP2, CHTF18, SOX21, INCENP, CD2AP, RAD51B, NTRK2, SHISA6, NBEA, E2F7, DPP10. ESRRG, DACH1, TCOF1, TTC39B, NLRX1, CACNA1B, SLC4A3, AFF2, CREB5, ROBO2, DLGAP2, SLCO3A1, LRTM2, MAP1S, NKAIN2, ERG, TDRD9, ZNF362, ZFP36L1, SDK1, RPS6KA2, TCF7L2, PIK3R5, MBNL3, DAB1, MSI2, DMD, CPE, RBM47, MCM8, MAGI2, A2M, ABI3BP, PDE10A, EDIL3, ARHGAP24, FLT3, KIFAP3, TP63, ADCYAP1R1, SPON1, RASSF8, WT1, RAB3A, PREX2, PTPRG, GLUL, CLMP, PAX5, TOX2, PKHD1, SHMT1, MRPL12, PPARGC1B, PLCD1, ASTN2, PTPRO, CRIM1, PRRT1, CNTN1, KIF1A, LTBP1, DGKI, CTNND2, PTPRT, ARHGAP19, RASGRF2, CTNNAL1, JAK3, NCAM1, SLC4A11, AGAP3, CBLB, MAP2K6, CNKSR2, PEX5L, EBF1, KIF18B, CDH13, ARHGEF10. ANK2, MARVELD3, ASPH, LAMA4, CADM1, DAAM2, GRIN2B, BCR, EMCN, TANC2, SLIT3, PKD1, MZB1, GCNT4, IRX1, STOX2, ROR2, SEMA6A, PFKL, CHST11, CHL1, TRPV1, NOTCH2, GABRB3, GABRB1, RELN, TCF7, STX18, CACNB2, COL1A1, DLG5, DBN1, FCHO1, ZNF521, SDCCAG8, HSPG2, NRG4, CERS1, EPHA6, DISC1, SYNGAP1, NRXN1, AKT3, TWIST2, MAP3K12, PROX1, DNER, ZDHHC15, BCL11A, RGL1, PIK3C2G, ACAP3, RORC, ZFHX3, ELOVL5, BCL11B, RTN4R, PARD3, TNKS1BP1, CHD7, SPTBN1, PLXND1, CDT1, FKBP6, KIF13A, PKNOX2, FOXL1, CCR10. FBN2, DOCK5, XPO4, FMOD, CTNNA2, VAX1, FGD4, RRH, FGF22, SKI, CCDC88A, FNIP2, ATP13A2, TRPS1, MTA3, LSM11, RILP, MGLL, RAPGEFL1, ZNF618, SIM2, PLCB2, CDKL5, WWC2, PDGFD, CTNNA3, HTR1F, VIPR1, FRMPD1, LRRN2, AUTS2, GLRA1, LCK, UTRN, TBX5, CAMK1, EPHA7, APBA1, SGCA, FZD3, CELSR1, TNNC2, PPP3CA, NSUN3, ONECUT3, CACNA1G, BMPR1B, MTSS1, ADAM8, ERBB3, ADCY5, SH2D3C, BSN, CPLX2, MNAT1, PCNT, PBX1, COL4A2, SCARF1, BICD1, DST, EPHB2, REM1, TUFT1, MYH10. FRMD4A, EPHB1, ANGPT2, PRKAG2, ADAM12, SACS, GABRA2, RIOK1, LRCH1, NHEJ1, MAP2, ZBTB40. ZNF385B, SAMD14, TSPAN18, UBE2Z, EFNB2, NKX2-2, ADORA2A, RBBP5, KCNIP2, GAL, ZFPM2, ZFHX4, PIK3C2B, CAMKMT, NSDHL, ABCG4, RAPGEF5, SLC9A9, MLLT3 |
| GO:0007420 | brain development | 0.014 | 0.014 | FOXP2, CNTN5, GRID2, CDH2, WNT7A, ZMIZ1, MACROD2, SOX21, NTRK2, AFF2, ROBO2, MAP1S, DAB1, PAX5, CNTN1, GRIN2B, BCR, RELN, DLG5, HSPG2, DISC1, NRXN1, AKT3, PROX1, BCL11B, RTN4R, CHD7, CTNNA2, VAX1, SKI, EPHA7, FZD3, MNAT1, PBX1, EPHB2, EPHB1, NKX2-2 |
| GO:0072359 | circulatory system development | 0.014 | 0.014 | PLXDC1, CDH2, WNT7A, ZMIZ1, DCN, TENM4, NTRK2, E2F7, ROBO2, ZFP36L1, TCF7L2, CPE, ABI3BP, ARHGAP24, WT1, GLUL, CDH13, ANK2, LAMA4, EMCN, SLIT3, PKD1, SEMA6A, NOTCH2, COL1A1, HSPG2, NRXN1, AKT3, PROX1, CHD7, PLXND1, FOXL1, SKI, PDGFD, LDB3, TBX5, NEBL, CACNA1G, ADAM8, ERBB3, MNAT1, COL4A2, EPHB2, EPHB1, ANGPT2, ADAM12, TSPAN18, EFNB2, ZFPM2, NSDHL, STAB2 |
| GO:0072311 | glomerular epithelial cell differentiation | 0.015 | 0.015 | CD2AP, MAGI2, WT1, PTPRO, NOTCH2 |
| GO:0097553 | calcium ion transmembrane import into cytosol | 0.015 | 0.015 | CACNA1B, DMD, ADCYAP1R1, PLCD1, ANK2, ASPH, GRIN2B, TRPV1, CHD7, PLCB2, LCK, PPP3CA, CACNA1G |
| GO:0060079 | excitatory postsynaptic potential | 0.015 | 0.015 | GRID2, WNT7A, DGKI, GRIN2B, TRPV1, RELN, NRXN1, GLRA1, PPP3CA, ADORA2A |
| GO:0048646 | anatomical structure formation involved in morphogenesis | 0.016 | 0.016 | GRID2, PLXDC1, WNT7A, DCN, TENM4, E2F7, TCOF1, ROBO2, SDK1, DAB1, ARHGAP24, TP63, WT1, GLUL, PPARGC1B, CNTN1, CDH13, ANK2, EMCN, IRX1, SEMA6A, NOTCH2, RELN, COL1A1, SDCCAG8, HSPG2, NRXN1, AKT3, PROX1, SPTBN1, PLXND1, FBN2, DOCK5, SKI, LDB3, TBX5, NEBL, CAMK1, FZD3, CELSR1, ADAM8, COL4A2, EPHB2, EPHB1, ANGPT2, ADAM12, TSPAN18, EFNB2, NBEAL2, SH3PXD2A, STAB2 |
| GO:0048167 | regulation of synaptic plasticity | 0.016 | 0.016 | GRID2, CD2AP, NTRK2, SHISA6, RAB3A, PRRT1, DGKI, RASGRF2, GRIN2B, RELN, DBN1, SYNGAP1, CPLX2, EPHB2, ADORA2A |
| GO:0050771 | negative regulation of axonogenesis | 0.016 | 0.016 | DAB1, SYNGAP1, BCL11A, RTN4R, EPHA7, EPHB2, MAP2 |
| GO:0048514 | blood vessel morphogenesis | 0.016 | 0.016 | PLXDC1, CDH2, WNT7A, ZMIZ1, DCN, NTRK2, E2F7, ZFP36L1, ARHGAP24, WT1, GLUL, CDH13, EMCN, SEMA6A, NOTCH2, HSPG2, NRXN1, AKT3, PROX1, CHD7, PLXND1, TBX5, ADAM8, COL4A2, EPHB2, EPHB1, ANGPT2, ADAM12, TSPAN18, EFNB2, ZFPM2, STAB2 |
| GO:0060134 | prepulse inhibition | 0.017 | 0.017 | GRID2, NRXN1, CTNNA2, ADORA2A |
| GO:0043491 | phosphatidylinositol 3-kinase/protein kinase B signal transduction | 0.017 | 0.017 | DCN, CD2AP, NTRK2, ZFP36L1, TCF7L2, PIK3R5, MAGI2, FLT3, PREX2, PKHD1, ROR2, RELN, NRXN1, PIK3C2G, CCDC88A, PDGFD, ADAM8, ERBB3, PIK3C2B |
| GO:0031346 | positive regulation of cell projection organization | 0.017 | 0.017 | CAMK1D, TENM1, DNM3, IL1RAPL1, NTRK2, ROBO2, DMD, MAGI2, CNTN1, ROR2, RELN, DBN1, DISC1, ZDHHC15, BCL11A, PLXND1, CCDC88A, CDKL5, AUTS2, CAMK1, SCARF1, EPHB2 |
| GO:0010001 | glial cell differentiation | 0.018 | 0.018 | CDH2, TENM4, NTRK2, DAB1, DMD, CNTN1, ARHGEF10. DAAM2, ROR2, RELN, DNER, PARD3, MXRA8, VAX1, SKI, ERBB3, NKX2-2, ADORA2A |
| GO:0050794 | regulation of cellular process | 0.018 | 0.018 | PTPRF, FOXP2, MYO6, GABBR2, GRID1, ARHGEF10L, GRID2, FGF14, CDH2, PRDM16, CAMK1D, WNT7A, NAV3, SUPT3H, KLF12, TENM1, ELMO1, DNM3, NOVA1, PANX3, ZMIZ1, KCNIP4, PLCL1, SH2B2, ZNF608, USH2A, RCAN2, CHRM5, CLSTN2, FCER1G, RBMS3, DCN, TENM4, ULK4, PPARGC1A, RBFOX1, IL1RAPL1, CTTNBP2, CHTF18, SOX21, INCENP, CD2AP, RAD51B, NTRK2, SHISA6, NBEA, E2F7, DPP10. ESRRG, DACH1, TCOF1, TTC39B, NLRX1, CACNA1B, AFF2, CREB5, ROBO2, SLCO3A1, LRTM2, MAP1S, ERG, TDRD9, ZNF362, ZFP36L1, RPS6KA2, TCF7L2, PIK3R5, MBNL3, DAB1, MSI2, DMD, CPE, RBM47, MAGI2, ABI3BP, PDE10A, EDIL3, ARHGAP24, FLT3, KIFAP3, TP63, ADCYAP1R1, SPON1, RASSF8, WT1, RAB3A, PREX2, PTPRG, GLUL, CLMP, PAX5, TOX2, PKHD1, SHMT1, MRPL12, PPARGC1B, PLCD1, ASTN2, PTPRO, CRIM1, PRRT1, CNTN1, KIF1A, LTBP1, DGKI, CTNND2, PTPRT, ARHGAP19, RASGRF2, CTNNAL1, JAK3, NCAM1, SLC4A11, CBLB, MAP2K6, CNKSR2, PEX5L, EBF1, KIF18B, CDH13, ARHGEF10. ANK2, MARVELD3, ASPH, LAMA4, CADM1, DAAM2, GRIN2B, BCR, EMCN, TANC2, SLIT3, PKD1, MZB1, IRX1, STOX2, ROR2, SEMA6A, PFKL, CHST11, CHL1, TRPV1, NOTCH2, GABRB3, GABRB1, RELN, TCF7, STX18, CACNB2, COL1A1, DLG5, DBN1, FCHO1, ZNF521, SDCCAG8, HSPG2, NRG4, CERS1, EPHA6, DISC1, SYNGAP1, NRXN1, AKT3, TWIST2, MAP3K12, PROX1, DNER, ZDHHC15, BCL11A, RGL1, PIK3C2G, ACAP3, RORC, ZFHX3, ELOVL5, BCL11B, RTN4R, PARD3, TNKS1BP1, CHD7, SPTBN1, PLXND1, CDT1, FKBP6, KIF13A, PKNOX2, FOXL1, CCR10. FBN2, DOCK5, XPO4, FMOD, CTNNA2, VAX1, FGD4, RRH, FGF22, SKI, CCDC88A, FNIP2, ATP13A2, TRPS1, MTA3, LSM11, RILP, MGLL, RAPGEFL1, ZNF618, SIM2, PLCB2, CDKL5, WWC2, PDGFD, CTNNA3, HTR1F, VIPR1, FRMPD1, LRRN2, AUTS2, GLRA1, LCK, UTRN, TBX5, CAMK1, EPHA7, APBA1, FZD3, CELSR1, PPP3CA, NSUN3, ONECUT3, CACNA1G, BMPR1B, MTSS1, ADAM8, ERBB3, ADCY5, SH2D3C, BSN, CPLX2, MNAT1, PCNT, PBX1, COL4A2, SCARF1, BICD1, DST, EPHB2, REM1, TUFT1, MYH10. FRMD4A, EPHB1, ANGPT2, PRKAG2, ADAM12, SACS, RIOK1, LRCH1, MAP2, ZBTB40. ZNF385B, SAMD14, UBE2Z, EFNB2, NKX2-2, ADORA2A, RBBP5, KCNIP2, GAL, ZFPM2, ZFHX4, PIK3C2B, CAMKMT, NSDHL, ABCG4, RAPGEF5, MLLT3 |
| GO:0045216 | cell-cell junction organization | 0.021 | 0.021 | CDH18, CDH2, CD2AP, CDH12, PKHD1, PTPRO, CDH13, ANK2, MARVELD3, DLG5, PARD3, TBX5, MTSS1, EPHB2, EFNB2 |
| GO:0097475 | motor neuron migration | 0.023 | 0.023 | DAB1, RELN, FZD3, CELSR1 |
| GO:0140650 | radial glia-guided pyramidal neuron migration | 0.023 | 0.023 | ZMIZ1, DAB1, DISC1 |
| GO:0021953 | central nervous system neuron differentiation | 0.024 | 0.024 | GRID2, MDGA2, WNT7A, ZMIZ1, NTRK2, ROBO2, GABRB1, DISC1, NRXN1, PROX1, BCL11B, BMPR1B, EPHB2, EPHB1, MAP2, NKX2-2 |
| GO:0072010 | glomerular epithelium development | 0.026 | 0.026 | CD2AP, MAGI2, WT1, PTPRO, NOTCH2 |
| GO:0046660 | female sex differentiation | 0.026 | 0.026 | CSMD1, DACH1, ROBO2, A2M, TP63, ADCYAP1R1, WT1, SLIT3, CHD7, BMPR1B, ZFPM2 |
| GO:0099068 | postsynapse assembly | 0.027 | 0.027 | GRID2, CDH2, WNT7A, ELMO1, CSMD2, RELN, NRXN1, RTN4R, EPHB2 |
| GO:0099084 | postsynaptic specialization organization | 0.028 | 0.028 | GRID2, CDH2, CSMD2, CNKSR2, RELN, SYNGAP1, NRXN1 |
| GO:0010977 | negative regulation of neuron projection development | 0.028 | 0.028 | DNM3, DAB1, PTPRG, PTPRO, SYNGAP1, BCL11A, RTN4R, EPHA7, EPHB2, MAP2, EFNB2 |
| GO:0051716 | cellular response to stimulus | 0.028 | 0.028 | PTPRF, MYO6, GABBR2, GRID1, ARHGEF10L, GRID2, FGF14, CDH2, PRDM16, CAMK1D, WNT7A, SUPT3H, TENM1, ELMO1, ZMIZ1, PLCL1, SH2B2, PTPRN2, MACROD2, RCAN2, CHRM5, FCER1G, RBMS3, DCN, TENM4, ULK4, PPARGC1A, IL1RAPL1, INCENP, CD2AP, RAD51B, NTRK2, SHISA6, E2F7, ESRRG, NLRX1, ROBO2, ERG, ZFP36L1, RPS6KA2, TCF7L2, PIK3R5, DAB1, DMD, CPE, RBM47, MCM8, MAGI2, PDE10A, ARHGAP24, FLT3, KIFAP3, TP63, ADCYAP1R1, RASSF8, WT1, PREX2, PTPRG, GLUL, PKHD1, SHMT1, PPARGC1B, PLCD1, PTPRO, CRIM1, PRRT1, CNTN1, LTBP1, DGKI, CTNND2, PTPRT, ARHGAP19, RASGRF2, CTNNAL1, JAK3, NCAM1, SLC4A11, AGAP3, CBLB, MAP2K6, CNKSR2, PEX5L, CDH13, ARHGEF10. ANK2, MARVELD3, ASPH, CADM1, DAAM2, GRIN2B, BCR, SLIT3, PKD1, MZB1, ROR2, SEMA6A, CCDC13, CHST11, CHL1, TRPV1, NOTCH2, GABRB3, GABRB1, RELN, TCF7, COL1A1, DLG5, FCHO1, HSPG2, NRG4, CERS1, EPHA6, DISC1, SYNGAP1, NRXN1, AKT3, MAP3K12, PROX1, DNER, BCL11A, RGL1, PIK3C2G, RORC, RTN4R, PARD3, TNKS1BP1, SPTBN1, PLXND1, CDT1, FOXL1, CCR10. FBN2, DOCK5, FMOD, FGD4, RRH, FGF22, SKI, CCDC88A, FNIP2, ATP13A2, MGLL, RAPGEFL1, PLCB2, WWC2, PDGFD, SCARA5, HTR1F, VIPR1, FRMPD1, LRRN2, AUTS2, GLRA1, LCK, CAMK1, EPHA7, FZD3, CELSR1, PPP3CA, BMPR1B, MTSS1, ADAM8, ERBB3, ADCY5, SH2D3C, MNAT1, PCNT, COL4A2, SCARF1, BICD1, DST, EPHB2, REM1, TUFT1, EPHB1, ANGPT2, PRKAG2, ADAM12, NHEJ1, ZBTB40. ZNF385B, CHD2, SAMD14, EFNB2, NKX2-2, ADORA2A, RBBP5, KCNIP2, GAL, PIK3C2B, CAMKMT, NSDHL, ABCG4, RAPGEF5, MLLT3 |
| GO:0021681 | cerebellar granular layer development | 0.028 | 0.028 | GRID2, WNT7A, NRXN1, PROX1 |
| GO:0097091 | synaptic vesicle clustering | 0.028 | 0.028 | CDH2, RAB3A, NRXN1, BSN |
| GO:0034332 | adherens junction organization | 0.030 | 0.030 | CDH18, CDH2, CDH12, CDH13, DLG5, MTSS1, EFNB2 |
| GO:0007167 | enzyme-linked receptor protein signaling pathway | 0.030 | 0.030 | PTPRF, PRDM16, ZMIZ1, SH2B2, DCN, CD2AP, NTRK2, MAGI2, FLT3, PTPRG, CRIM1, LTBP1, PTPRT, JAK3, CBLB, CDH13, CADM1, MZB1, ROR2, SEMA6A, CHST11, NOTCH2, COL1A1, NRG4, EPHA6, NRXN1, SPTBN1, FBN2, FMOD, FGF22, SKI, CCDC88A, PDGFD, LCK, EPHA7, BMPR1B, MTSS1, ERBB3, COL4A2, EPHB2, EPHB1, ANGPT2, EFNB2 |
| GO:0099565 | chemical synaptic transmission, postsynaptic | 0.030 | 0.030 | GRID2, WNT7A, DGKI, GRIN2B, TRPV1, RELN, NRXN1, GLRA1, PPP3CA, ADORA2A |
| GO:0001822 | kidney development | 0.030 | 0.030 | CD2AP, ROBO2, MAGI2, WT1, PKHD1, PTPRO, PKD1, GCNT4, IRX1, NOTCH2, DLG5, PROX1, PLXND1, PDGFD, EPHA7, PPP3CA, MTSS1, PBX1, ANGPT2, EFNB2 |
| GO:0007169 | cell surface receptor protein tyrosine kinase signaling pathway | 0.033 | 0.033 | SH2B2, DCN, CD2AP, NTRK2, FLT3, PTPRG, CRIM1, PTPRT, JAK3, CBLB, CDH13, CADM1, MZB1, ROR2, SEMA6A, COL1A1, NRG4, EPHA6, NRXN1, FGF22, CCDC88A, PDGFD, LCK, EPHA7, MTSS1, ERBB3, COL4A2, EPHB2, EPHB1, ANGPT2, EFNB2 |
| GO:0070232 | regulation of T cell apoptotic process | 0.034 | 0.034 | KIFAP3, JAK3, RORC, BCL11B, ADAM8 |
| GO:0009887 | animal organ morphogenesis | 0.034 | 0.034 | MYO6, CSMD1, CDH2, WNT7A, ZMIZ1, DCN, NTRK2, ROBO2, SDK1, CPE, TTC39C, TP63, WT1, PAX5, PKHD1, PPARGC1B, BCR, SLIT3, PKD1, GCNT4, IRX1, SEMA6A, CHST11, NOTCH2, COL1A1, DLG5, AKT3, PROX1, BCL11B, CHD7, PLXND1, FBN2, CTNNA2, FGF22, SKI, TBX5, FZD3, CELSR1, PPP3CA, BMPR1B, PBX1, EPHB2, TUFT1, EPHB1, EFNB2, ZFPM2 |
| GO:0060857 | establishment of glial blood-brain barrier | 0.034 | 0.034 | DMD, MXRA8 |
| GO:0097104 | postsynaptic membrane assembly | 0.034 | 0.034 | CDH2, NRXN1, EPHB2 |
| GO:0097118 | neuroligin clustering involved in postsynaptic membrane assembly | 0.034 | 0.034 | CDH2, NRXN1 |
| GO:1990709 | presynaptic active zone organization | 0.034 | 0.034 | RAB3A, NRXN1, BSN |
| GO:0060856 | establishment of blood-brain barrier | 0.035 | 0.035 | WNT7A, DMD, TRPV1, MXRA8 |
| GO:0010721 | negative regulation of cell development | 0.036 | 0.036 | WNT7A, DAB1, KIFAP3, JAK3, DAAM2, TRPV1, SYNGAP1, PROX1, BCL11A, RTN4R, VAX1, SKI, EPHA7, PPP3CA, EPHB2, MAP2 |
| GO:0007166 | cell surface receptor signaling pathway | 0.037 | 0.037 | PTPRF, GRID1, GRID2, CDH2, PRDM16, WNT7A, ZMIZ1, SH2B2, FCER1G, RBMS3, DCN, CD2AP, NTRK2, SHISA6, ROBO2, TCF7L2, DAB1, CPE, RBM47, MAGI2, FLT3, TP63, ADCYAP1R1, PTPRG, PTPRO, CRIM1, CNTN1, LTBP1, DGKI, CTNND2, PTPRT, JAK3, NCAM1, CBLB, CDH13, CADM1, DAAM2, GRIN2B, SLIT3, PKD1, MZB1, ROR2, SEMA6A, CHST11, TRPV1, NOTCH2, RELN, TCF7, COL1A1, DLG5, FCHO1, HSPG2, NRG4, EPHA6, DISC1, NRXN1, DNER, RTN4R, SPTBN1, PLXND1, FOXL1, CCR10. FBN2, FMOD, FGF22, SKI, CCDC88A, PDGFD, VIPR1, GLRA1, LCK, EPHA7, FZD3, CELSR1, PPP3CA, BMPR1B, MTSS1, ERBB3, ADCY5, COL4A2, DST, EPHB2, EPHB1, ANGPT2, ADAM12, EFNB2, NKX2-2, ADORA2A, NSDHL, MLLT3 |
| GO:0007214 | gamma-aminobutyric acid signaling pathway | 0.039 | 0.039 | GABBR2, PLCL1, GABRB3, GABRB1, GABRA2 |
| GO:0035556 | intracellular signal transduction | 0.042 | 0.042 | MYO6, ARHGEF10L, FGF14, CDH2, WNT7A, TENM1, ELMO1, ZMIZ1, PLCL1, SH2B2, RCAN2, DCN, ULK4, INCENP, CD2AP, NTRK2, ESRRG, NLRX1, ZFP36L1, RPS6KA2, TCF7L2, PIK3R5, DAB1, DMD, MAGI2, PDE10A, ARHGAP24, FLT3, TP63, ADCYAP1R1, PREX2, PKHD1, PPARGC1B, PLCD1, DGKI, ARHGAP19, RASGRF2, CTNNAL1, JAK3, CBLB, MAP2K6, CNKSR2, PEX5L, CDH13, ARHGEF10. ANK2, MARVELD3, ASPH, GRIN2B, BCR, ROR2, SEMA6A, TRPV1, NOTCH2, RELN, DLG5, DISC1, SYNGAP1, NRXN1, AKT3, MAP3K12, PROX1, RGL1, PIK3C2G, RORC, RTN4R, SPTBN1, CDT1, CCR10. DOCK5, FGD4, FGF22, SKI, CCDC88A, FNIP2, RAPGEFL1, PLCB2, WWC2, PDGFD, AUTS2, LCK, CAMK1, EPHA7, CELSR1, PPP3CA, ADAM8, ERBB3, ADCY5, SH2D3C, EPHB2, REM1, TUFT1, EPHB1, PRKAG2, ZNF385B, SAMD14, GAL, PIK3C2B, RAPGEF5 |
| GO:0072001 | renal system development | 0.042 | 0.042 | CD2AP, ROBO2, MAGI2, WT1, PKHD1, PTPRO, PKD1, GCNT4, IRX1, NOTCH2, DLG5, PROX1, PLXND1, PDGFD, EPHA7, PPP3CA, MTSS1, PBX1, ANGPT2, EFNB2 |
| GO:0030155 | regulation of cell adhesion | 0.042 | 0.042 | ZMIZ1, DAB1, DMD, ABI3BP, EDIL3, KIFAP3, PKHD1, PTPRO, JAK3, CBLB, CDH13, LAMA4, EMCN, PKD1, SEMA6A, COL1A1, DLG5, FCHO1, HSPG2, DISC1, ZFHX3, PLXND1, DOCK5, LCK, UTRN, EPHA7, PPP3CA, ADAM8, ERBB3, EPHB2, ANGPT2, EFNB2, ADORA2A |
| GO:0021535 | cell migration in hindbrain | 0.043 | 0.043 | DAB1, CTNNA2, EPHB2, EPHB1 |
| GO:0072073 | kidney epithelium development | 0.044 | 0.044 | CD2AP, ROBO2, MAGI2, WT1, PTPRO, PKD1, IRX1, NOTCH2, EPHA7, MTSS1, PBX1, EFNB2 |
| GO:0051968 | positive regulation of synaptic transmission, glutamatergic | 0.045 | 0.045 | GRIN2B, ROR2, RELN, NRXN1, ADORA2A |
| GO:0050896 | response to stimulus | 0.046 | 0.046 | PTPRF, MYO6, GABBR2, CSMD1, GRID1, ARHGEF10L, GRID2, FGF14, CDH2, PRDM16, CAMK1D, WNT7A, SUPT3H, TENM1, ELMO1, ZMIZ1, PLCL1, SH2B2, HLCS, PTPRN2, MACROD2, RCAN2, CHRM5, FCER1G, RBMS3, DCN, TENM4, ULK4, PPARGC1A, CUTA, IL1RAPL1, CUBN, INCENP, CD2AP, RAD51B, NTRK2, SHISA6, E2F7, ESRRG, DACH1, NLRX1, CACNA1B, ROBO2, LRTM2, ERG, DHX16, ZFP36L1, SDK1, RPS6KA2, TCF7L2, PIK3R5, DAB1, DMD, CPE, RBM47, MCM8, MAGI2, A2M, ABI3BP, PDE10A, ARHGAP24, FLT3, KIFAP3, TP63, ADCYAP1R1, RASSF8, WT1, RAB3A, PREX2, PTPRG, GLUL, PKHD1, SHMT1, PPARGC1B, PLCD1, PTPRO, CRIM1, PRRT1, CNTN1, LTBP1, SCARA3, DGKI, CTNND2, PTPRT, ARHGAP19, RASGRF2, CTNNAL1, JAK3, NCAM1, SLC4A11, AGAP3, CBLB, MAP2K6, CNKSR2, PEX5L, CDH13, ARHGEF10. ANK2, MARVELD3, ASPH, CADM1, DAAM2, GRIN2B, BCR, SLIT3, PKD1, MZB1, ROR2, SEMA6A, CCDC13, PFKL, CHST11, CHL1, TRPV1, NOTCH2, GABRB3, GABRB1, RELN, TCF7, COL1A1, DLG5, FCHO1, HSPG2, NRG4, CERS1, EPHA6, DISC1, SYNGAP1, NRXN1, AKT3, TWIST2, MAP3K12, PROX1, DNER, BCL11A, RGL1, PIK3C2G, RORC, ZFHX3, RTN4R, PARD3, TNKS1BP1, CHD7, SPTBN1, PLXND1, CDT1, FOXL1, CCR10. VWCE, FBN2, DOCK5, FMOD, CTNNA2, FGD4, RRH, FGF22, SKI, CCDC88A, FNIP2, ATP13A2, MGLL, RAPGEFL1, PLCB2, WWC2, PDGFD, SCARA5, HTR1F, VIPR1, FRMPD1, LRRN2, AUTS2, GLRA1, LCK, CAMK1, EPHA7, SGCA, FZD3, CELSR1, PPP3CA, CACNA1G, BMPR1B, MTSS1, ADAM8, ERBB3, ADCY5, SH2D3C, CPLX2, MNAT1, PCNT, COL4A2, SCARF1, BICD1, D2HGDH, DST, EPHB2, REM1, TUFT1, EPHB1, ANGPT2, PRKAG2, ADAM12, LRCH1, NHEJ1, ZBTB40. ZNF385B, CHD2, SAMD14, TSPAN18, EFNB2, NKX2-2, ADORA2A, RBBP5, KCNIP2, GAL, PIK3C2B, CAMKMT, NSDHL, ABCG4, RAPGEF5, SLC9A9, STAB2, MLLT3 |
| GO:0042063 | gliogenesis | 0.047 | 0.047 | CDH2, ZMIZ1, TENM4, NTRK2, DAB1, DMD, CNTN1, ARHGEF10. DAAM2, ROR2, RELN, DISC1, DNER, PARD3, MXRA8, VAX1, SKI, ERBB3, NKX2-2, ADORA2A |
| GO:0097106 | postsynaptic density organization | 0.047 | 0.047 | GRID2, CDH2, CSMD2, RELN, SYNGAP1, NRXN1 |
| GO:0022603 | regulation of anatomical structure morphogenesis | 0.047 | 0.047 | WNT7A, DNM3, DCN, TENM4, IL1RAPL1, NTRK2, ROBO2, DAB1, WT1, GLUL, PKHD1, KIF1A, BCR, TANC2, SEMA6A, RELN, DBN1, HSPG2, DISC1, SYNGAP1, AKT3, ZDHHC15, BCL11A, RTN4R, PLXND1, FGD4, CDKL5, EPHA7, PPP3CA, COL4A2, EPHB2, MYH10. ANGPT2, ADAM12, MAP2, TSPAN18, EFNB2 |
| GO:0097119 | postsynaptic density protein 95 clustering | 0.048 | 0.048 | CDH2, RELN, NRXN1 |
| GO:0051056 | regulation of small GTPase mediated signal transduction | 0.050 | 0.050 | ARHGEF10L, SH2B2, CD2AP, ARHGAP24, ADCYAP1R1, PREX2, DGKI, ARHGAP19, RASGRF2, ARHGEF10. BCR, NOTCH2, RELN, SYNGAP1, RTN4R, FGD4, AUTS2, EPHB2 |
| GO:0043005 | neuron projection | <0.0001 | <0.0001 | PTPRF, CNTN5, GABBR2, GRID2, PLXDC1, CDH2, TENM1, DNM3, KCNIP4, USH2A, CHRM5, CLSTN2, TENM4, IL1RAPL1, CTTNBP2, CD2AP, NTRK2, SHISA6, ROBO2, MAP1S, DMD, MAGI2, KIFAP3, TP63, RAB3A, PTPRO, CNTN1, KIF1A, DGKI, CTNND2, CDH13, SLC38A8, ANK2, CADM1, GRIN2B, BCR, TANC2, ROR2, SEMA6A, DCTN2, CHL1, TRPV1, GABRB3, GABRB1, RELN, DBN1, SDCCAG8, EPHA6, DISC1, SYNGAP1, NRXN1, MAP3K12, DNER, ACAP3, ELOVL5, BCL11B, RTN4R, PARD3, SPTBN1, PLXND1, CTNNA2, RRH, ATP13A2, CDKL5, HTR1F, AUTS2, GLRA1, EPHA7, APBA1, FZD3, PPP3CA, BMPR1B, SH2D3C, BSN, CPLX2, DST, EPHB2, EPHB1, SACS, ATL1, GABRA2, MAP2, SAMD14, ADORA2A, KCNIP2 |
| GO:0098794 | postsynapse | <0.0001 | <0.0001 | GABBR2, GRID1, DLGAP1, GRID2, CDH2, ELMO1, DNM3, CHRM5, CLSTN2, IL1RAPL1, CTTNBP2, NTRK2, SHISA6, NBEA, CSMD2, DLGAP2, DMD, MAGI2, KIFAP3, RAB3A, CLMP, PTPRO, PRRT1, CNTN1, DGKI, CTNND2, PTPRT, AGAP3, CBLB, CNKSR2, ANK2, CADM1, GRIN2B, BCR, TANC2, ROR2, TRPV1, GABRB3, GABRB1, DLG5, DBN1, FCHO1, DISC1, SYNGAP1, ZDHHC15, BCL11A, SPTBN1, CDKL5, GLRA1, UTRN, APBA1, PPP3CA, BSN, CPLX2, EPHB2, MYH10. GABRA2, SAMD14, EFNB2, ADORA2A |
| GO:0030425 | dendrite | <0.0001 | <0.0001 | GRID2, PLXDC1, DNM3, KCNIP4, CHRM5, CLSTN2, IL1RAPL1, CTTNBP2, CD2AP, NTRK2, SHISA6, MAP1S, MAGI2, TP63, PTPRO, KIF1A, DGKI, CTNND2, CADM1, BCR, TANC2, ROR2, CHL1, TRPV1, GABRB3, GABRB1, RELN, DBN1, EPHA6, SYNGAP1, DNER, ELOVL5, RTN4R, CDKL5, HTR1F, GLRA1, EPHA7, APBA1, FZD3, PPP3CA, BMPR1B, BSN, CPLX2, EPHB2, EPHB1, SACS, GABRA2, MAP2, SAMD14, ADORA2A, KCNIP2 |
| GO:0097447 | dendritic tree | <0.0001 | <0.0001 | GRID2, PLXDC1, DNM3, KCNIP4, CHRM5, CLSTN2, IL1RAPL1, CTTNBP2, CD2AP, NTRK2, SHISA6, MAP1S, MAGI2, TP63, PTPRO, KIF1A, DGKI, CTNND2, CADM1, BCR, TANC2, ROR2, CHL1, TRPV1, GABRB3, GABRB1, RELN, DBN1, EPHA6, SYNGAP1, DNER, ELOVL5, RTN4R, CDKL5, HTR1F, GLRA1, EPHA7, APBA1, FZD3, PPP3CA, BMPR1B, BSN, CPLX2, EPHB2, EPHB1, SACS, GABRA2, MAP2, SAMD14, ADORA2A, KCNIP2 |
| GO:0042995 | cell projection | <0.0001 | <0.0001 | PTPRF, LRGUK, MYO6, CNTN5, GABBR2, GRID2, PLXDC1, CDH2, TENM1, DNM3, KCNIP4, SH2B2, USH2A, CHRM5, CLSTN2, TENM4, IL1RAPL1, CTTNBP2, CUBN, CD2AP, NTRK2, SHISA6, ROBO2, MAP1S, DMD, MAGI2, ARHGAP24, KIFAP3, TP63, TNS3, RAB3A, GLUL, PKHD1, PTPRO, CNTN1, KIF1A, IFT43, DGKI, CTNND2, CDH13, SLC38A8, ANK2, CADM1, GRIN2B, BCR, TANC2, PKD1, ROR2, SEMA6A, CCDC13, DCTN2, CHL1, TRPV1, NOTCH2, GABRB3, GABRB1, RELN, DLG5, DBN1, SDCCAG8, EPHA6, DISC1, SYNGAP1, NRXN1, MAP3K12, DNER, ACAP3, ELOVL5, BCL11B, RTN4R, PARD3, SPTBN1, PLXND1, MXRA8, DOCK5, CTNNA2, FGD4, RRH, CCDC88A, ATP13A2, RILP, CDKL5, CTNNA3, HTR1F, LDB3, AUTS2, GLRA1, UTRN, EPHA7, APBA1, FZD3, PPP3CA, BMPR1B, MTSS1, ADCY5, SH2D3C, BSN, CPLX2, DST, AKAP14, EPHB2, MYH10. EPHB1, ANGPT2, SACS, ATL1, ZMYND12, GABRA2, LRRC23, MAP2, SAMD14, ADORA2A, KCNIP2, SH3PXD2A |
| GO:0036477 | somatodendritic compartment | <0.0001 | <0.0001 | PTPRF, GRID2, PLXDC1, DNM3, KCNIP4, USH2A, CHRM5, CLSTN2, IL1RAPL1, CTTNBP2, CD2AP, NTRK2, SHISA6, CACNA1B, MAP1S, MAGI2, TP63, ASTN2, PTPRO, KIF1A, DGKI, CTNND2, CNKSR2, CADM1, BCR, TANC2, ROR2, CHL1, TRPV1, GABRB3, GABRB1, RELN, DBN1, EPHA6, SYNGAP1, NRXN1, DNER, ELOVL5, RTN4R, ATP13A2, CDKL5, HTR1F, GLRA1, EPHA7, APBA1, FZD3, PPP3CA, BMPR1B, BSN, CPLX2, EPHB2, EPHB1, SACS, GABRA2, MAP2, SAMD14, ADORA2A, KCNIP2, GAL |
| GO:0120025 | plasma membrane bounded cell projection | <0.0001 | <0.0001 | PTPRF, MYO6, CNTN5, GABBR2, GRID2, PLXDC1, CDH2, TENM1, DNM3, KCNIP4, SH2B2, USH2A, CHRM5, CLSTN2, TENM4, IL1RAPL1, CTTNBP2, CUBN, CD2AP, NTRK2, SHISA6, ROBO2, MAP1S, DMD, MAGI2, KIFAP3, TP63, RAB3A, GLUL, PKHD1, PTPRO, CNTN1, KIF1A, IFT43, DGKI, CTNND2, CDH13, SLC38A8, ANK2, CADM1, GRIN2B, BCR, TANC2, PKD1, ROR2, SEMA6A, DCTN2, CHL1, TRPV1, NOTCH2, GABRB3, GABRB1, RELN, DLG5, DBN1, SDCCAG8, EPHA6, DISC1, SYNGAP1, NRXN1, MAP3K12, DNER, ACAP3, ELOVL5, BCL11B, RTN4R, PARD3, SPTBN1, PLXND1, MXRA8, CTNNA2, FGD4, RRH, CCDC88A, ATP13A2, RILP, CDKL5, CTNNA3, HTR1F, LDB3, AUTS2, GLRA1, UTRN, EPHA7, APBA1, FZD3, PPP3CA, BMPR1B, MTSS1, ADCY5, SH2D3C, BSN, CPLX2, DST, AKAP14, EPHB2, MYH10. EPHB1, SACS, ATL1, ZMYND12, GABRA2, LRRC23, MAP2, SAMD14, ADORA2A, KCNIP2 |
| GO:0098978 | glutamatergic synapse | <0.0001 | <0.0001 | GRID1, DLGAP1, GRID2, WNT7A, ELMO1, DNM3, CLSTN2, TENM4, IL1RAPL1, CTTNBP2, SHISA6, NBEA, CSMD2, DLGAP2, DAB1, KIFAP3, PTPRO, PRRT1, CNTN1, DGKI, PTPRT, RASGRF2, AGAP3, CBLB, CNKSR2, BCR, ROR2, DLG5, DBN1, DISC1, SYNGAP1, NRXN1, RTN4R, SPTBN1, PLXND1, CDKL5, EPHA7, APBA1, FZD3, PPP3CA, BSN, CPLX2, EPHB2, EPHB1, EFNB2, ADORA2A |
| GO:0030054 | cell junction | <0.0001 | <0.0001 | CDH18, CNTN5, GABBR2, GRID1, DLGAP1, GRID2, PLXDC1, CDH2, WNT7A, ELMO1, DNM3, PARVG, PANX3, PTPRN2, USH2A, CHRM5, CLSTN2, TENM4, IL1RAPL1, CTTNBP2, CD2AP, NTRK2, SHISA6, NBEA, CSMD2, CACNA1B, DLGAP2, MAP1S, CDH12, SDK1, RPS6KA2, DAB1, DMD, MAGI2, ARHGAP24, KIFAP3, TNS3, ADCYAP1R1, RAB3A, CLMP, PTPRO, PRRT1, CNTN1, KIF1A, DGKI, CTNND2, PTPRT, RASGRF2, AGAP3, CBLB, CNKSR2, CDH13, ANK2, MARVELD3, LAMA4, CADM1, GRIN2B, BCR, TANC2, ROR2, TRPV1, GABRB3, GABRB1, CACNB2, DLG5, DBN1, FCHO1, SDCCAG8, HSPG2, DISC1, SYNGAP1, NRXN1, ZDHHC15, BCL11A, RTN4R, PARD3, TNKS1BP1, SPTBN1, PLXND1, MXRA8, DOCK5, CTNNA2, CDKL5, CTNNA3, HTR1F, LDB3, GLRA1, UTRN, EPHA7, APBA1, SGCA, KAZN, FZD3, PPP3CA, CACNA1G, BSN, CPLX2, DST, EPHB2, MYH10. FRMD4A, EPHB1, MRC2, GABRA2, SAMD14, EFNB2, ADORA2A, KCNIP2, SH3PXD2A, TNS1 |
| GO:0099572 | postsynaptic specialization | <0.0001 | <0.0001 | GRID1, DLGAP1, GRID2, CDH2, DNM3, CLSTN2, NTRK2, SHISA6, CSMD2, DLGAP2, DMD, MAGI2, PTPRO, PRRT1, DGKI, CTNND2, PTPRT, AGAP3, CNKSR2, CADM1, GRIN2B, BCR, GABRB3, GABRB1, DLG5, DBN1, DISC1, SYNGAP1, ZDHHC15, SPTBN1, CDKL5, BSN, GABRA2, SAMD14, EFNB2 |
| GO:0045211 | postsynaptic membrane | <0.0001 | <0.0001 | GABBR2, GRID1, GRID2, CDH2, DNM3, CHRM5, CLSTN2, IL1RAPL1, SHISA6, NBEA, CSMD2, DMD, CLMP, PTPRO, PRRT1, CNTN1, PTPRT, CNKSR2, ANK2, GRIN2B, TRPV1, GABRB3, GABRB1, DBN1, GLRA1, UTRN, EPHB2, GABRA2, EFNB2, ADORA2A |
| GO:0045202 | synapse | <0.0001 | <0.0001 | CNTN5, GABBR2, GRID1, DLGAP1, GRID2, CDH2, WNT7A, ELMO1, DNM3, PTPRN2, USH2A, CHRM5, CLSTN2, TENM4, IL1RAPL1, CTTNBP2, CD2AP, NTRK2, SHISA6, NBEA, CSMD2, CACNA1B, DLGAP2, MAP1S, SDK1, RPS6KA2, DAB1, DMD, MAGI2, KIFAP3, RAB3A, CLMP, PTPRO, PRRT1, CNTN1, KIF1A, DGKI, CTNND2, PTPRT, RASGRF2, AGAP3, CBLB, CNKSR2, CDH13, ANK2, LAMA4, CADM1, GRIN2B, BCR, TANC2, ROR2, TRPV1, GABRB3, GABRB1, CACNB2, DLG5, DBN1, FCHO1, DISC1, SYNGAP1, NRXN1, ZDHHC15, BCL11A, RTN4R, SPTBN1, PLXND1, CDKL5, HTR1F, GLRA1, UTRN, EPHA7, APBA1, FZD3, PPP3CA, CACNA1G, BSN, CPLX2, EPHB2, MYH10. EPHB1, GABRA2, SAMD14, EFNB2, ADORA2A, KCNIP2 |
| GO:0071944 | cell periphery | <0.0001 | <0.0001 | CDH18, PTPRF, MYO6, CNTN5, GABBR2, GRID1, PCDH7, KCNK17, DLGAP1, GRID2, PLXDC1, CDH2, MDGA2, WNT7A, TENM1, ELMO1, DNM3, PARVG, PANX3, KCNIP4, SH2B2, PTPRN2, USH2A, CHRM5, CLSTN2, FCER1G, DCN, TENM4, IL1RAPL1, CTTNBP2, CUBN, CD2AP, NTRK2, SHISA6, NBEA, CSMD2, DPP10. NLRX1, CACNA1B, SLC4A3, ROBO2, DLGAP2, SLCO3A1, CDH12, NKAIN2, SDK1, PIK3R5, DMD, CPE, MAGI2, ADAMTS2, A2M, ABI3BP, EDIL3, FLT3, KIFAP3, KCNQ5, ADCYAP1R1, SPON1, RAB3A, PREX2, PTPRG, GLUL, CLMP, PKHD1, PLCD1, ASTN2, PTPRO, CRIM1, PRRT1, CNTN1, LTBP1, DGKI, CTNND2, PTPRT, ARHGAP19, RASGRF2, CTNNAL1, JAK3, NCAM1, SLC4A11, AGAP3, CBLB, CNKSR2, MUC6, CDH13, SLC38A8, ANK2, ASPH, LAMA4, CADM1, GRIN2B, BCR, EMCN, COL22A1, PKD1, NID2, PODXL2, ROR2, SEMA6A, CHL1, TRPV1, NOTCH2, GABRB3, GABRB1, RELN, TRPM3, CACNB2, COL1A1, DLG5, DBN1, FCHO1, GAL3ST1, HSPG2, NRG4, EPHA6, SYNGAP1, NRXN1, MAP3K12, LRP1B, DNER, RGL1, CPAMD8, PIK3C2G, RTN4R, PARD3, TNKS1BP1, SPTBN1, PLXND1, CCR10. MXRA8, FBN2, DOCK5, FMOD, CTNNA2, RRH, CCDC88A, MGLL, RAPGEFL1, CDKL5, SCARA5, HTR1F, VIPR1, FRMPD1, LRRN2, GLRA1, LCK, UTRN, EPHA7, APBA1, SGCA, KAZN, FZD3, CELSR1, PPP3CA, CACNA1G, BMPR1B, MTSS1, ADAM8, ERBB3, ADCY5, SH2D3C, BSN, COL4A2, SCARF1, ANTXRL, DST, EPHB2, REM1, MYH10. EPHB1, ANGPT2, ADAM12, GABRA2, TSPAN18, EFNB2, ADORA2A, NBEAL2, KCNIP2, PIK3C2B, ABCG4, RAPGEF5, SLC9A9, STAB2 |
| GO:0032279 | asymmetric synapse | <0.0001 | <0.0001 | GRID1, DLGAP1, GRID2, CDH2, DNM3, CLSTN2, NTRK2, SHISA6, CSMD2, DLGAP2, MAGI2, PTPRO, PRRT1, DGKI, CTNND2, PTPRT, AGAP3, CNKSR2, CADM1, GRIN2B, BCR, DLG5, DBN1, DISC1, SYNGAP1, ZDHHC15, SPTBN1, CDKL5, BSN, SAMD14, EFNB2, ADORA2A |
| GO:0097060 | synaptic membrane | <0.0001 | <0.0001 | CNTN5, GABBR2, GRID1, GRID2, CDH2, DNM3, CHRM5, CLSTN2, IL1RAPL1, SHISA6, NBEA, CSMD2, DMD, CLMP, PTPRO, PRRT1, CNTN1, DGKI, PTPRT, CNKSR2, ANK2, GRIN2B, TRPV1, GABRB3, GABRB1, DBN1, NRXN1, GLRA1, UTRN, APBA1, FZD3, EPHB2, GABRA2, EFNB2, ADORA2A |
| GO:0014069 | postsynaptic density | <0.0001 | <0.0001 | GRID1, DLGAP1, GRID2, CDH2, DNM3, CLSTN2, NTRK2, SHISA6, CSMD2, DLGAP2, MAGI2, PTPRO, PRRT1, DGKI, CTNND2, PTPRT, AGAP3, CNKSR2, CADM1, GRIN2B, BCR, DLG5, DBN1, DISC1, SYNGAP1, ZDHHC15, SPTBN1, CDKL5, BSN, SAMD14, EFNB2 |
| GO:0098984 | neuron to neuron synapse | <0.0001 | <0.0001 | GRID1, DLGAP1, GRID2, CDH2, DNM3, CLSTN2, NTRK2, SHISA6, CSMD2, DLGAP2, MAGI2, PTPRO, PRRT1, DGKI, CTNND2, PTPRT, AGAP3, CNKSR2, CADM1, GRIN2B, BCR, DLG5, DBN1, DISC1, SYNGAP1, ZDHHC15, SPTBN1, CDKL5, BSN, EPHB2, SAMD14, EFNB2, ADORA2A |
| GO:0030424 | axon | <0.0001 | <0.0001 | CNTN5, DNM3, USH2A, IL1RAPL1, CD2AP, NTRK2, ROBO2, RAB3A, PTPRO, CNTN1, KIF1A, DGKI, SLC38A8, BCR, TANC2, SEMA6A, DCTN2, DBN1, DISC1, NRXN1, MAP3K12, ACAP3, RTN4R, PARD3, SPTBN1, PLXND1, CTNNA2, CDKL5, AUTS2, FZD3, SH2D3C, BSN, CPLX2, DST, EPHB2, EPHB1, SACS, ATL1, GABRA2, MAP2, ADORA2A |
| GO:0005886 | plasma membrane | <0.0001 | <0.0001 | CDH18, PTPRF, MYO6, CNTN5, GABBR2, GRID1, PCDH7, KCNK17, DLGAP1, GRID2, PLXDC1, CDH2, MDGA2, WNT7A, TENM1, ELMO1, DNM3, PARVG, PANX3, KCNIP4, SH2B2, PTPRN2, USH2A, CHRM5, CLSTN2, FCER1G, TENM4, IL1RAPL1, CUBN, CD2AP, NTRK2, SHISA6, NBEA, CSMD2, DPP10. NLRX1, CACNA1B, SLC4A3, ROBO2, DLGAP2, SLCO3A1, CDH12, NKAIN2, SDK1, PIK3R5, DMD, CPE, MAGI2, FLT3, KIFAP3, KCNQ5, ADCYAP1R1, RAB3A, PREX2, PTPRG, GLUL, CLMP, PKHD1, PLCD1, PTPRO, CRIM1, PRRT1, CNTN1, DGKI, CTNND2, PTPRT, ARHGAP19, RASGRF2, CTNNAL1, JAK3, NCAM1, SLC4A11, CBLB, CNKSR2, MUC6, CDH13, ANK2, ASPH, CADM1, GRIN2B, BCR, EMCN, PKD1, NID2, PODXL2, ROR2, SEMA6A, CHL1, TRPV1, NOTCH2, GABRB3, GABRB1, RELN, TRPM3, CACNB2, DLG5, DBN1, FCHO1, GAL3ST1, HSPG2, NRG4, EPHA6, SYNGAP1, NRXN1, MAP3K12, LRP1B, DNER, RGL1, CPAMD8, PIK3C2G, RTN4R, PARD3, TNKS1BP1, SPTBN1, PLXND1, CCR10. MXRA8, DOCK5, CTNNA2, RRH, CCDC88A, MGLL, RAPGEFL1, CDKL5, SCARA5, HTR1F, VIPR1, FRMPD1, GLRA1, LCK, UTRN, EPHA7, APBA1, SGCA, KAZN, FZD3, CELSR1, PPP3CA, CACNA1G, BMPR1B, MTSS1, ADAM8, ERBB3, ADCY5, SH2D3C, SCARF1, ANTXRL, DST, EPHB2, REM1, MYH10. EPHB1, ADAM12, GABRA2, TSPAN18, EFNB2, ADORA2A, NBEAL2, KCNIP2, PIK3C2B, ABCG4, RAPGEF5, SLC9A9, STAB2 |
| GO:0099634 | postsynaptic specialization membrane | 0.001 | 0.001 | GRID1, GRID2, CDH2, CLSTN2, SHISA6, CSMD2, PTPRO, PRRT1, PTPRT, CNKSR2, GRIN2B, GABRB3, GABRB1, GABRA2, EFNB2 |
| GO:0098590 | plasma membrane region | 0.002 | 0.002 | MYO6, CNTN5, GABBR2, GRID1, GRID2, CDH2, DNM3, USH2A, CHRM5, CLSTN2, IL1RAPL1, CUBN, SHISA6, NBEA, CSMD2, ROBO2, SLCO3A1, DMD, KIFAP3, ADCYAP1R1, CLMP, PKHD1, PTPRO, PRRT1, CNTN1, DGKI, PTPRT, SLC4A11, CNKSR2, CDH13, ANK2, CADM1, GRIN2B, PKD1, TRPV1, GABRB3, GABRB1, DBN1, NRXN1, PARD3, SPTBN1, PLXND1, MXRA8, CDKL5, GLRA1, UTRN, APBA1, FZD3, ERBB3, SH2D3C, DST, EPHB2, MYH10. GABRA2, EFNB2, ADORA2A |
| GO:0098982 | GABA-ergic synapse | 0.002 | 0.002 | CNTN5, GRID1, NBEA, DMD, PTPRO, CDH13, TRPV1, GABRB1, DISC1, NRXN1, BSN, GABRA2 |
| GO:0019897 | extrinsic component of plasma membrane | 0.003 | 0.003 | CDH18, CDH2, CUBN, NBEA, CDH12, JAK3, CNKSR2, CDH13, CTNNA2, LCK, PPP3CA |
| GO:0005911 | cell-cell junction | 0.003 | 0.003 | CDH18, PLXDC1, CDH2, PANX3, CD2AP, CDH12, MAGI2, ARHGAP24, ADCYAP1R1, CLMP, CTNND2, CDH13, ANK2, MARVELD3, CADM1, DLG5, DBN1, SDCCAG8, PARD3, TNKS1BP1, MXRA8, CTNNA2, CTNNA3, LDB3, SGCA, KAZN, PPP3CA, FRMD4A, EFNB2 |
| GO:0009986 | cell surface | 0.006 | 0.006 | CDH2, WNT7A, CLSTN2, FCER1G, IL1RAPL1, CUBN, SLC4A3, ROBO2, DMD, ADCYAP1R1, CLMP, PKHD1, PTPRT, NCAM1, CDH13, GRIN2B, EMCN, PKD1, ROR2, TRPV1, NOTCH2, NRXN1, RTN4R, CCR10. MXRA8, SCARA5, GLRA1, FZD3, ADAM8, BSN, ANTXRL, EPHB2, TNS1, STAB2 |
| GO:0019898 | extrinsic component of membrane | 0.008 | 0.008 | CDH18, CDH2, CUBN, NBEA, CDH12, PIK3R5, JAK3, CNKSR2, CDH13, PIK3C2G, CTNNA2, LCK, PPP3CA, PIK3C2B |
| GO:0098839 | postsynaptic density membrane | 0.011 | 0.011 | GRID1, GRID2, CLSTN2, SHISA6, CSMD2, PTPRO, PRRT1, PTPRT, CNKSR2, GRIN2B, EFNB2 |
| GO:0043197 | dendritic spine | 0.011 | 0.011 | GRID2, DNM3, CTTNBP2, NTRK2, SHISA6, PTPRO, DGKI, BCR, TANC2, TRPV1, GABRB3, APBA1, PPP3CA, EPHB2 |
| GO:0044309 | neuron spine | 0.013 | 0.013 | GRID2, DNM3, CTTNBP2, NTRK2, SHISA6, PTPRO, DGKI, BCR, TANC2, TRPV1, GABRB3, APBA1, PPP3CA, EPHB2 |
| GO:0005856 | cytoskeleton | 0.014 | 0.014 | LRGUK, MYO6, CDH2, NAV3, TENM1, DNM3, PARVG, SH2B2, USH2A, CTTNBP2, INCENP, CD2AP, DLGAP2, MAP1S, SMTNL2, PIK3R5, DMD, MAGI2, ARHGAP24, KIFAP3, TNS3, CLMP, PKHD1, CCNB3, KIF1A, MAP7D3, IFT43, CTNNAL1, JAK3, MAP2K6, KATNAL1, KIF18B, ARHGEF10. ANK2, LRRCC1, GRIN2B, ROR2, CCDC13, DCTN2, DLG5, DBN1, SDCCAG8, DISC1, PARD3, CEP85L, TNKS1BP1, SPTBN1, KIF13A, PKNOX2, DOCK5, CTNNA2, FGD4, SKI, CCDC88A, FNIP2, FAM110B, RILP, CDKL5, CTNNA3, LDB3, FRMPD1, AUTS2, LCK, UTRN, NEBL, SGCA, KAZN, TNNC2, MTSS1, ADAM8, BSN, PCNT, CCDC6, BICD1, DST, AKAP14, MYH10. FRMD4A, LRRC23, MAP2, SAMD14, ADORA2A, SH3PXD2A, TNS1 |
| GO:0005912 | adherens junction | 0.019 | 0.019 | CDH18, CDH2, CDH12, ARHGAP24, CTNND2, CDH13, DLG5, PARD3, TNKS1BP1, CTNNA2, CTNNA3, LDB3, FRMD4A, EFNB2 |
| GO:0044297 | cell body | 0.019 | 0.019 | PTPRF, PLXDC1, KCNIP4, USH2A, CACNA1B, MAP1S, GLUL, ASTN2, CTNND2, CNKSR2, ROR2, TRPV1, DISC1, NRXN1, DNER, ELOVL5, RTN4R, PLXND1, ATP13A2, GLRA1, FZD3, BMPR1B, CPLX2, EPHB2, GABRA2, MAP2, ADORA2A, GAL |
| GO:0036057 | slit diaphragm | 0.019 | 0.019 | CD2AP, MAGI2, PPP3CA |
| GO:0036056 | filtration diaphragm | 0.019 | 0.019 | CD2AP, MAGI2, PPP3CA |
| GO:0016342 | catenin complex | 0.020 | 0.020 | CDH18, CDH2, CDH12, CDH13, CTNNA2 |
| GO:0043235 | receptor complex | 0.020 | 0.020 | GABBR2, GRID2, PLXDC1, PTPRN2, CUBN, NTRK2, SHISA6, FLT3, ADCYAP1R1, PEX5L, GRIN2B, ROR2, NOTCH2, GABRB3, GABRB1, LRP1B, PLXND1, VIPR1, BMPR1B, ERBB3, GABRA2 |
| GO:0030312 | external encapsulating structure | 0.020 | 0.020 | CDH2, WNT7A, USH2A, DCN, ADAMTS2, A2M, ABI3BP, EDIL3, SPON1, LTBP1, NCAM1, MUC6, CDH13, LAMA4, COL22A1, NID2, RELN, COL1A1, HSPG2, FBN2, FMOD, LRRN2, COL4A2, DST, ANGPT2 |
| GO:0031012 | extracellular matrix | 0.020 | 0.020 | CDH2, WNT7A, USH2A, DCN, ADAMTS2, A2M, ABI3BP, EDIL3, SPON1, LTBP1, NCAM1, MUC6, CDH13, LAMA4, COL22A1, NID2, RELN, COL1A1, HSPG2, FBN2, FMOD, LRRN2, COL4A2, DST, ANGPT2 |
| GO:0043025 | neuronal cell body | 0.024 | 0.024 | PTPRF, PLXDC1, KCNIP4, USH2A, CACNA1B, MAP1S, ASTN2, CTNND2, CNKSR2, ROR2, TRPV1, NRXN1, DNER, ELOVL5, RTN4R, ATP13A2, GLRA1, FZD3, BMPR1B, CPLX2, EPHB2, GABRA2, MAP2, ADORA2A, GAL |
| GO:0032589 | neuron projection membrane | 0.026 | 0.026 | USH2A, SHISA6, ROBO2, TRPV1, SPTBN1, GABRA2, ADORA2A |
| GO:1902710 | GABA receptor complex | 0.026 | 0.026 | GABBR2, GABRB3, GABRB1, GABRA2 |
| GO:0062023 | collagen-containing extracellular matrix | 0.030 | 0.030 | CDH2, DCN, ADAMTS2, A2M, ABI3BP, EDIL3, SPON1, LTBP1, NCAM1, CDH13, LAMA4, COL22A1, NID2, COL1A1, HSPG2, FBN2, FMOD, COL4A2, ANGPT2 |
| GO:0098685 | Schaffer collateral - CA1 synapse | 0.037 | 0.037 | WNT7A, DGKI, BCR, GABRB1, NRXN1, APBA1, PPP3CA, BSN, EFNB2 |
| GO:0030426 | growth cone | 0.040 | 0.040 | CD2AP, PTPRO, DCTN2, DBN1, DISC1, NRXN1, MAP3K12, ACAP3, RTN4R, CDKL5, AUTS2, MAP2 |
| GO:0048786 | presynaptic active zone | 0.048 | 0.048 | CDH2, RAB3A, GABRB1, NRXN1, APBA1, FZD3, BSN, ADORA2A |
| GO:0030427 | site of polarized growth | 0.048 | 0.048 | CD2AP, PTPRO, DCTN2, DBN1, DISC1, NRXN1, MAP3K12, ACAP3, RTN4R, CDKL5, AUTS2, MAP2 |
| GO:0004888 | transmembrane signaling receptor activity | <0.0001 | <0.0001 | PTPRF, GABBR2, GRID1, GRID2, PTPRN2, CHRM5, FCER1G, NTRK2, ROBO2, FLT3, ADCYAP1R1, PTPRG, PTPRO, CRIM1, LTBP1, PTPRT, GRIN2B, PKD1, ROR2, TRPV1, GABRB3, GABRB1, EPHA6, NRXN1, DNER, RTN4R, PLXND1, CCR10. RRH, HTR1F, VIPR1, GLRA1, EPHA7, FZD3, CELSR1, BMPR1B, ERBB3, SCARF1, ANTXRL, EPHB2, EPHB1, GABRA2, ADORA2A, GAL |
| GO:0060089 | molecular transducer activity | <0.0001 | <0.0001 | PTPRF, GABBR2, GRID1, GRID2, FGF14, WNT7A, PTPRN2, CHRM5, FCER1G, CUBN, NTRK2, ESRRG, ROBO2, DHX16, FLT3, ADCYAP1R1, PTPRG, PKHD1, PTPRO, CRIM1, PRRT1, LTBP1, PTPRT, GRIN2B, PKD1, ROR2, SEMA6A, TRPV1, NOTCH2, GABRB3, GABRB1, RELN, NRG4, EPHA6, NRXN1, DNER, RORC, RTN4R, PLXND1, CCR10. FBN2, RRH, FGF22, PDGFD, HTR1F, VIPR1, LRRN2, GLRA1, EPHA7, FZD3, CELSR1, BMPR1B, ERBB3, SCARF1, ANTXRL, EPHB2, EPHB1, ANGPT2, MRC2, GABRA2, ADORA2A, GAL |
| GO:0038023 | signaling receptor activity | <0.0001 | <0.0001 | PTPRF, GABBR2, GRID1, GRID2, FGF14, WNT7A, PTPRN2, CHRM5, FCER1G, CUBN, NTRK2, ESRRG, ROBO2, DHX16, FLT3, ADCYAP1R1, PTPRG, PKHD1, PTPRO, CRIM1, PRRT1, LTBP1, PTPRT, GRIN2B, PKD1, ROR2, SEMA6A, TRPV1, NOTCH2, GABRB3, GABRB1, RELN, NRG4, EPHA6, NRXN1, DNER, RORC, RTN4R, PLXND1, CCR10. FBN2, RRH, FGF22, PDGFD, HTR1F, VIPR1, LRRN2, GLRA1, EPHA7, FZD3, CELSR1, BMPR1B, ERBB3, SCARF1, ANTXRL, EPHB2, EPHB1, ANGPT2, MRC2, GABRA2, ADORA2A, GAL |
| GO:0005509 | calcium ion binding | 0.002 | 0.002 | CDH18, PCDH7, CDH2, KCNIP4, CLSTN2, GPD2, CUBN, CACNA1B, CDH12, EDIL3, PLCD1, ASTN2, LTBP1, CBLB, CDH13, ASPH, SLIT3, NID2, NOTCH2, HSPG2, NRXN1, LRP1B, DNER, VWCE, FBN2, PLCB2, SGCA, CELSR1, TNNC2, PPP3CA, ADAM8, DST, KCNIP2, STAB2 |
| GO:0005201 | extracellular matrix structural constituent | 0.004 | 0.004 | DCN, ABI3BP, EDIL3, SPON1, LTBP1, MUC6, LAMA4, COL22A1, NID2, COL1A1, HSPG2, FBN2, FMOD, COL4A2, TUFT1 |
| GO:0019199 | transmembrane receptor protein kinase activity | 0.012 | 0.012 | NTRK2, FLT3, CRIM1, LTBP1, ROR2, EPHA6, EPHA7, BMPR1B, ERBB3, EPHB2, EPHB1 |
| GO:0005001 | transmembrane receptor protein tyrosine phosphatase activity | 0.012 | 0.012 | PTPRF, PTPRN2, PTPRG, PTPRO, PTPRT |
| GO:0019198 | transmembrane receptor protein phosphatase activity | 0.012 | 0.012 | PTPRF, PTPRN2, PTPRG, PTPRO, PTPRT |
| GO:0008013 | beta-catenin binding | 0.014 | 0.014 | CDH18, CDH2, CDH12, TCF7L2, CTNND2, PTPRT, CDH13, TCF7, DLG5, CTNNA2, CTNNA3 |
| GO:0008046 | axon guidance receptor activity | 0.017 | 0.017 | ROBO2, EPHA7, EPHB2, EPHB1 |
| GO:0003779 | actin binding | 0.026 | 0.026 | MYO6, WIPF3, PARVG, CD2AP, MAP1S, DMD, CTNNAL1, DAAM2, CACNB2, DBN1, SPTBN1, PKNOX2, CTNNA2, FGD4, CCDC88A, CTNNA3, LDB3, UTRN, NEBL, TNNC2, MTSS1, DST, MYH10. SAMD14, TNS1 |
| GO:0005230 | extracellular ligand-gated monoatomic ion channel activity | 0.028 | 0.028 | GRID1, GRID2, GRIN2B, TRPV1, GABRB3, GABRB1, GLRA1, GABRA2 |
| GO:0004714 | transmembrane receptor protein tyrosine kinase activity | 0.029 | 0.029 | NTRK2, FLT3, CRIM1, ROR2, EPHA6, EPHA7, ERBB3, EPHB2, EPHB1 |
| GO:0005005 | transmembrane-ephrin receptor activity | 0.029 | 0.029 | EPHA6, EPHA7, EPHB2, EPHB1 |
| GO:0030021 | extracellular matrix structural constituent conferring compression resistance | 0.033 | 0.033 | DCN, HSPG2, FMOD, TUFT1 |
| GO:0099507 | ligand-gated monoatomic ion channel activity involved in regulation of presynaptic membrane potential | 0.033 | 0.033 | GRIN2B, GABRB1, GLRA1, GABRA2 |
| GO:0008092 | cytoskeletal protein binding | 0.041 | 0.041 | MYO6, WIPF3, NAV3, DNM3, PARVG, USH2A, CD2AP, MAP1S, DMD, KIFAP3, RAB3A, KIF1A, MAP7D3, PTPRT, CTNNAL1, KATNAL1, KIF18B, ARHGEF10. ANK2, DAAM2, DCTN2, CACNB2, DLG5, DBN1, DISC1, SPTBN1, KIF13A, PKNOX2, CTNNA2, FGD4, CCDC88A, CTNNA3, LDB3, UTRN, NEBL, TNNC2, MTSS1, BICD1, DST, MYH10. MAP2, SAMD14, ADORA2A, TNS1 |
| GO:0022824 | transmitter-gated monoatomic ion channel activity | 0.041 | 0.041 | GRID1, GRID2, GRIN2B, GABRB3, GABRB1, GLRA1, GABRA2 |
| GO:0022835 | transmitter-gated channel activity | 0.041 | 0.041 | GRID1, GRID2, GRIN2B, GABRB3, GABRB1, GLRA1, GABRA2 |
| GO:0043167 | ion binding | 0.041 | 0.041 | CDH18, LRGUK, FOXP2, MYO6, NUDT14, PCDH7, ARHGEF10L, CDH2, PRDM16, CAMK1D, B4GALT4, NAV3, KLF12, ELMO1, DNM3, ZMIZ1, KCNIP4, HLCS, ZNF608, CLSTN2, ULK4, CUTA, GPD2, CUBN, CHTF18, RAD51B, NTRK2, ESRRG, NLRX1, CACNA1B, CREB5, ADARB2, CDH12, RNF217, TDRD9, DHX16, ZNF362, ZFP36L1, RPS6KA2, MBNL3, DMD, CPE, MCM8, ADAMTS2, PDE10A, EDIL3, FLT3, TP63, TNS3, SPON1, WT1, RAB3A, PREX2, GLUL, SHMT1, PLCD1, MTHFS, IP6K3, ASTN2, KIF1A, LTBP1, DGKI, RASGRF2, JAK3, AGAP3, CBLB, MAP2K6, KATNAL1, EBF1, KIF18B, CDH13, ARHGEF10. ASPH, GRIN2B, BCR, SLIT3, PCCA, NID2, ROR2, PFKL, TRPV1, NOTCH2, RELN, B4GALT3, COL1A1, ZNF521, YPEL4, HSPG2, EPHA6, NRXN1, AKT3, MAP3K12, LRP1B, DNER, ZDHHC15, BCL11A, RGL1, PIK3C2G, ACAP3, RORC, ZFHX3, BCL11B, RTN4R, PARD3, CHD7, KIF13A, VWCE, FBN2, DOCK5, FGD4, SIL1, SKI, CCDC88A, ATP13A2, TRPS1, MTA3, MCCC2, RAPGEFL1, ZNF618, PLCB2, CDKL5, HTR1F, LDB3, IDH2, GLRA1, LCK, UTRN, CAMK1, EPHA7, SGCA, CELSR1, TNNC2, PPP3CA, DDX52, BMPR1B, ADAM8, ERBB3, ADCY5, SH2D3C, BSN, MNAT1, D2HGDH, ANTXRL, DST, EPHB2, REM1, MYH10. EPHB1, AMPD1, ANGPT2, GALNT18, PRKAG2, ADAM12, ATL1, ZMYND12, RIOK1, ZBTB40. ZNF385B, CHD2, UBE2Z, KCNIP2, ZFPM2, ZFHX4, SH3PXD2A, EXD3, PIK3C2B, ABCG4, TNS1, RAPGEF5, STAB2 |
| GO:0005003 | ephrin receptor activity | 0.043 | 0.043 | EPHA6, EPHA7, EPHB2, EPHB1 |
| GO:0050682 | AF-2 domain binding | 0.043 | 0.043 | ESRRG, PPARGC1B |
| GO:0005198 | structural molecule activity | 0.047 | 0.047 | DNM3, PANX3, DCN, CD2AP, DMD, ABI3BP, EDIL3, SPON1, MRPL12, LTBP1, MUC6, ANK2, ASPH, LAMA4, COL22A1, NID2, COL1A1, HSPG2, SPTBN1, FBN2, FMOD, CTNNA2, NEBL, BSN, COL4A2, CCDC6, BICD1, DST, TUFT1, MAP2 |
| HP:0011097 | Epileptic spasm | 0.001 | 0.001 | GABBR2, CDH2, RUSC2, NTRK2, CACNA1B, DHX16, KCNQ5, GLUL, GRIN2B, GABRB3, TRPM3, SYNGAP1, AKT3, CEP85L, SPTBN1, CTNNA2, CDKL5, PPP3CA, D2HGDH, SLC25A10. CHD2 |
| HP:0002317 | Unsteady gait | 0.004 | 0.004 | GABBR2, RUSC2, NTRK2, CACNA1B, DHX16, DAB1, PDE10A, KCNQ5, KIF1A, CNKSR2, SYNGAP1, BCL11B, FGD4, SGCA, PPP3CA, CACNA1G, SACS, GABRA2, CHD2 |
| HP:0025373 | Interictal EEG abnormality | 0.004 | 0.004 | GABBR2, RUSC2, NTRK2, NBEA, CACNA1B, AMT, CNKSR2, GRIN2B, GABRB3, GABRB1, RELN, TRPM3, SYNGAP1, AKT3, PGAP2, SPTBN1, CTNNA2, CCDC88A, CDKL5, PPP3CA, CACNA1G, D2HGDH, SLC25A10. PRKAG2, GABRA2, CHD2 |
| HP:0011182 | Interictal epileptiform activity | 0.005 | 0.005 | GABBR2, RUSC2, NTRK2, NBEA, CACNA1B, AMT, CNKSR2, GRIN2B, GABRB3, GABRB1, RELN, SYNGAP1, AKT3, PGAP2, SPTBN1, CTNNA2, CCDC88A, CDKL5, PPP3CA, CACNA1G, D2HGDH, SLC25A10. PRKAG2, GABRA2, CHD2 |
| HP:0002353 | EEG abnormality | 0.005 | 0.005 | GABBR2, PRDM16, RUSC2, NTRK2, NBEA, CACNA1B, AMT, MTHFS, CTNND2, CNKSR2, GRIN2B, GABRB3, GABRB1, RELN, TRPM3, HSPG2, CERS1, SYNGAP1, AKT3, RTN4R, PGAP2, CEP85L, SPTBN1, CTNNA2, SKI, CCDC88A, CDKL5, PPP3CA, CACNA1G, D2HGDH, SLC25A10. PRKAG2, GABRA2, CHD2 |
| HP:0012469 | Infantile spasms | 0.005 | 0.005 | GABBR2, CDH2, RUSC2, NTRK2, DHX16, KCNQ5, GLUL, GRIN2B, GABRB3, TRPM3, CEP85L, SPTBN1, CTNNA2, CDKL5, D2HGDH, SLC25A10 |
| HP:0000006 | Autosomal dominant inheritance | 0.006 | 0.006 | FOXP2, MYO6, GABBR2, FGF14, CDH2, PRDM16, WNT7A, ZMIZ1, DCN, TENM4, GPD2, NTRK2, NBEA, TCOF1, SLC4A3, ROBO2, ERG, DHX16, TCF7L2, DAB1, PDE10A, FLT3, TP63, KCNQ5, WT1, GLUL, PLCD1, KIF1A, SLC4A11, ARHGEF10. ANK2, LAMA4, GRIN2B, TANC2, PKD1, ROR2, NOTCH2, GABRB3, GABRB1, RELN, TRPM3, CACNB2, COL1A1, SYNGAP1, AKT3, TWIST2, BCL11A, ZFHX3, ELOVL5, BCL11B, RTN4R, CEP85L, CHD7, SPTBN1, FOXL1, FBN2, SKI, TRPS1, CTNNA3, LDB3, IDH2, AUTS2, GLRA1, TBX5, CELSR1, TNNC2, PPP3CA, CACNA1G, BMPR1B, ERBB3, ADCY5, PBX1, COL4A2, ANGPT2, PRKAG2, ATL1, GABRA2, CHD2, GAL, ZFPM2, ZFHX4 |
| HP:0002527 | Falls | 0.006 | 0.006 | DAB1, DMD, GABRB3, CERS1, SPTBN1, ATP13A2, SACS, CHD2 |
| HP:0001311 | Abnormal nervous system electrophysiology | 0.008 | 0.008 | GABBR2, PRDM16, RUSC2, NTRK2, NBEA, CACNA1B, DHX16, AMT, MTHFS, KIF1A, CTNND2, CNKSR2, ARHGEF10. GRIN2B, GABRB3, GABRB1, RELN, TRPM3, HSPG2, CERS1, SYNGAP1, AKT3, RTN4R, PGAP2, CEP85L, SPTBN1, CTNNA2, FGD4, SKI, CCDC88A, CDKL5, PPP3CA, CACNA1G, D2HGDH, SLC25A10. PRKAG2, SACS, ATL1, GABRA2, CHD2 |
| HP:0200134 | Epileptic encephalopathy | 0.012 | 0.012 | GABBR2, NTRK2, CACNA1B, KCNQ5, GLUL, GRIN2B, GABRB3, GABRB1, SYNGAP1, NRXN1, CDKL5, PPP3CA, ALG9, CHD2 |
| HP:0011198 | EEG with generalized epileptiform discharges | 0.014 | 0.014 | GABBR2, RUSC2, NTRK2, CACNA1B, AMT, CNKSR2, GRIN2B, GABRB3, GABRB1, SYNGAP1, AKT3, PGAP2, SPTBN1, CTNNA2, CCDC88A, CDKL5, PPP3CA, D2HGDH, PRKAG2, GABRA2, CHD2 |
| HP:0030178 | Abnormality of central nervous system electrophysiology | 0.014 | 0.014 | GABBR2, PRDM16, RUSC2, NTRK2, NBEA, CACNA1B, AMT, MTHFS, CTNND2, CNKSR2, GRIN2B, GABRB3, GABRB1, RELN, TRPM3, HSPG2, CERS1, SYNGAP1, AKT3, RTN4R, PGAP2, CEP85L, SPTBN1, CTNNA2, SKI, CCDC88A, CDKL5, PPP3CA, CACNA1G, D2HGDH, SLC25A10. PRKAG2, SACS, GABRA2, CHD2 |
| HP:0100710 | Impulsivity | 0.017 | 0.017 | GABBR2, CDH2, IL1RAPL1, NTRK2, CACNA1B, AFF2, CNKSR2, GABRB3, RELN, SYNGAP1, SPTBN1, PPP3CA, SUGCT, GABRA2 |
| HP:0100660 | Dyskinesia | 0.023 | 0.023 | GABBR2, FGF14, NTRK2, CACNA1B, PDE10A, CNKSR2, GRIN2B, GABRB3, SYNGAP1, SIL1, ATP13A2, CDKL5, PPP3CA, ADCY5, SLC25A10. GABRA2 |
| HP:0002521 | Hypsarrhythmia | 0.023 | 0.023 | GABBR2, RUSC2, NTRK2, CACNA1B, CNKSR2, GRIN2B, GABRB3, GABRB1, SYNGAP1, SPTBN1, CTNNA2, CCDC88A, CDKL5, PPP3CA, D2HGDH, GABRA2 |
| HP:0100539 | Periorbital edema | 0.029 | 0.029 | CD2AP, MAGI2, ADAMTS2, ARHGAP24, WT1, PTPRO, DAAM2, COL1A1, SPTBN1, FKBP6, FRMD4A |
| HP:0031504 | Foamy urine | 0.032 | 0.032 | CD2AP, MAGI2, ARHGAP24, WT1, PTPRO, DAAM2 |
| HP:0011203 | EEG with abnormally slow frequencies | 0.039 | 0.039 | GABBR2, NTRK2, CACNA1B, MTHFS, CNKSR2, SYNGAP1, CDKL5, PPP3CA, GABRA2, CHD2 |
| HP:0006855 | Cerebellar vermis atrophy | 0.040 | 0.040 | GRID2, PIK3R5, DAB1, KIF1A, BCL11A, ELOVL5, CACNA1G, SACS |
| HP:0025783 | Diagnostic behavioral phenotype | 0.041 | 0.041 | FOXP2, GABBR2, CDH2, PRDM16, ZMIZ1, USH2A, IL1RAPL1, NTRK2, NBEA, CACNA1B, AFF2, CNKSR2, TANC2, GABRB3, TRPM3, SDCCAG8, HSPG2, SYNGAP1, BCL11A, BCL11B, RTN4R, PGAP2, CEP85L, CHD7, SPTBN1, PLXND1, FKBP6, CTNNA2, SKI, CDKL5, AUTS2, PPP3CA, GABRA2, CHD2, ZFPM2 |
| HP:0000729 | Autistic behavior | 0.041 | 0.041 | FOXP2, GABBR2, CDH2, PRDM16, ZMIZ1, IL1RAPL1, NTRK2, NBEA, CACNA1B, AFF2, CNKSR2, TANC2, GABRB3, TRPM3, SDCCAG8, HSPG2, SYNGAP1, BCL11A, BCL11B, PGAP2, CEP85L, CHD7, SPTBN1, PLXND1, FKBP6, CTNNA2, SKI, CDKL5, AUTS2, PPP3CA, GABRA2, CHD2, ZFPM2 |
| HP:0012579 | Minimal change glomerulonephritis | 0.047 | 0.047 | CD2AP, MAGI2, ARHGAP24, WT1, PTPRO, DAAM2 |
